# Resource Recovery from Wastewater By Directing Microbial Metabolism Toward Production of Value-added Biochemicals

**DOI:** 10.1101/2024.09.21.614227

**Authors:** Xueyang Zhou, Bharat Manna, Boyu Lyu, Gavin Lear, Joanne M. Kingsbury, Naresh Singhal

## Abstract

Transitioning wastewater treatment from mere pollutant removal to resource recovery necessitates exploiting the metabolic capabilities of microbial communities. Studies suggest that under low-oxygen conditions, microbes activate oxygen-responsive regulons that suppress the tricarboxylic acid (TCA) cycle, diverting carbon flux towards biosynthetic pathways and accumulating valuable organic metabolites. We hypothesized that dynamically altering dissolved oxygen levels in activated sludge would disrupt aerobic metabolic equilibrium, enhancing the production of valuable biochemicals like amino acids and fatty acids. To test this, batch experiments were conducted with activated sludge under constant aeration and rapid cycling between oxygen-rich and oxygen-poor states. Fluctuating oxygen concentrations between 0 and 2 mg/L significantly increased valuable biochemical production compared to constant aeration (*P*<0.05). Continuous oxygen perturbations increased free amino acids by 35.7±7.6% and free fatty acids by 76.4±13.0%, while intermittent perturbations with anoxic periods enhanced free amino acids by 42.4±8.1% and free fatty acids by 39.3±7.7%. Notably, 14 standard amino acids showed significant increases, and most fatty acids had carbon chain lengths between C12-C22. Mechanistically, compared to stable oxygen, oxygen perturbations activated the FNR and ArcA regulons, resulting in lower relative abundances of TCA cycle enzymes such as malate dehydrogenase, isocitrate dehydrogenase, and 2-oxoglutarate dehydrogenase, while higher relative abundances of amino acid (ilv cluster) and fatty acid (acc cluster) biosynthetic enzymes. Our findings demonstrate that introducing controlled oxygen fluctuations in wastewater treatment can enhance the biochemical value of activated sludge with minimal process modifications, facilitating resource recovery.

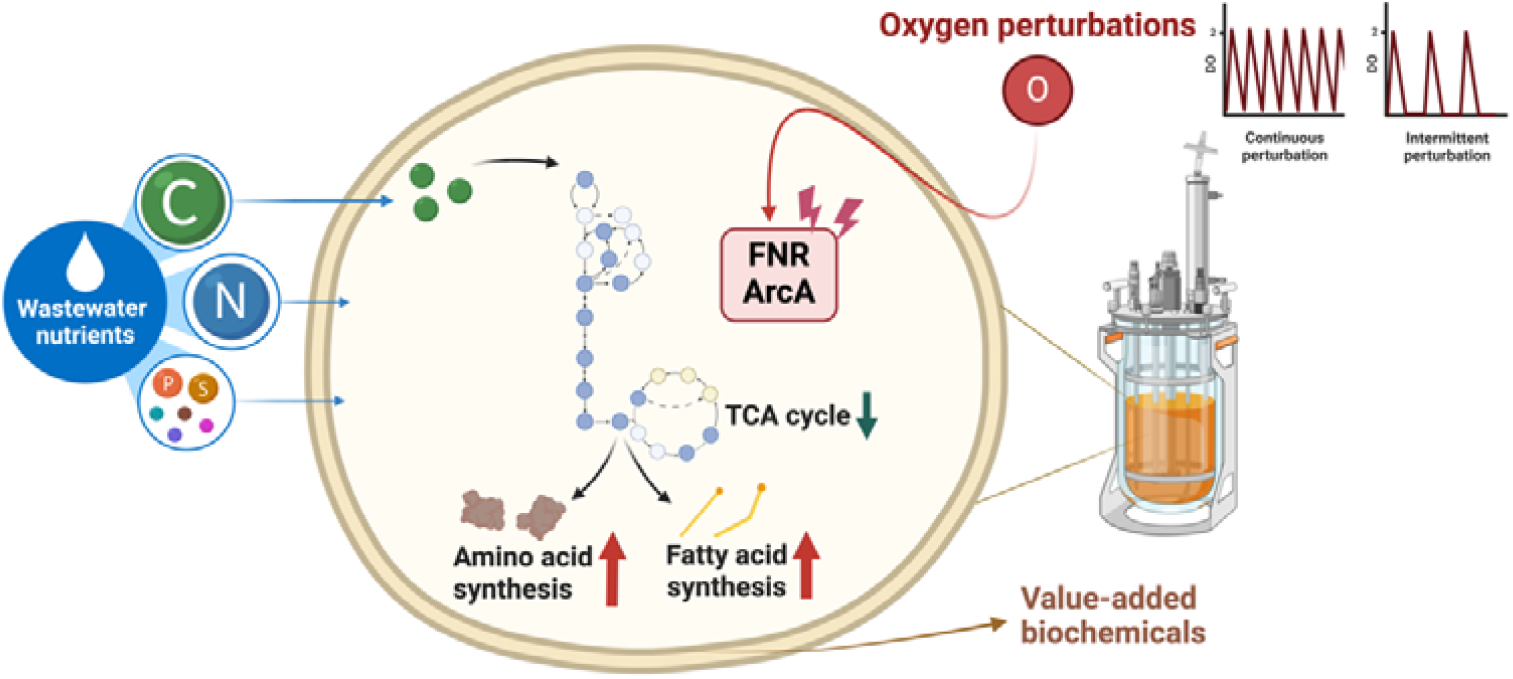

## 1. Introduction

The global surge in wastewater generation poses significant environmental and public health challenges, with billions of gallons produced annually. However, this challenge also presents an opportunity: bioresource recovery in wastewater treatment plants (WWTPs) offers a transformative solution by converting waste into valuable resources. The nutrients in wastewater, including carbon, nitrogen, and phosphorus, can be metabolized by microbes into biomass constituents such as proteins, fatty acids, and carbohydrates^1^. These biochemicals have extensive applications across various industries, from energy to biomaterial production^2^. For instance, energy recovery from sludge biomass in WWTPs can potentially save up to 5000 MWh annually^3^, while the global market for bioplastics is expected to reach USD 27.9 billion by 2025^4^.

Traditionally, wastewater treatment has focused on pollutant removal, with high dissolved oxygen (DO) levels being the norm. Maintaining DO levels above 2 mg/L has been shown to improve COD removal and ammonia oxidation rates significantly^5^. However, this approach oxidizes a large amount of carbon into carbon dioxide, which is not optimal for resource recovery. If activated sludge can convert and accumulate more valuable bioproducts, such as amino acids and fatty acids (especially medium-chain and long-chain fatty acids) and other biomass compounds, the hydrolysis efficiency in the subsequent anaerobic digestion process will be enhanced, thereby improving the overall efficiency and economic viability of the resource recovery process^6^.

Recent advancements in wastewater treatment technologies have shifted focus towards resource recovery, with methods such as Chemical Enhanced Primary Treatment (CEPT), High-Rate Activated Sludge (HRAS), and various anaerobic processes being developed^7–9^. Placing these processes before biological treatment units allows for the capture of carbon sources that would otherwise be oxidized and consumed, storing them as organic matter in the activated sludge biomass^10^. This biomass can then be fed into anaerobic digestion units to produce methane and digestate^11^. However, these pre-treatment processes for capturing carbon face limitations, including complex process control, high operational costs, and challenges in treating low-concentration wastewater efficiently^10^. Enhancing the capture and enrichment efficiency of organic matter without increasing process complexity has become an important direction in current research.

The effects of varying DO levels on microbial communities have been extensively studied, particularly in the context of hypoxia. Under low oxygen conditions, many microorganisms shift from aerobic respiration to anaerobic respiration, leading to significant changes in metabolite production^12^. Recent research has shown that intermittent aeration or alternating aerobic/anoxic strategies offer advantages in improving wastewater treatment efficiency and resource recovery^13,14^. For instance, studies have demonstrated that intermittent aeration can enhance total nitrogen removal, reduce energy consumption, and optimize certain metabolite production^15–17^. However, oxygen modulation as a strategy to direct microbial metabolism toward the production of specific high-value chemicals in activated sludge biomass, particularly amino acids and fatty acids, remains largely unexplored.

Mechanistically, changes in oxygen concentration reconfigure metabolic equilibrium primarily through two key regulatory factors, FNR and ArcA^18,19^. Under low oxygen conditions, FNR promotes anaerobic respiration^20^, while ArcA inhibits genes associated with oxidative TCA cycle enzymes^21^. This results in reduced carbon flux through oxidative pathways, channeling resources into synthetic pathways for amino acids and fatty acids^22^. Additionally, ArcA’s inhibition of carbon source oxidation decreases the catabolic rates of various compounds, including amino acids and fatty acids^19,23^. As microbes adapt to varying oxygen levels, dynamic adjustments of enzyme expression by oxyregulons drive the flow of ingested carbon toward the accumulation of these valuable metabolites^24^.

We hypothesized that by intentionally changing oxygen availability in wastewater treatment systems to keep microorganisms in alternating aerobic and anoxic states, we could strategically manipulate activated sludge microbial pathways to produce targeted, value-added chemicals. This approach could provide an alternative that aids in resource recovery, avoiding significant modifications to existing wastewater treatment systems while achieving high-quality chemical recovery and reducing aeration energy consumption.

This study aimed to investigate the impact of controlled DO oscillations on microbial biosynthesis of valuable biochemicals in activated sludge. Specifically, we sought to determine whether high-frequency oxygen perturbation conditions, with or without obvious anoxic phases, can enhance the accumulation of organic metabolites, particularly amino acids and fatty acids, within the sludge microbiomes. Furthermore, we explored the mechanistic effects of oxygen variations on regulon activation and the subsequent diversion of carbon flux toward amino acid and fatty acid metabolic pathways.

Additionally, this research encompassed a comprehensive analysis of wastewater treatment performance, including carbon removal and nitrogen conversion, biomass growth and compositional changes in activated sludge, shifts in overall microbial enzymatic functions, and oxidative stress-induced antioxidant mechanisms. These analyses provide a holistic understanding of the processes involved, offering valuable insights for both researchers and practitioners in the field of wastewater treatment and resource recovery.

## 2. Materials and methods

### 2.1 Bioreactor operation and performance

Activated sludge for this study was obtained from the Māngere Wastewater Treatment Plant in Auckland, New Zealand. Three identical acrylic cylindrical bioreactors, each having an effective volume of 1 L, were operated in parallel under different aeration conditions at 20□. The same activated sludge was run thrice as biological triplicates. Each 1 L volume of the microbial reaction solution contained 2.75 g/L mixed liquor-suspended solid (MLSS) activated sludge, 3.84 g/L NaHCO_3_ as inorganic carbon, and 1 mL trace element solution (Table S1). For 48 hours, 50 mL of concentrated artificial wastewater (Table S2) was distributed equally and continuously fed into the system using a syringe pump and mixed with a magnetic stirrer.

Three aeration conditions included Constant Aeration (CA) (stable at 1.8-2.2 mg/L DO), Continuous Perturbation (CP) (DO varying between 0.1 and 2.0 mg/L in ∼3 min cycles), and Intermittent Perturbation (IP) (DO varying between 0 and 2.0 mg/L; aerobic time: anoxic time = 3 min:3 min) (Figure 1a, 1b, 1c). DO concentration monitoring and aeration control were achieved using an Arduino Mega2560-based microcontroller system^25^. Rapid DO increases in CP and IP were accomplished by controlling the airflow to 1.5 L/min. Aeration stopped automatically when DO reached 2.0 mg/L, allowing DO to fall naturally by microbial consumption. A lower aeration rate of 0.5 L/min stabilized DO for CA.

**Fig. 1:**
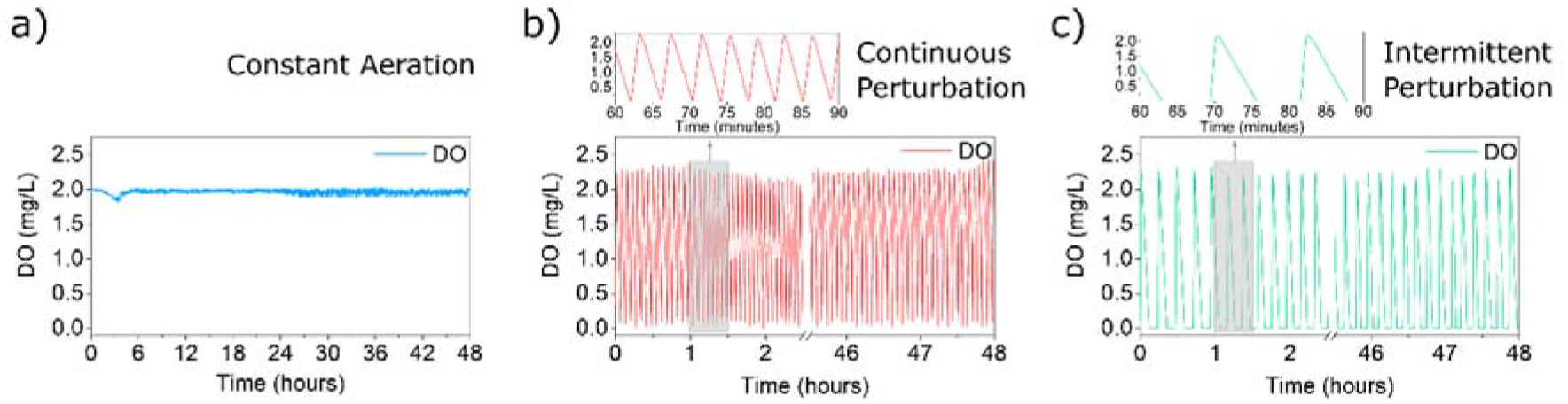
Dissolved oxygen concentrations controlled under different aeration strategies: (a) Constant Aeration maintains a stable dissolved oxygen (DO) level at 2.0±0.2 mg/L. (b) Continuous Perturbation involves continuously varying the DO concentration between 0.1 and 2.0 mg/L without intermission. (c) Intermittent Perturbation varies the DO concentration between 0 and 2.0 mg/L, with an equal duration of aerobic and anoxic periods.

### 2.2 Determination of amino acids, fatty acids and other metabolites

Activated sludge samples were dried, ground, and digested, then analyzed for organic carbon and nitrogen using a TOC and TN analyzer (Shimadzu, Japan). The total concentrations of free amino acids and fatty acids per unit dry weight of activated sludge were measured by L-Amino Acid Assay Kit (Abcam, UK) and Free Fatty Acid Assay Kit (Abcam, UK), respectively, according to the manufacturers’ instructions.

The analysis of individual intracellular metabolites in activated sludge was conducted using gas chromatography-mass spectrometry to assess the changes in metabolic profiles under various experimental conditions. This involved metabolite extraction and methyl chloroformate derivatization, with both biological and technical replicates included for robustness. A specific focus was placed on the relative abundance of free amino acids and fatty acids as indicators of metabolic adaptation. Detailed metabolite extraction, analysis, and data processing procedures are documented in the supplementary material (Section S1.3).

### 2.3 Metaproteomics analysis

Non-targeted metaproteomic analysis of sludge samples was performed to explore microbial protein expression under various experimental conditions. This involved collecting three biological replicates per condition, followed by protein extraction, quantification, and analysis using nanoscale liquid chromatography coupled to tandem mass spectrometry. This analysis provided insights into the microbial community’s response to different oxygen perturbation strategies. The detailed methodology, including sample preparation, protein digestion, and mass spectrometry analysis, is available in the Supplementary Materials (Section S1.4).

In the acquired metaproteomic dataset, the relative abundance of 4850 identified proteins from various microbial species before and after treatment under different aeration conditions was analyzed. To examine differences in the overall metabolic function of activated sludge, the differential abundance of functional enzymes was analyzed, treating the microbial system as a unified entity (Section 3.1). This consolidation refined the dataset to 1,110 distinct proteins. For a detailed exploration of enzyme profile changes and their implications, refer to Supplementary Materials (Section S2.3).

The unconsolidated protein abundance data were used for mechanistic analysis. Changes in enzyme abundance regulated by regulons (FNR, ArcA, SoxRS, OxyR, NorR, and NsrR) after 48 hours of aeration treatment (T48) were compared with non-aeration treatment (T0) (Result 3.3). Differences in enzyme abundance between CP, IP, and CA treatments were also determined (Result 3.4). Data were analyzed using paired t-test statistical methods. Enzyme information was obtained from the RegulonDB database^26^.

## 3. Results

### 3.1. Effect of different oxygen perturbations on metabolite abundance in activated sludge

Carbon and nitrogen transformation processes were investigated in activated sludge from wastewater treatment systems exposed to the aeration treatments of constant aeration, continuous perturbation, and intermittent perturbation (Table 1). All three aeration regimes effectively reduced COD to below 50 mg/L, demonstrating efficient carbon removal. However, obvious differences were observed in nitrogen conversion. Under CA and CP treatments, the process was predominantly characterized by nitrification (Figure 2c). CA and CP demonstrated high nitrification efficiency, with 94.3±1.4% and 81.7±2.1% of ammonia nitrogen accumulating as nitrate nitrogen respectively. IP showed a lower ammonia conversion rate but performed both nitrification and denitrification (Figure 2c). IP achieved the highest nitrogen removal percentages, with 71.3 ± 4.5% of consumed ammonia nitrogen being released in gaseous form. Detailed findings are provided in Supplementary Material (Section S2.1).

**Table 1.**
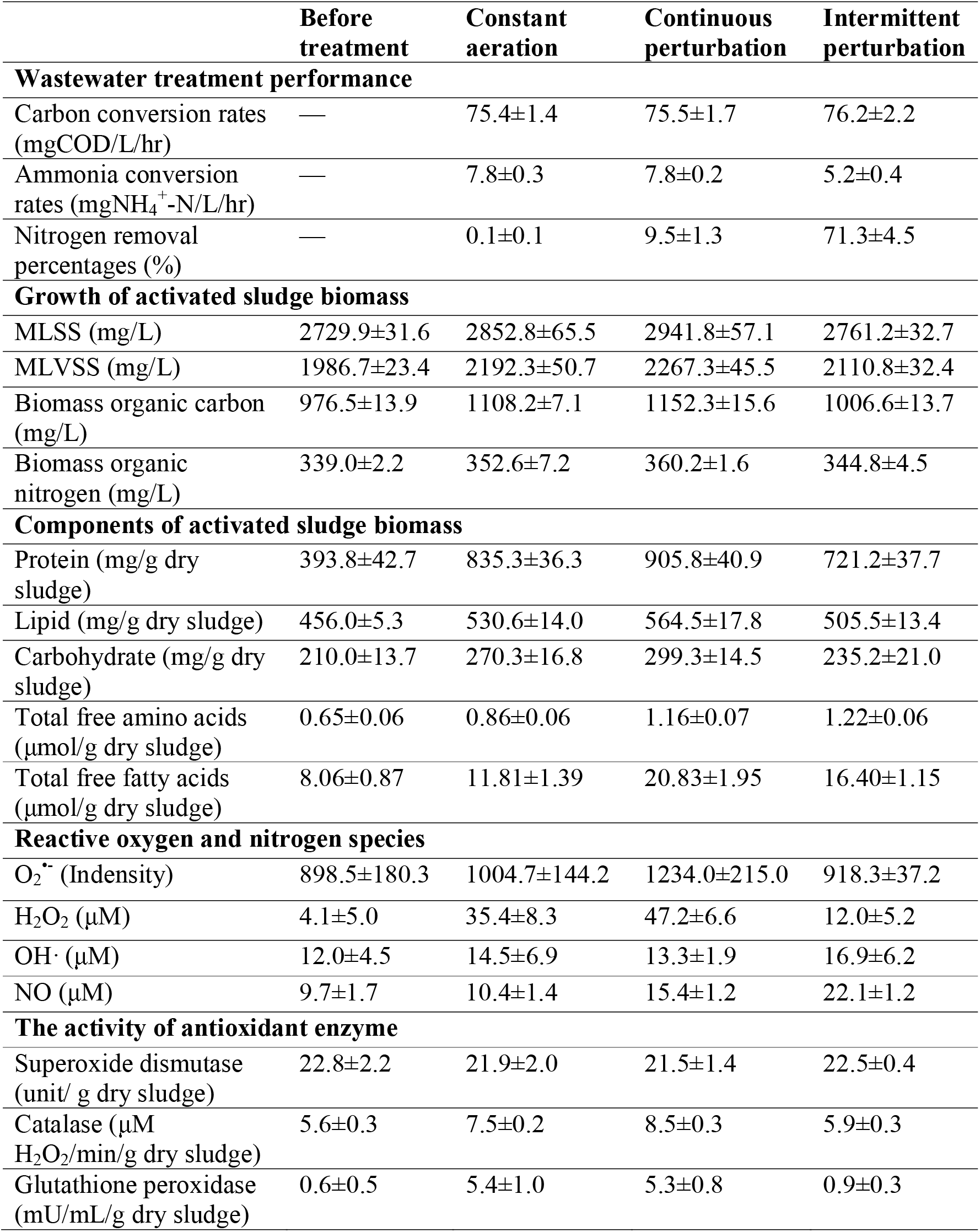
Wastewater treatment performance, activated sludge growth, composition, and oxidative stress before and after 48 hours of exposure to different aeration conditions

|  | Before treatment | Constant aeration | Continuous perturbation | Intermittent perturbation |
| --- | --- | --- | --- | --- |
| **Wastewater treatment performance** |  |  |  |  |
| Carbon conversion rates (mgCOD/L/hr) | — | 75.4±1.4 | 75.5±1.7 | 76.2±2.2 |
| Ammonia conversion rates (mgNH<sub>4</sub><sup>+</sup>-N/L/hr) | — | 7.8±0.3 | 7.8±0.2 | 5.2±0.4 |
| Nitrogen removal percentages (%) | — | 0.1±0.1 | 9.5±1.3 | 71.3±4.5 |
| **Growth of activated sludge biomass** |  |  |  |  |
| MLSS (mg/L) | 2729.9±31.6 | 2852.8±65.5 | 2941.8±57.1 | 2761.2±32.7 |
| MLVSS (mg/L) | 1986.7±23.4 | 2192.3±50.7 | 2267.3±45.5 | 2110.8±32.4 |
| Biomass organic carbon (mg/L) | 976.5±13.9 | 1108.2±7.1 | 1152.3±15.6 | 1006.6±13.7 |
| Biomass organic nitrogen (mg/L) | 339.0±2.2 | 352.6±7.2 | 360.2±1.6 | 344.8±4.5 |
| **Components of activated sludge biomass** |  |  |  |  |
| Protein (mg/g dry sludge) | 393.8±42.7 | 835.3±36.3 | 905.8±40.9 | 721.2±37.7 |
| Lipid (mg/g dry sludge) | 456.0±5.3 | 530.6±14.0 | 564.5±17.8 | 505.5±13.4 |
| Carbohydrate (mg/g dry sludge) | 210.0±13.7 | 270.3±16.8 | 299.3±14.5 | 235.2±21.0 |
| Total free amino acids (μmol/g dry sludge) | 0.65±0.06 | 0.86±0.06 | 1.16±0.07 | 1.22±0.06 |
| Total free fatty acids (μmol/g dry sludge) | 8.06±0.87 | 11.81±1.39 | 20.83±1.95 | 16.40±1.15 |
| **Reactive oxygen and nitrogen species** |  |  |  |  |
| O<sub>2</sub><sup>•-</sup> (Intensity) | 898.5±180.3 | 1004.7±144.2 | 1234.0±215.0 | 918.3±37.2 |
| H<sub>2</sub>O<sub>2</sub> (μM) | 4.1±5.0 | 35.4±8.3 | 47.2±6.6 | 12.0±5.2 |
| OH<sup>•</sup> (μM) | 12.0±4.5 | 14.5±6.9 | 13.3±1.9 | 16.9±6.2 |
| NO (μM) | 9.7±1.7 | 10.4±1.4 | 15.4±1.2 | 22.1±1.2 |
| **The activity of antioxidant enzyme** |  |  |  |  |
| Superoxide dismutase (unit/ g dry sludge) | 22.8±2.2 | 21.9±2.0 | 21.5±1.4 | 22.5±0.4 |
| Catalase (μM H<sub>2</sub>O<sub>2</sub>/min/g dry sludge) | 5.6±0.3 | 7.5±0.2 | 8.5±0.3 | 5.9±0.3 |
| Glutathione peroxidase (mU/mL/g dry sludge) | 0.6±0.5 | 5.4±1.0 | 5.3±0.8 | 0.9±0.3 |

**Fig. 2:**
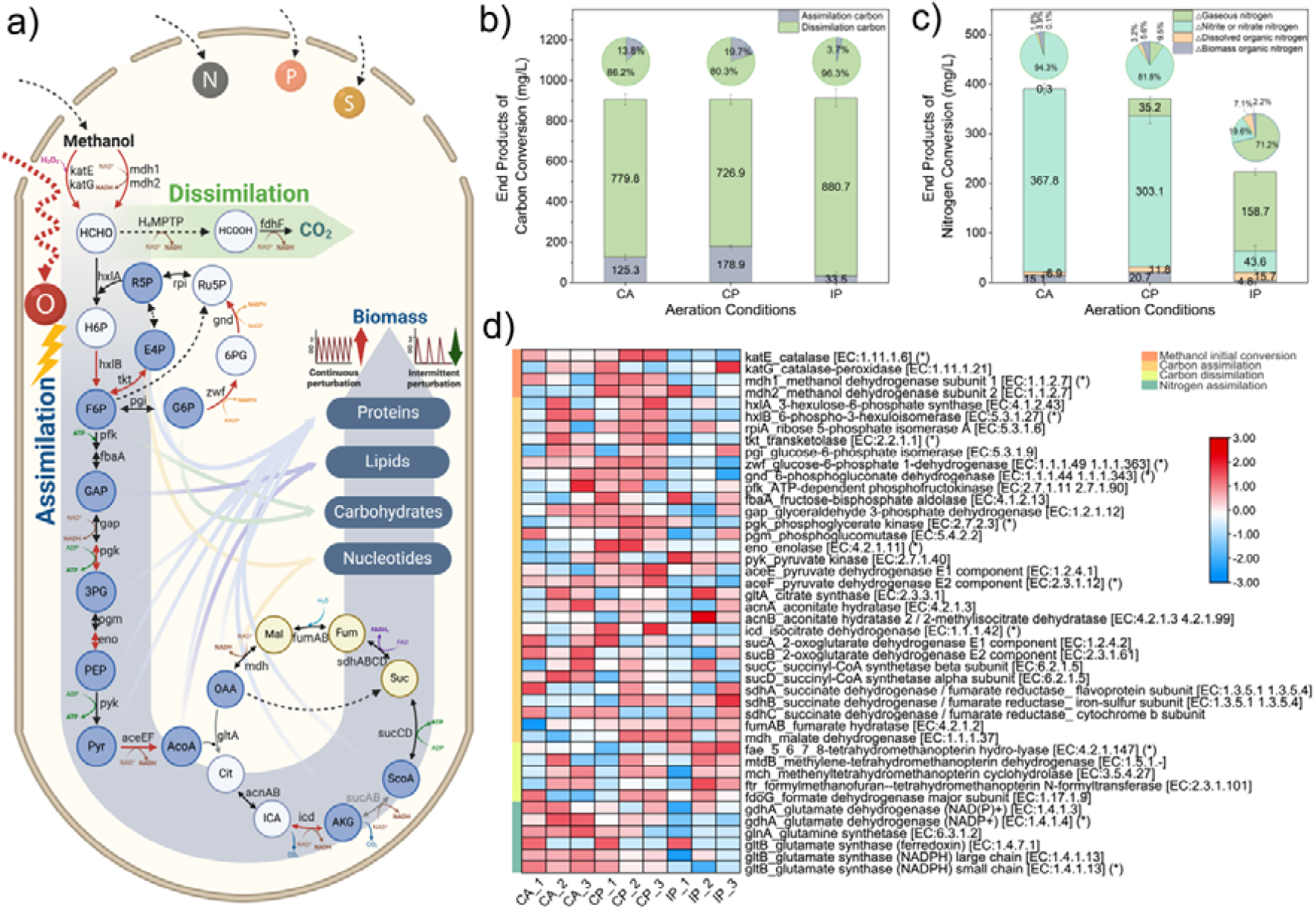
Higher biomass assimilation efficiency in activated sludge systems under continuous perturbation (CP) compared to intermittent perturbation (IP). (a) Schematic of carbon (methanol) uptake, assimilation, and dissimilation pathways in activated sludge systems, with reactions marked in red indicating significantly higher enzyme abundance under CP compared to IP. (b) Amounts of assimilated and dissimilated carbon in activated sludge systems after 48 hours under three aeration conditions. (c) Amounts of ammonia nitrogen converted to different final nitrogen products in activated sludge systems after 48 hours under three aeration conditions. (d) Heatmap showing the differences in enzyme abundances related to initial methanol conversion, carbon assimilation, carbon dissimilation, and nitrogen assimilation after 48 hours under three aeration conditions, with (*) indicating significant differences between CP and IP (T-test, *P*<0.05). Samples under each aeration condition include three biological replicates; Error bars represent standard deviations.

After 48 hours of aeration treatment, there were significant treatment-specific differences in biomass growth and the content of organic components (Table 1). CP yielded the highest biomass synthesis, converting 19.8±1.0% of carbon and 5.6±0.2% of ammonia nitrogen into biomass. This surpassed CA, which converted 13.9±1.7% of carbon and 3.9±1.6% of nitrogen (Figure 2b, 2c). CP also resulted in the highest protein and lipid content increases, with protein content rising from 144.2±15.1 mg/g to 313.6±5.1 mg/g, and lipid content increasing from 167.0±2.9 mg/g to 191.9±3.1 mg/g. In contrast, IP showed the lowest biomass synthesis, converting only 3.7±2.4% of carbon and 2.1±1.7% of ammonia nitrogen into biomass (Figure 2b, 2c), accompanied by the slightest increase in protein and lipid content. Further details, including MLSS, MLVSS, biomass 3D-EEM analysis, protein, lipid, and carbohydrate, are provided in Supplementary Material (Section S2.2).

The microbial ecosystem within the activated sludge is a consortium composed of diverse species. To evaluate its collective capacity for assimilation under different aeration conditions, we analyzed a dataset of the relative abundances of 1,110 identified proteins after 48 hours of treatment across three aeration patterns. Detail discussed below focuses on the analysis of enzymes associated with the assimilatory metabolism for biomass synthesis. A more comprehensive exploration of other changes to enzyme profiles and their implications for systemic metabolic functions is provided in Supplementary Material (Section S2.3).

The metabolic schematic in Figure 2 illustrates the differences in abundance of a suite of enzymes involved in methanol conversion and biomass production under CP and IP. The abundance of key enzymes supporting assimilation was higher under CP conditions compared with IP conditions (T-test, *P*<0.05) (Figure 2d). These enzymes are involved in various metabolic processes, including the conversion of methanol, with examples such as catalase (katE) and methanol dehydrogenase (mdh1); generation of biomass precursors, highlighted by enzymes like 6-phospho-3-hexuloisomerase (hxlB), transketolase (tkt), phosphoglycerate kinase (pgk), pyruvate dehydrogenase (aceF), enolase (eno), and isocitrate dehydrogenase (icd); the production of NADPH, as seen in G6PD (zwf) and 6PGD (gnd); and nitrogen assimilation enzymes including glutamate dehydrogenase (gdhA) and glutamate synthase (gltB).

This differential expression underscores the impact of the oxygen-disturbed state, particularly the anoxic phase within the perturbation cycles, on the metabolic state of the entire microbial system. The successive stimuli in CP appear to induce an adaptive upregulation in microbes, potentially optimizing the assimilation of carbon sources into cellular biomass. In contrast, the presence of anoxic phases in IP may reduce microbial assimilation, likely a conservative strategy in response to insufficient oxygen supply. However, we found that the activated sludge systems under both CP and IP conditions exhibited a stronger capability of enzymes to compete for oxygen compared to CA conditions, primarily manifested by a significant increase in the induction of cytochrome cbb3-type oxidase (Figure S18). This enhancement aids them in utilizing the electron transport chain (ETC) to generate energy under unstable oxygen supply conditions.

### 3.2. Both continuous and intermittent perturbations promoted the contents of free amino acids and fatty acids in biomass

We investigated the impact of oxygen perturbation on free amino acids and fatty acids in activated sludge biomass under different aeration patterns (Table 1). Compared to the initial state (T0), where the total amino acid content was 0.65±0.06 μmol/g dry sludge, there was an increase under continuous perturbation (1.16±0.07 μmol/g dry sludge) and intermittent perturbation (1.22±0.06 μmol/g dry sludge). These levels were 35.7±7.6% and 42.4±8.1% higher than under constant aeration (0.86±0.06 μmol/g dry sludge), respectively. Similarly, the total fatty acid content was 8.06±0.87 μmol/g dry sludge at T0, and elevated to 20.83±1.95 μmol/g dry sludge under continuous perturbation and 16.40±1.15 μmol/g dry sludge under intermittent perturbation, representing 76.4±13.0% and 39.3±7.7% higher levels respectively, compared to constant aeration (11.81±1.39 μmol/g dry sludge) (Figure 3a, 3b).

**Fig. 3:**
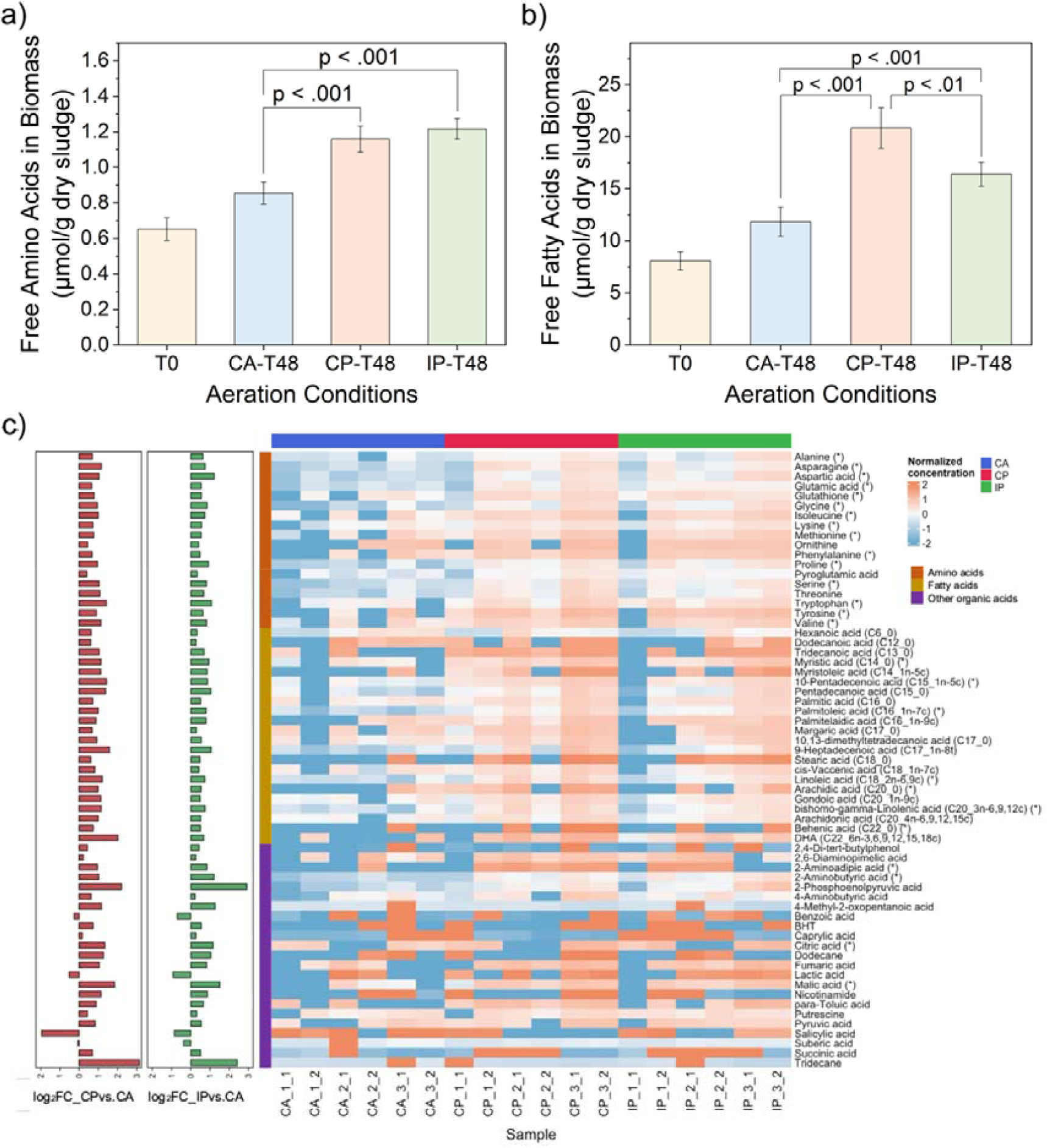
Oxygen perturbations increased the content of free amino acids and fatty acids in activated sludge biomass. (a) The content of total free amino acids in biomass. (b) The content of total free fatty acids in biomass. (c) The relative abundance of intracellular primary metabolites (amino acids, fatty acids, and organic acids) of microorganisms in activated sludge systems after 48 hours of exposure to different aeration strategies. The data is normalized by an internal standard, subtracting T0 values, and with log_10_ transformation and Pareto scaling. An asterisk (*) signifies that the abundance of this metabolite is significantly higher in the Continuous Perturbation (CP) and Intermittent Perturbation (IP) treatments after 48 hours compared to CA (ANOVA Tukey test, *P*<0.05). The bar chart on the left shows the log_2_ fold change in the relative abundance of metabolites in activated sludge exposed to oxygen perturbations compared with constant aeration (technical duplication results for biological triplicates; n=6).

The analysis of the relative abundance of individual metabolites also showed that the main organic acids that increased in abundance under oxygen perturbations compared with CA in the biomass belonged to the amino acid and fatty acid groups. Of the 63 metabolites identified, 26 showed significantly higher abundance under CP and IP than under CA (Figure 3c). The observed increases in the abundance of 14 out of 16 amino acids and 7 out of 22 fatty acids were due to oxygen perturbations. This broad increase in amino acids reflects a systemic impact of oxygen perturbation on microbial metabolism. The differentially produced amino acids include alanine, asparagine, aspartic acid, glutamic acid, glycine, isoleucine, lysine, methionine, phenylalanine, proline, pyroglutamic acid, serine, tryptophan, tyrosine, and valine, indicating an extensive and beneficial metabolic adaptation to oxygen perturbations. Moreover, under CP and IP conditions, there was a significant shift in the fatty acid profile relative to CA. The fatty acids that increased in abundance belonged to the categories of long-chain fatty acids and very long-chain fatty acids. This group includes myristic acid (C14_0), 10-pentadecenoic acid (C15_1n-5c), palmitoleic acid (C16_1n-7c), linoleic acid (C18_2n-6,9c), arachidic acid (C20_0), bishomo-gamma-linolenic acid (C20_3n-6,9,12c), and behenic acid (C22_0), which are of higher value for bioresource applications.

### 3.3. Activation of regulons by oxygen variation and reactive oxygen and nitrogen species under oxygen perturbations

During the 48-hour aeration treatments, significant changes were observed in the concentration of reactive oxygen and nitrogen species (RONS) within the activated sludge system (Table 1). Specifically, CP conditions resulted in H_2_O_2_ concentrations ranging between 32-56 μM, compared to 21-45 μM under CA and below 20 μM under IP. NO concentrations were highest under IP (16-24 μM), followed by CP (11-18 μM) and CA (9-13 μM). These changes in RONS levels corresponded with alterations in antioxidant enzyme activities (Table 1). While SOD activity showed no significant differences, CAT and GSH-POD activities, which convert H_2_O_2_, were specifically affected by aeration patterns, with increases observed under CA and CP conditions. For more detailed information, please refer to Supplementary Material (Section S2.5). The results indicate that the metabolic response of the microbial system under oxygen perturbation is related not only to changes in oxygen concentration but also to elevated levels of RONS. Therefore, the regulons studied in this section include oxygen-responsive regulons (FNR and ArcA), ROS-responsive regulons (SoxRS and OxyR), and NO-responsive regulons (NorR and NsrR).

Additionally, by analyzing the metaproteomic data, we found that the orthologous enzyme is regulated differently by different species. Merging data for the orthologous proteins across different species is only suitable for assessing the overall level of enzyme functions, but cannot serve as a basis for exploring the response mechanisms of individual microbes to oxygen perturbations. Therefore, we utilized a refined dataset of 4,850 proteins distinguished by microbial origin for mechanistic exploration. This approach allowed us to infer the responses of individual microbes within a mixed community. Figure 4 displays the enzymes regulated by the regulons of oxygen and stress response (FNR, ArcA, SoxRS, OxyR, NorR, and NsrR), in which there were statistically significant changes in abundance after 48 hours of aeration treatment (adjusted *P*<0.05, |log_2_FC|>0.58). It is worth noting that within the range of enzymes regulated by the regulons of interest, more enzymes were significantly up- or down-regulated under CP and IP conditions than under CA conditions. Further detail on different enzyme expressions under the different aeration conditions is provided in Table S7, S8, and S9.

**Fig. 4:**
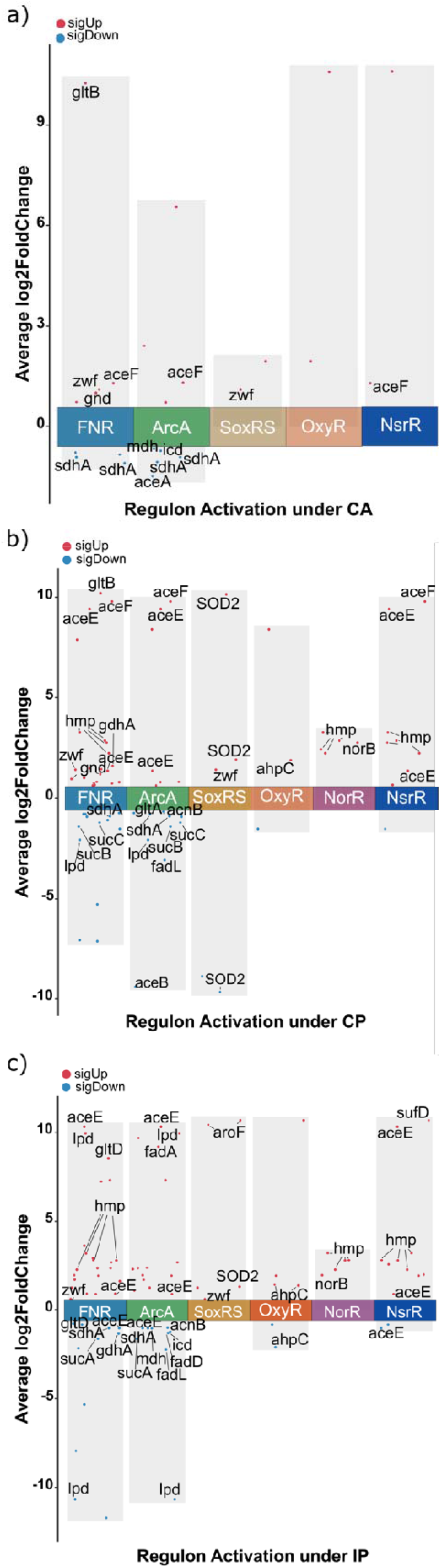
The enzymes with significant changes in abundance after 48 hours of treatment under three aeration strategies, grouped by their respective regulons. CA, CP, and IP represent constant aeration, continuous perturbation, and intermittent perturbation, respectively. (*P*<0.05, |log_2_FC|>0.58).

For the subset of enzymes regulated by the FNR and ArcA regulons, pronounced changes in relative levels were observed under CP and IP conditions compared to CA conditions after 48 hours of treatment. This regulation affected enzymes involved in specific pathways including carbon assimilation and ammonia assimilation pathways. In the carbon assimilation pathway, oxygen perturbations resulted in a greater number of key enzymes with decreased abundance. Under CA conditions, succinate dehydrogenase (sdhA) was downregulated (sdhA with log_2_FCs of -1.07 and -0.92) while pyruvate dehydrogenase (aceF) was upregulated (log_2_FC of 1.31). Regulation of these two enzymes was similar under steady oxygen and oxygen perturbations, but more other enzymes showed a decreasing abundance trend under oxygen perturbations. Enzymes significantly regulated under both CP and IP conditions included succinate dehydrogenase (sdhA with log_2_FCs of -0.89 and -0.78 under CP; -1.02 under IP), pyruvate dehydrogenase (aceF with log_2_FC of 9.82 and aceE with log_2_FCs of 9.42 and 1.39 under CP; aceE with log_2_FCs of 10.34, -0.99 and 0.91 under IP), dihydrolipoamide dehydrogenase (lpd with log_2_FC of -2.07 under CP; and -10.651, 9.96 under IP), and 2-oxoglutarate dehydrogenase (sucB with log_2_FC of -1.41 under CP; sucA with log_2_FC of -1.60 under IP). Significant downregulation under CP only was observed for succinyl-CoA synthetase (sucC with log_2_FC of -1.20) and citrate synthase (gltA with log_2_FC of -0.66). Under IP only, aconitate hydratase (acnB with log_2_FC of -0.99) and malate dehydrogenase (mdh with log_2_FC of -1.02) were significantly downregulated.

Within the carbon assimilation pathway, the pentose phosphate pathway, which produces NADPH, was upregulated after 48 hours of treatment under all three aeration conditions, albeit to varying extents. Expression under CA was similar to CP; glucose-6-phosphate 1-dehydrogenase (G6PD) (zwf with log_2_FC of 1.12 under CA; 1.42 under CP) and 6-phosphogluconate dehydrogenase (6PGD) (gnd with log_2_FCs of 1.03 under CA; 1.21, 0.97 under CP) were significantly upregulated. Under IP, only glucose-6-phosphate 1-dehydrogenase (zwf with log_2_FC of 0.61) was significantly upregulated.

Regarding the ammonia assimilation pathway, CP and IP conditions showed different regulation responses on key enzymes compared to the CA condition. Under CA, only glutamate synthase (gltB with log_2_FC of 10.25) was significantly upregulated, while CP led to significant upregulation of both glutamate synthase (gltB with log_2_FC of 10.23) and glutamate dehydrogenase (gdhA with log_2_FC of 1.61). In contrast, under IP, glutamate synthase (gltD with log_2_FCs of -0.76, 8.57) and glutamate dehydrogenase (gdhA with log_2_FC of -1.32) exhibited significant regulation.

SoxRS, OxyR, NorR, and NsrR regulate microbial stress metabolism in response to RONS concentrations. Compared to CA conditions, oxygen perturbations (CP and IP) resulted in the regulation of a greater number of antioxidative stress enzymes. Within the panel of enzymes regulated by SoxRS and OxyR, SOD and peroxiredoxin, responsible for converting O_2_^•-^ and H_2_O_2_ respectively, there was significant regulation under CP and IP conditions but not under CA. SOD (SOD2 with a log_2_FC of 10.16, -8.87, and 1.93 under CP; 1.34 under IP) and peroxiredoxin (ahpC with log_2_FC of 1.91 under CP; 1.45, 1.40, and -2.07 under IP) showed significant regulation.

NorR and NsrR, NO stress sensors, regulate nitric oxide dioxygenase and nitric oxide reductase expression for NO detoxification. Both enzymes were significantly upregulated under oxygen perturbations. Nitric oxide dioxygenase showed substantial upregulation (hmp showed log_2_FCs of 3.30, 2.88, 2.78, and 2.24 under CP; 3.22, 2.82, 2.80, and 2.29 under IP), as did nitric oxide reductase (norB had a log_2_FC of 2.45 under CP; 1.97 under IP).

### 3.4. Oxygen perturbations affect expression of amino acid and fatty acid metabolism enzymes

Metaproteomic data analysis targeting regulatory elements has revealed that oxygen perturbations activate regulons controlling carbon and nitrogen metabolism pathways, likely impacting amino acid and fatty acid metabolism differently compared to stable oxygen. Therefore, we compared whether there were significant differences in the abundance of enzymes related to amino acid and fatty acid metabolism after treatment under oxygen perturbation conditions versus stable aeration. The results revealed that, compared to CA condition, some key enzymes involved in amino acid and fatty acid synthesis showed significantly higher abundance under both oxygen perturbation conditions.

We calculated log_2_FC in protein abundance to compare the effects of oxygen perturbations against stable oxygen conditions. All log_2_FC values presented in this section are comparisons to the CA condition. Figure 5 depicts the differential enzyme abundance for CP and IP conditions compared to CA. In both Figure 5a and 5b, enzymes with statistically significant abundance differences (*P*<0.05, log_2_FC>0) are highlighted, providing a visual representation of differential metabolic changes caused by oxygen perturbation compared to no perturbation. Some enzymes crucial to metabolism are marked with their regulatory gene IDs in the figures, and detailed enzyme information can be found in Table S11 and S12.

**Fig. 5:**
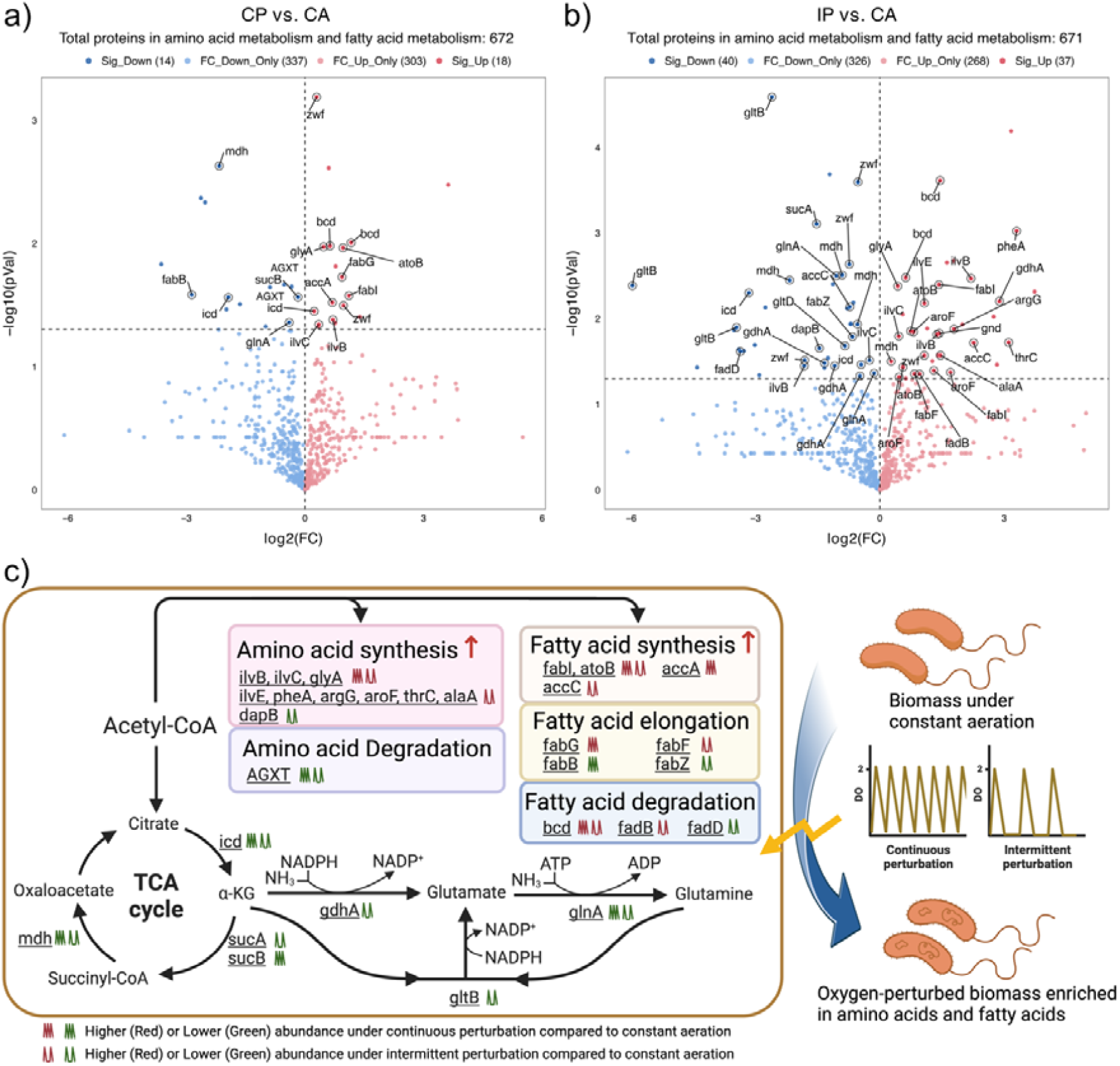
Oxygen perturbation redirects carbon flux from the TCA cycle to amino acid and fatty acid synthesis: (a) and (b) Comparison of enzymes with significant differential abundance related to amino acid and fatty acid metabolism in activated sludge microbial systems after 48 hours of continuous perturbation (CP) and intermittent perturbation (IP) treatments relative to constant aeration (CA) treatment; (c) Metabolic regulation schematic under oxygen perturbations, where red indicates significantly higher enzyme expression compared to CA, and green indicates significantly lower (*P*<0.05).

Compared to CA, some key enzymes in the TCA cycle showed significantly lower abundance in microbial systems after CP and IP treatments, including malate dehydrogenase (mdh with log_2_FC of -2.17 for CP; -2.18, -0.91, -0.54, and 0.27 for IP), isocitrate dehydrogenase (icd with log_2_FC of -0.94, and 0.23 for CP; -3.16, and -0.45 for IP), and 2-oxoglutarate dehydrogenase (sucB with log2FC of -0.18 for CP; sucA with log_2_FC of -1.53 for IP). The same scenario occurs in nitrogen fixation metabolism, such as glutamate dehydrogenase (gdhA with log_2_FC of -1.34, -1.09, -0.48, and 2.88 for IP), glutamine synthetase (glnA with log_2_FC of -0.40 for CP; -1.05, and -0.14 for IP), and glutamate synthase (gltB with log_2_FC of -5.97, -3.46, and -2.60 for IP).

For amino acid metabolism, the ilv cluster is crucial in the branched-chain amino acid synthesis pathway, and enzymes showed higher abundance under oxygen perturbation. Specifically, acetolactate synthase (ilvB with log_2_FC of 0.71 for CP; 2.20, 1.38, and -1.82 for IP), ketol-acid reductoisomerase (ilvC with log_2_FC of 0.35 for CP; 0.46, and -0.25 for IP), and branched-chain amino acid aminotransferase (ilvE with log_2_FC of 1.07 for IP). Other enzymes involved in amino acid pathways also showed higher abundance under oxygen perturbation, especially under IP compared to stable oxygen. For example, those involved in glycine synthesis (glycine hydroxymethyltransferase, glyA, log_2_FC of 0.47 for CP); biosynthesis of phenylalanine and tyrosine (chorismate mutase/prephenate dehydratase, pheA, log_2_FC of 3.30 for IP); arginine biosynthesis (argininosuccinate synthase, argG, log_2_FC of 1.79 for IP); tryptophan biosynthesis (3-deoxy-7-phosphoheptulonate synthase, aroF, log_2_FC of 1.70, 0.81, and 0.46 for IP), threonine synthesis (threonine synthase, thrC, log_2_FC of 3.11 for IP); and alanine biosynthesis (alanine-synthesizing transaminase, alaA, log_2_FC of 1.46, and -1.21 for IP). The alanine-glyoxylate transaminase/serine-glyoxylate transaminase/serine-pyruvate transaminase (AGXT with log_2_FC of -0.34 for CP; -0.79 for IP) showed lower abundance under both oxygen perturbation conditions, which might lead to the accumulation of their substrates, alanine and serine.

For fatty acids, the acc cluster is the first enzyme in fatty acid biosynthesis, catalyzing the conversion of acetyl-CoA to malonyl-CoA, which is a rate-limiting step. It showed higher enzyme abundance under oxygen perturbation: acetyl-CoA carboxylase (accA with log_2_FC of 0.69 for CP and accC with log_2_FC of 2.26 and -0.72 for IP). The fab cluster, involved in fatty acid biosynthesis and elongation, exhibited various regulatory expression patterns, with the enzyme enoyl-[acyl-carrier protein] reductase (fabI with log_2_FC of 1.11 for CP; accC with log_2_FC of 1.43 for IP) showing higher abundance under oxygen perturbation. The 3-oxoacyl-[acyl-carrier protein] reductase (fabG with log_2_FC of 0.96 for CP) involved in the fatty acid chain elongation cycle showed higher abundance under CP, as did the 3-oxoacyl-[acyl-carrier-protein] synthase (fabF with log_2_FC of 0.83 for IP) involved in extending the fatty acid chain under IP. The 3-oxoacyl-[acyl-carrier-protein]_synthase (fabB with log_2_FC of - 2.86 for CP) involved in unsaturated fatty acid synthesis showed lower abundance under CP. For fatty acid degradation, IP conditions seem more favorable for the accumulation of long-chain fatty acids rather than short-chain fatty acids. Long-chain fatty acids need the long-chain acyl-CoA synthetase (fadD with log_2_FC of -3.37 for IP) to participate, and its abundance is lower under IP. The 3-hydroxybutyryl-CoA dehydrogenase (fadB with log_2_FC of 0.96 for IP), which helps in the degradation of short-chain fatty acids, showed higher abundance under IP conditions.

## 4. Discussion

In traditional activated sludge treatment systems, the oxygen supply is often continuous and constant, which may not be conducive to achieving an optimal balance between energy consumption and treatment outcomes. In contrast, by adopting variable oxygen supply strategies, the flexibility and adaptability of microbial metabolism can be stimulated, facilitating specific biochemical reactions^27^. The strategic oxygen perturbations in activated sludge systems revealed a complex interplay between oxygen supply regimes, treatment performance, and metabolite synthesis^28–30^. The CP conditions, characterized by oxygen fluctuations between 0-2 mg/L, demonstrated an intriguing adaptive response in aerobic respiration microorganisms. The upregulation of cytochrome cbb3-type oxidase, known for its high oxygen affinity, suggests a metabolic shift towards more efficient oxygen utilization under variable conditions^31^. This adaptation aligns with the observed maintenance of organic matter and ammonia nitrogen oxidation rates, coupled with increased biomass synthesis efficiency. These findings indicate that timely oxygen resupply can effectively mitigate the energy uncoupling effects often associated with oxygen stress, maintaining aerobic respiration efficiency and reducing non-functional ATP loss. In contrast, the IP conditions, which introduced anoxic phases, revealed a trade-off between oxidation and reduction efficiency. The observed reduction in carbon and nitrogen oxidation rates, accompanied by enhanced nitrate respiration, suggests a potential for optimizing overall nitrogen removal in wastewater treatment systems. The concomitant reduction in biomass synthesis under IP conditions not only corroborates the energy uncoupling theory^32^ but also presents an opportunity for sludge reduction strategies, addressing a persistent challenge in wastewater treatment operations^33–35^.

A key finding of this study is the significant increase in free amino acids and fatty acids under both CP and IP conditions. The observed shift towards long-chain fatty acids (≥C12) and the specific amino acid profile changes suggest a metabolic shift in response to frequent switching between aerobic and anoxic. This adaptive response likely contributes to cellular homeostasis, energy storage, and stress signaling mechanisms^36–42^. The correlation between enzyme upregulation and increased metabolite levels provides mechanistic insight into these metabolic shifts, offering potential targets for bioprocess optimization. The analysis of FNR and ArcA regulons provides a molecular basis for understanding the observed metabolic changes. These oxygen-responsive transcription factors were activated under both CP and IP conditions, controlling over 80% of the metabolic flux in fluctuating oxygen environments^21,43^. Their cooperative regulation suppresses genes encoding key enzymes in the oxidative TCA cycle^21,44^ while activating anabolic pathways^45^, resulting in a rebalancing of catabolic and anabolic metabolism. This redirects carbon flux towards amino acid^46^ and fatty acid^47–49^ biosynthetic pathways that utilize acetyl-CoA^50^.

Changes in RONS levels and antioxidant enzyme activities further support the induction of adaptive stress responses. Under CP conditions, increased H_2_O_2_ and NO levels were observed, indicating oxidative stress. However, the concurrent elevation of CAT and GSH-POD activities, along with upregulation of G6PD, suggests the induction of oxidative eustress rather than distress^51^. Furthermore, H_2_O_2_ production under CP conditions remained below the critical sublethal threshold^52,53^ indicative of a balanced stress response. Under IP conditions, lower H_2_O_2_ levels were observed compared to CP conditions, suggesting a role for anoxic phases in maintaining redox balance. However, persistent NO concentrations may induce additional adaptive responses, as evidenced by the upregulation of enzymes regulated by NorR and NsrR, key transcription factors in the nitrosative stress response pathway^54^.

These findings have implications for both wastewater treatment and resource recovery. By strategically manipulating oxygen supply regimes, it may be possible to simultaneously enhance treatment efficiency, reduce excess sludge production, and promote the synthesis of valuable metabolites. This approach aligns with the growing emphasis on circular economy principles in wastewater management. However, several challenges remain to be addressed. The application of this aeration control strategy in wastewater treatment processes requires reliable oxygen control systems capable of accurately monitoring and adjusting oxygen levels.

Moreover, the long-term stability of these metabolic shifts in full-scale systems needs to be investigated to ensure the practical applicability of these findings.

## 5. Conclusions

Our study demonstrates the potential of strategic oxygen perturbations as a tool for optimizing biological processes in environmental engineering. By elucidating the complex interplay between oxygen supply regimes, treatment performance, and metabolite synthesis, we have uncovered new possibilities for enhancing resource recovery from wastewater treatment systems while maintaining efficient pollutant removal. These findings contribute to the broader goal of developing more sustainable and resource-efficient wastewater treatment technologies, aligning with global efforts towards circular economy approaches in water management.

## Abbreviations

Abbreviation Full Form

TCA=: Tricarboxylic Acid
WWTPs=: Wastewater Treatment Plants
DO=: Dissolved Oxygen
CEPT=: Chemical Enhanced Primary Treatment
HRAS=: High-Rate Activated Sludge
MLSS=: Mixed Liquor-Suspended Solid
CA=: Continuous Aeration
CP=: Continuous Perturbation
IP=: Intermittent Perturbation
ETC=: Electron Transport Chain
RONS=: Reactive Oxygen and Nitrogen Species
G6PD=: Glucose-6-Phosphate 1-Dehydrogenase
6PGD=: 6-Phosphogluconate Dehydrogenase

## Associated content

### Data availability statement

The mass spectrometry proteomics data have been deposited to ProteomeXchange Consortium^55^ (http://proteomecentral.proteomexchange.org) via the PRIDE partner repository with the dataset identifier PXD044490.

The mass spectrometry metabolomics data have been deposited to MetaboLights^56^ (https://www.ebi.ac.uk/metabolights/MTBLS8331) with the dataset identifier MTBLS8331.

### Supporting information

Detailed materials and methods, carbon and nitrogen conversion, biomass growth, enzymatic profiling analysis, reactive oxygen and nitrogen species, antioxidant system, and enzyme information (DOCX)

## Author information

### Notes

The authors declare no competing financial interest.

## Acknowledgements

We thank Martin Middleditch, George Guo, Saras Green, and Alastair Harris for their assistance with metaproteomics and metabolomics analysis. We also thank Watercare Services Limited for providing the activated sludge culture from the Māngere Wastewater Treatment Plant. The authors acknowledge assistance from the New Zealand eScience Infrastructure (NeSI) high-performance computing facilities and the Centre for eResearch at the University of Auckland.

