## Supplementary for "Resource Recovery from Wastewater By Directing Microbial Metabolism Toward Production of Value-added Biochemicals"

Figure S2. Changes in ammonia ($NH_4^+$-N), nitrite ($NO_2^-$-N), nitrate ($NO_3^-$-N), and COD concentrations in activated sludge systems during 48 h of exposure to the continuous perturbation condition. Error bars represent standard deviations (biological triplicates; n=3). . 10

Figure S3. Changes in ammonia ($NH_4^+$-N), nitrite ($NO_2^-$-N), nitrate ($NO_3^-$-N), and COD concentrations in activated sludge systems during 48 h of exposure to the intermittent perturbation condition. Error bars represent standard deviations (biological triplicates; n=3). . 11

### S1. SUPPLEMENTARY METHODS

#### S1.1. Bioreactor operation and performance

**Table S1.** Composition of the trace element solution

| Chemical | Concentration (g/L) |
|---|---|
| EDTA | 2.50 |
| $ZnSO_4 \cdot 7H_2O$ | 1.10 |
| $CoCl_2 \cdot 6H_2O$ | 0.80 |
| $MnCl_2 \cdot 4H_2O$ | 2.55 |
| $MgSO_4 \cdot 7H_2O$ | 20.0 |
| $CuSO_4 \cdot 5H_2O$ | 0.86 |
| $(NH_4)_6Mo_7O_{24} \cdot 4H_2O$ | 0.07 |
| $CaCl_2 \cdot 2H_2O$ | 2.75 |
| $FeSO_4 \cdot 7H_2O$ | 2.57 |

**Table S2.** Composition of the concentrated artificial wastewater

| Chemical | Concentration |
|---|---|
| Methanol (mL/L) | 26.94 |
| $NH_4Cl$ (g/L) | 24.46 (IP); 30.57 (CP or CA) |
| $KH_2PO_4$ (g/L) | 2.76 |
| $K_2HPO_4$ (g/L) | 2.76 |
| $NaHCO_3$ (g/L) | 76.80 |

#### S1.2. Water Quality and Biomass Analysis

Chemical oxygen demand (COD) was analysed using COD LR reagent vials (Hach, USA). $NH_4^+$-N concentration was measured with an AmVer Test N Tube Reagent Set (Hach, USA). $NO_2^-$-N and $NO_3^-$-N concentrations were measured using an ICS2100 Ion Chromatograph (ThermoFisher, USA). Total nitrogen (TN) was measured by a TN analyser (Shimadzu, Japan). The concentration of dissolved organic nitrogen was calculated by subtracting dissolved inorganic nitrogen from TN. MLSS and Mixed Liquor Volatile Suspended Solids (MLVSS) were analyzed using standard methods[2]. Gaseous nitrogen production was calculated by subtracting dissolved and biomass nitrogen from $NH_4^+$-N consumed by the system. Throughout the manuscript, the symbol ± denotes the standard deviation of the data.

For the determination of organic matter content in biomass, the 10 mL activated sludge solution was centrifuged at 6,113 x g for 10 min at 4°C. After discarding the supernatant, the pellet was resuspended in 0.85% NaCl solution, washed twice by re-centrifuging, discarding the supernatant and resuspending the pellet, and finally resuspended in 10 mL of 2% EDTA solution at 4°C for 3 hours. Subsequently, the sludge underwent three cycles of freeze/thaw treatment. Each cycle consisted of freezing the sample at -80°C for 30 minutes, followed by thawing at room temperature. After the freeze/thaw cycles, the sludge was sonicated for 20 cycles with 15 seconds of on/off intervals on ice. After centrifugation at 17,467 x g for 20 min, the supernatant was filtered through a 0.45 μm cellulose acetate filter for subsequent analysis.

Excitation-emission matrix (EEM) analysis was conducted using an Aqualog A-TEEM Spectrometer (Horiba, Japan) to determine the presence and concentration of fluorescent organic substances in sludge biomass. These substances include biosynthetic products such as protein-like and humic-like components, which are crucial indicators in biosynthesis. During EEM analysis, excitation wavelengths ranged from 240 to 600 nm (3 nm intervals), and emission wavelengths ranged from 212.70 to 622.21 nm (3.28 nm intervals). Fluorescence and absorbance data were modelled using PARAFAC by staRdom to determine relative amounts of organic matter in biomass[3]. Component properties were identified using the Openfluor database[4].

The protein concentration was measured by the RC DC™ Protein Assay Kit I (Bio-Rad, USA). The standard curve was established using a bovine γ-globulin standard. The contents of lipids and carbohydrates in the sludge were measured by the Soxhlet method[5] and the phenol-sulfuric acid method with glucose as the standard[6], respectively.

**S1.3. Metabolomics analysis**

The relative abundance of intracellular metabolites in activated sludge was determined using a GC/MS platform[7]. Three biological and two technical replicates were analyzed for each experimental condition. Sludge samples (10 mL) were centrifuged, the supernatant was discarded and the pellet was quenched in liquid nitrogen and stored at -80°C. To extract metabolites, 2.5 mL of cold methanol-water solution and 0.2 μmol of an internal standard, 2,3,3,3-d4-alanine, were added to the sludge pellets. The intracellular metabolites were released by three cycles of freeze-thawing followed by vigorous shaking, and the supernatant was collected by centrifugation at 22,707 x g for 15 min at -20°C. A further 2.5 ml of cold methanol-water solution was added to further extract the metabolites from the sludge pellet.

The metabolite extract was reconstituted in 400 μL of sodium hydroxide solution (1 M), supplemented with 68 μL of pyridine and 334 μL of methanol. A volume of 40 μL of methyl chloroformate was incorporated twice, and each addition was followed by a

30-second period of intense vortexing. Then 400 μL of chloroform was added and vortexed for a further 10 seconds. Subsequently, the blend was augmented with 800 μL of a sodium bicarbonate solution (50 mM), stirred, and subjected to centrifugation. The aqueous layer was disposed of, and any residual water in the chloroform phase was eliminated using anhydrous sodium sulfate.

The derivatized sample was analyzed with a gas chromatography–mass spectrometry (GC-MS) system (Agilent GC7890 coupled to an MSD597 unit) equipped with a ZB-1701 GC capillary column (30 m × 250 μm (id) × 0.15 μm film thickness) with a 5 m guard column (Phenomenex, Torrance, CA, USA). The obtained GC-MS chromatograms were deconvoluted by AMDIS software (NIST, Boulder, CO, USA). The mass spectra were compared to the in-house MS library to identify metabolites[7]. Data filtering was performed using the "MassOmics" R package[8], and metabolite abundance values were normalized with the internal standard. Baseline calibration was conducted by subtracting blank information. The relative abundance of intracellular metabolites over 48 hours was obtained and visually analyzed using MetaboAnalyst 5.0[9], and ComplexHeatmap[10].

**S1.4. Metaproteomics analysis**

Metaproteomic analysis was performed on sludge samples under various experimental conditions, using three biological replicates per condition. Sludge samples (5 mL) were pelleted via centrifugation at 6,113 x g for 5 min at 4°C, and washed twice with 0.85% NaCl solution. The pellets washed were flash-frozen in liquid nitrogen and stored at -80°C. Protein extraction was conducted by resuspending the pellet in lysis buffer (pH 8) containing 2% (w/v) SDS, 2 mM EDTA, 2 mM phenylmethylsulfonyl fluoride, 0.15% Triton X-100, 20 mM DTT, and 50 mM HEPES and sonicating for 20 cycles with 15 seconds of on/off intervals on ice. The lysate was collected following centrifugation at 17,467 x g for 30 min at 4°C.

The lysate was mixed with 20% trichloroacetic acid in acetone, vortexed, and incubated on ice. The mixture was centrifuged at 15,000 x g for 6 minutes at 4°C, and the supernatant was discarded. The pellet was washed with acetone, centrifuged, and washed again with 80% acetone in water. The protein pellet was air-dried and dissolved in 50 mM Tris buffer. Protein purification was further performed using SpeedBead carboxylate-modified E3 and E7 magnetic particles (Sera-Mag, USA). Protein concentration was quantified using the EZQ protein assay kit as per the manufacturer's instructions (Invitrogen, USA).

A 250 μL sample containing 150 μg protein extract was reduced by adding 5 mM DTT and incubation at 56°C for 15 minutes. Alkylation was carried out by adding 15 mM iodoacetamide and incubating in the dark for 30 minutes. The reaction was quenched by adding 15 mM cysteine. Protein digestion was performed by adding 1.5 μg trypsin and incubating at 37°C for 1 hour, followed by 18 hours at room temperature.

After digestion, the sample was diluted with 100 mM ammonium bicarbonate and centrifuged with a Vivaspin centrifugal concentrator (Sartorius, Germany) for 55 minutes at 28°C to intercept large molecules (>?kDa?) and obtain peptides filtered through. The sample was then diluted with 0.1% formic acid and peptides were extracted using Oasis Prime HLB 1cc (30 mg) solid-phase extraction. The peptide sample was loaded onto cartridges, which were subsequently washed with 1 mL of 0.1% formic acid. Peptides were eluted with 300 μL of 50% acetonitrile in 0.1% formic acid and concentrated to 15-20 μL using a speed vacuum.

A 10 μL aliquot of each peptide sample was analyzed by nano LC-MS/MS. The sample was desalted and separated using a NanoLC 400 UPLC system (Eksigent, USA). The TripleTOF 6600 Quadrupole-Time-of-Flight mass spectrometer (Sciex, USA) was used for mass spectrometry analysis.

The resulting data were searched against a database established by metagenomics results of the samples using MetaProteomeAnalyzer version 3.4[11]. The X-tandem was selected as the primary peptides and protein identification search engine[12]. The parameters were as follows: precursor and fragments tolerance set as 15 ppm; the cysteine alkylation set as iodoacetamide; the trypsin as the digestion enzyme with up to one mis-cleavage; a 1% false discovery rate was set as the filter of the final global protein groups. The resulting group file exported from X-tandem was converted to MetaProteomeAnalyzer for metaproteomics analysis and clustering. Information on enzymes regulated by regulons comes from RegulonDB v12.0[13].

#### S1.5. Reactive Oxygen and Nitrogen Species (RONS) and activity assay of ROS-scavenging enzymes

To assess the levels of RONS generated under different oxygen perturbation conditions, we conducted hourly measurements of the microbial RONS levels using assay kits at intervals from 0 to 8 hours, 24 to 36 hours, and at the 48th hour. Specifically, the relative concentrations of superoxide ($O_2^{\bullet -}$), hydrogen peroxide ($H_2O_2$), hydroxyl radical ($OH\cdot$), and nitric oxide (NO) were detected using CellROX™ Green Reagent (Invitrogen, USA), OxiVision™ Green Reagent (AAT Bioquest, USA), Invitrogen™ HPF Reagent (Invitrogen, USA), and DAF-FM DA Reagent (Abcam, UK), respectively. The fluorescence reaction system was set up in 96-well plates, comprising 50 μL of fluorescent probe working solution and 50 μL of the sample (with an additional 10 μL of 0.3% sodium azide in the $H_2O_2$ detection system). The reaction was incubated in the dark at 20°C for 60 minutes, and the fluorescence intensity was monitored at an Ex/Em wavelength of 490/525 nm.

The induction of antioxidant systems was ascertained by measuring superoxide dismutase (SOD), catalase (CAT), and glutathione peroxidase (GSH-POD), as follows. the 10 mL replicates of sludge sample was centrifuged at 5,000 g for 15 min at 4°C. The sludge pellet was washed with 50 mM Tris buffer twice and then disrupted by ultrasound set to 40% AMP and ten-second switch alternation for 10 min on ice (QSONICA, USA). The SOD, CAT and GSH-POD activities in the enzyme extracts were assayed according to the kit instructions using Superoxide Dismutase Activity Assay Kit (Abcam, UK), Catalase Activity Assay (Abcam, UK) and GSH-POD Activity Assay Kits (Abcam, UK), respectively.

### S2. Results

#### S2.1. The performance of carbon and nitrogen conversion under different aeration patterns

Three aeration conditions, CA, CP, and IP were compared based on their impact on wastewater treatment performance in terms of carbon and nitrogen conversion. The carbon sources provided were consumed in different aeration patterns, and the COD concentration in the dissolved state was maintained below 50 mg/L. Between the different treatments, there were greater differences in nitrogen conversion than carbon catabolism. CA and CP showed similar performance for nitrification (Figure S1, S2, S3). In contrast, IP, exhibited simultaneous nitrification and denitrification. Under CA, a stable aerobic environment, nitrogen conversion was dominated by aerobic nitrification from ammonia-oxidizing bacteria (AOB) and nitrite-oxidizing bacteria (NOB), resulting in 94.3±1.4% ammonia nitrogen accumulation as $NO_3^-$ and minimal gaseous nitrogen production.

Under the fluctuating aerobic conditions of CP, 81.7±2.1% of nitrogen was accumulated as $NO_3^-$-N in the system, and 9.5±1.3% conversion to gaseous nitrogen indicated the involvement of denitrification processes. There was no noticeable difference in ammonia conversion rates under the two aerobic experimental conditions (Figure S4). This indicates that the CP did not affect the capacity to convert ammonia nitrogen by AOB. Meanwhile, it might slightly increase the efficiency of nitrogen removal.

Under IP, the aerobic phase time was half of that under CP. This was found to reduce the ammonia conversion rate by AOBs (Figure S4). However, the alternating aerobic/anoxic conditions likely supported $NO_3^-$-N produced by the nitrification reaction to be used as a substrate for the denitrification reaction, and the anoxic phases provided time for denitrifying bacterial metabolism. This resulted in 71.3 ± 4.5% of consumed ammonia nitrogen being released in a gaseous form, achieving simultaneous nitrification and denitrification.

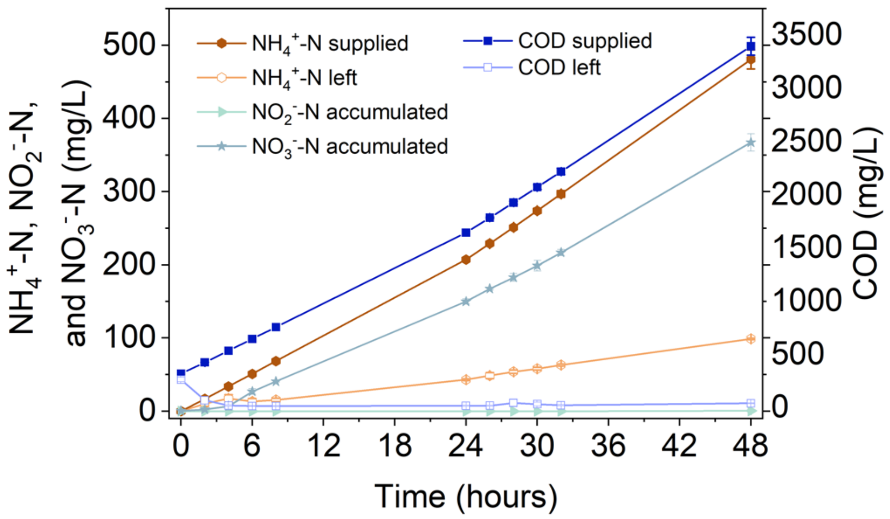

**Figure S1.** Changes in ammonia ($NH_4^+$-N), nitrite ($NO_2^-$-N), nitrate ($NO_3^-$-N), and COD concentrations in activated sludge systems during 48 h of exposure to the constant aeration condition. Error bars represent standard deviations (biological triplicates; n=3).

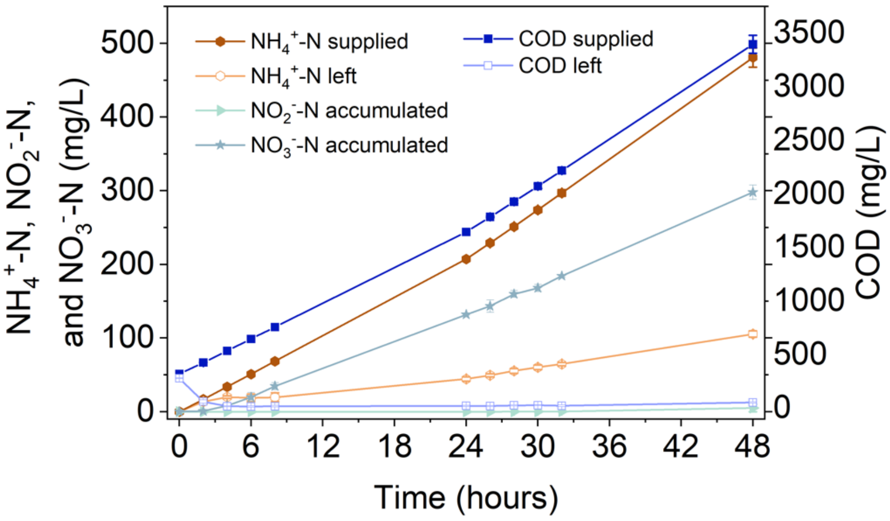

**Figure S2.** Changes in ammonia ($NH_4^+$-N), nitrite ($NO_2^-$-N), nitrate ($NO_3^-$-N), and COD concentrations in activated sludge systems during 48 h of exposure to the continuous perturbation condition. Error bars represent standard deviations (biological triplicates; n=3).

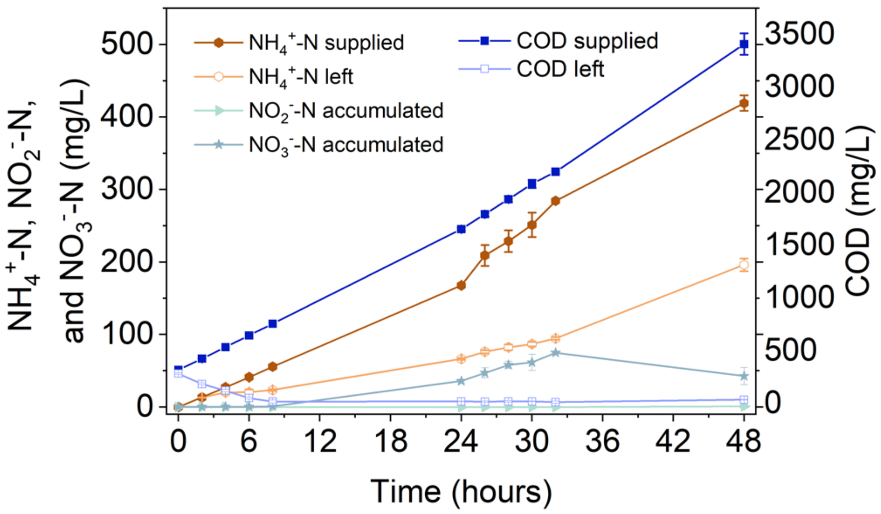

**Figure S3.** Changes in ammonia ($NH_4^+$-N), nitrite ($NO_2^-$-N), nitrate ($NO_3^-$-N), and COD concentrations in activated sludge systems during 48 h of exposure to the intermittent perturbation condition. Error bars represent standard deviations (biological triplicates; n=3).

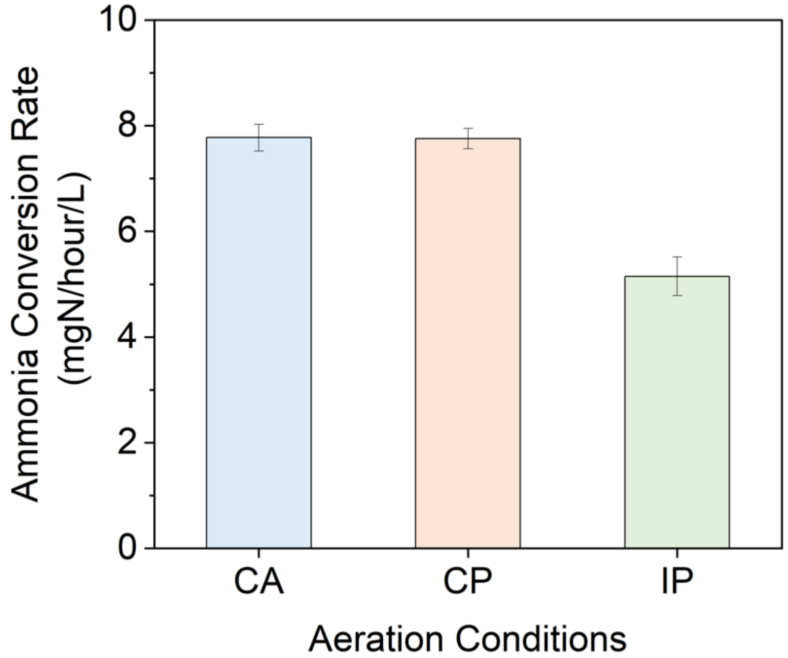

**Figure S4.** Ammonia conversion rate under different aeration conditions: constant aeration (CA), continuous perturbation (CP), and intermittent perturbation (IP) for 48 hours. Error bars represent standard deviations (biological triplicates; n=3).

#### S2.2. Analysis of biomass growth and organic composition under different aeration patterns

To evaluate biomass growth efficiencies, MLSS and MLVSS were analyzed both at the start and after 48 hours of aeration under these various conditions. The results show that perturbation strategies can differentially affect the efficiency of biomass synthesis in activated sludge systems. CP promoted more biomass synthesis compared to the CA condition, while biomass synthesis was the lowest under the IP condition (Figure S5, S6).

The relative abundance of organic substances was obtained through semi-quantitative analysis of lysed sludge samples using PARAFAC modelling for 3D-EEM data. Four fluorescent dissolved organic matters (fDOM) were identified (Figure S7, S8, S9), with a model fit of 99.3% R-squared. After 48 hours, the content of Component 1 increased more under CP (relative concentration, proportion; 0.0307±0.0013, 56.78±1.69%) than under CA (0.0259±0.0018, 53.99±1.58%) or IP (0.0208±0.0025, 49.18±2.63%) (Table S3). The other three components exhibited decreasing trends, with no significant differences (ANOVA Tukey test, $P>0.05$) under different aeration conditions. Based on the Openfluor database, these components were matched as protein-like (Component 1), humic-like (Components 2 and 4), and an unmatched substance (Component 3) (Table S4).

To evaluate changes in the organic content of biomass over 48 hours under varying aeration conditions, quantitative analysis focusing on protein, lipid, and carbohydrate content was performed. Notably, the CP conditions resulted in an increase in protein and lipid synthesis compared with CA conditions. Specifically, protein content in the sludge under CP conditions increased from 144.2±15.1 mg/g to 313.6±5.1 mg/g, while lipid content rose from 167.0±2.9 mg/g to 191.9±3.1 mg/g. In comparison, CA treatment resulted in protein and lipid contents of 293.0±15.0 mg/g and 186.0±2.1 mg/g, respectively. Statistical analysis based on ANOVA Tukey test revealed minor differences between CP and CA treatments, emphasizing the nuanced impacts of specific aeration strategies on biomass composition (Figure S10, S11). No statistical difference was observed in carbohydrate content between CP and CA (Figure S12).

Under IP conditions, the growth rate of activated sludge biomass was lower compared to CA and CP, with MLSS showing no significant changes post-treatment. However, MLVSS saw a rise by 6.3±0.7% (Figure S5, S6). Protein and lipid contents increased to 261.1±12.7 and 183.0±3.4 mg/g dry sludge, marking growth rates of 179.3±13.8% and 109.6±1.3% respectively (Figure S10, S11). While these increases were modest compared to those under CA and CP conditions, they still enhanced the valuable components within the biomass, underscoring the effectiveness of IP in sludge management.

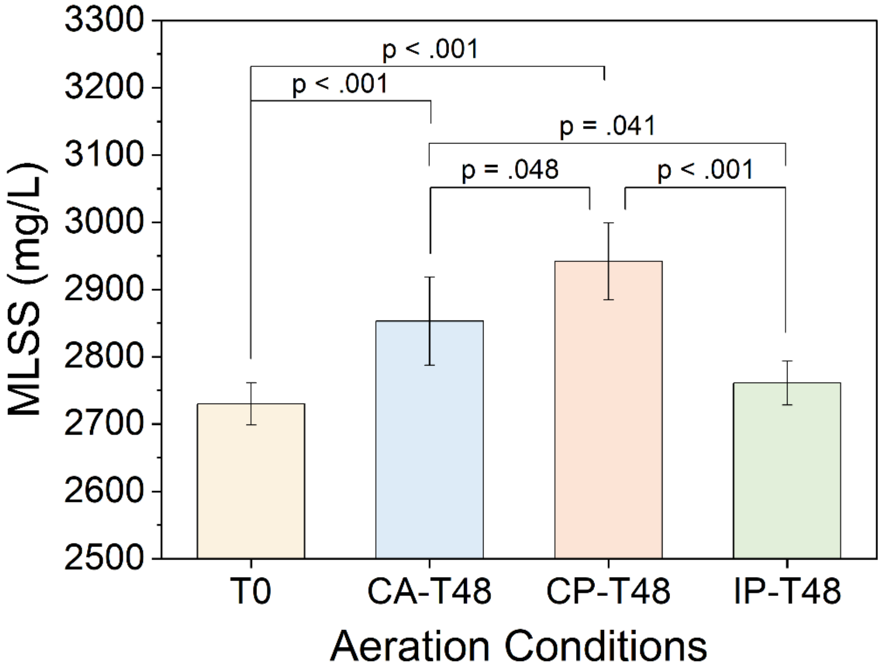

**Figure S5.** Mass of mixed liquor suspended solid (MLSS) in the activated sludge system at the beginning (T0) and after 48 hours of exposure to different aeration conditions: constant aeration (CA-T48), continuous perturbation (CP-T48), and intermittent perturbation (IP-T48), P values obtained from ANOVA Tukey test.

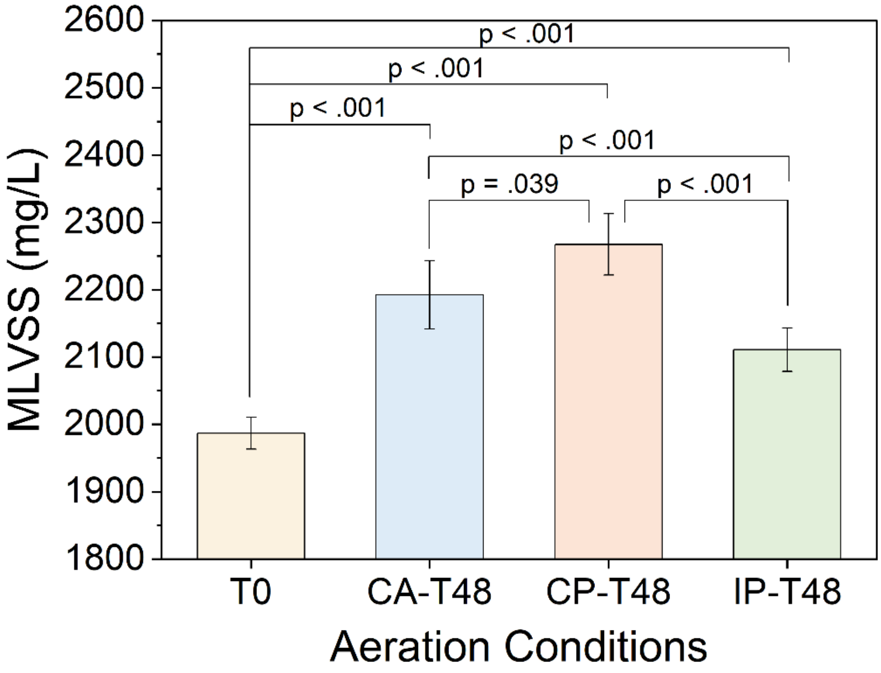

**Figure S6.** Mass of mixed liquor volatile suspended solid (MLVSS) in the activated sludge system at the beginning (T0) and after 48 hours of exposure to different aeration conditions:

constant aeration (CA-T48), continuous perturbation (CP-T48), and intermittent perturbation (IP-T48), P values obtained from ANOVA Tukey test.

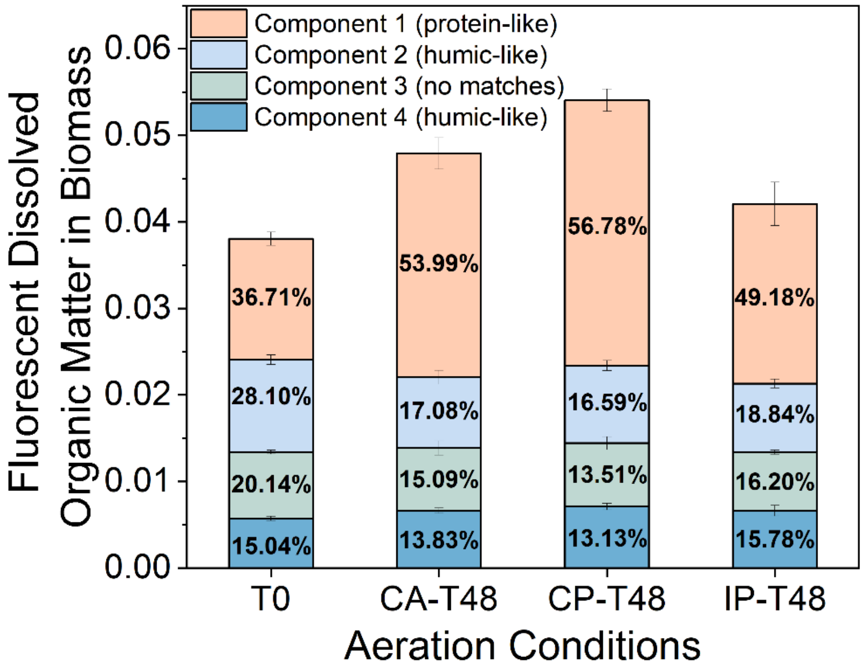

**Figure S7.** Fluorescent dissolved organic matter contents in biomass in the activated sludge system at the beginning (T0) and after 48 hours of exposure to different aeration conditions: constant aeration (CA-T48), continuous perturbation (CP-T48), and intermittent perturbation (IP-T48), based on PARAFAC analysis of EEM data. Error bars in the subfigures represent standard deviations of biological triplicates.

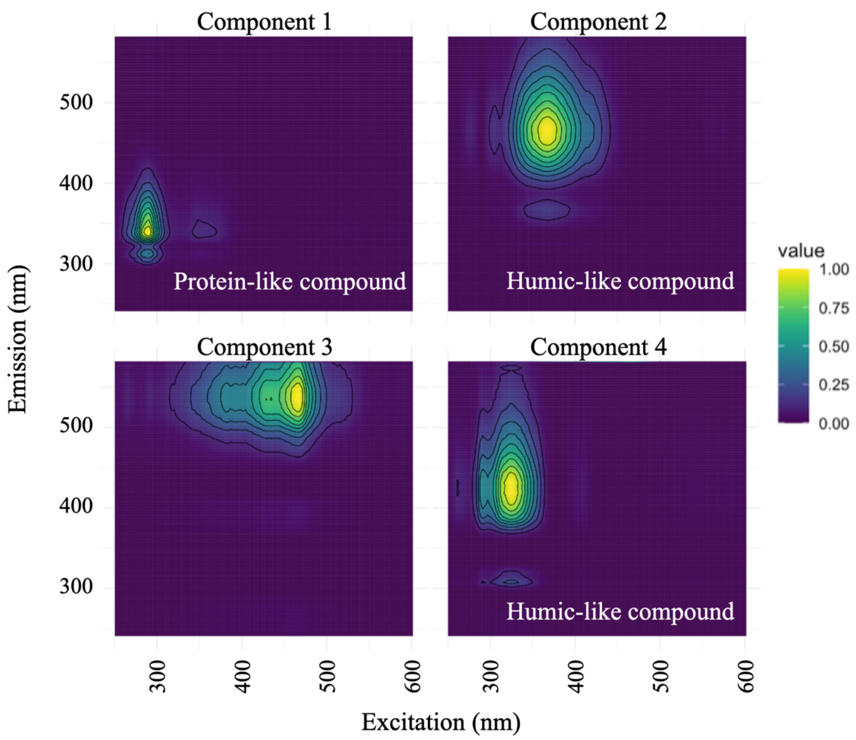

**Figure S8.** EEM fluorescence spectra generated by PARAFAC model analysis.

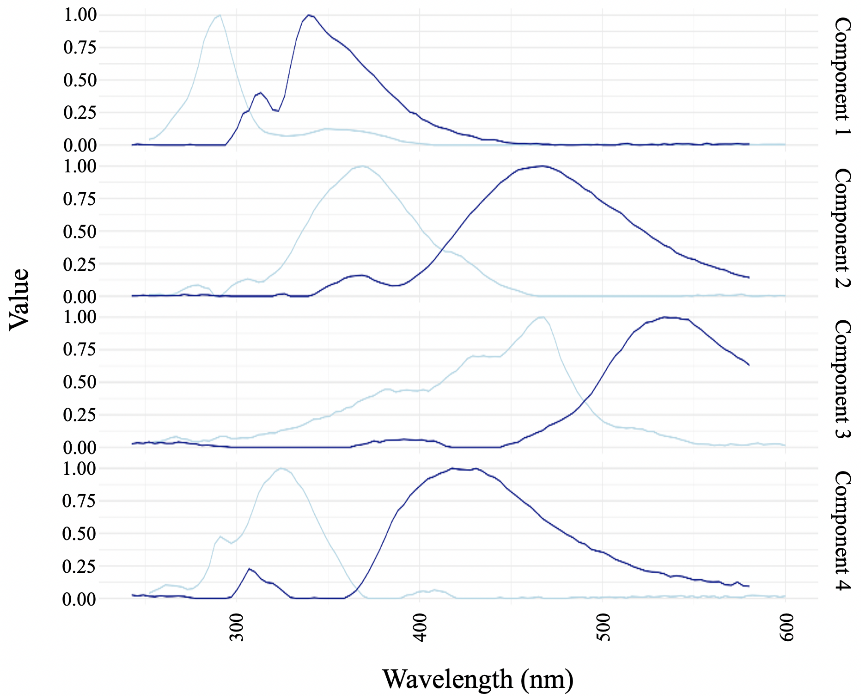

**Figure S9.** Excitation (light blue) and emission (dark blue) curves of EEM fluorescence spectra

**Table S3.** The relative concentrations of the four components in the activated sludge biomass samples from activated sludge systems after 48 hours of exposure to different aeration conditions: constant aeration (CA), continuous perturbation (CP), and intermittent perturbation (IP) (biological triplicates for T48 samples; n=3)

| Biomass samples | Component 1 | Component 2 | Component 3 | Component 4 |
|---|---|---|---|---|
| T0_1 | 0.0151 | 0.0109 | 0.0080 | 0.0061 |
| T0_2 | 0.0134 | 0.0099 | 0.0077 | 0.0056 |
| T0_3 | 0.0146 | 0.0113 | 0.0077 | 0.0060 |
| T0_4 | 0.0128 | 0.0100 | 0.0073 | 0.0053 |
| T0_5 | 0.0142 | 0.0111 | 0.0077 | 0.0057 |
| T0_6 | 0.0137 | 0.0109 | 0.0076 | 0.0056 |
| CA_T48_1 | 0.0279 | 0.0079 | 0.0077 | 0.0066 |
| CA_T48_2 | 0.0261 | 0.0093 | 0.0079 | 0.0070 |
| CA_T48_3 | 0.0235 | 0.0074 | 0.0061 | 0.0062 |
| CP_T48_1 | 0.0324 | 0.0087 | 0.0076 | 0.0070 |
| CP_T48_2 | 0.0304 | 0.0098 | 0.0081 | 0.0076 |
| CP_T48_3 | 0.0292 | 0.0084 | 0.0063 | 0.0067 |
| IP_T48_1 | 0.0172 | 0.0072 | 0.0066 | 0.0066 |
| IP_T48_2 | 0.0227 | 0.0081 | 0.0071 | 0.0058 |
| IP_T48_3 | 0.0224 | 0.0085 | 0.0066 | 0.0073 |

**Table S4.** The characteristics of the four components in the activated sludge biomass samples were matched by comparing the wavelengths of excitation and emission with the Openfluor database

| Component | Excitation maximum | Emission maximum | Assignment | References |
|---|---|---|---|---|
| Component 1 | 291 nm | 339 nm | Protein-like compound | 14 |
| Component 2 | 369 nm | 467 nm | Humic-like compound | 15 |
| Component 3 | 468 nm | 534 nm | No matches | |

| | | | | |
|---|---|---|---|---|
| Component 4 | 324 nm | 418 nm | Humic-like compound | 16 |

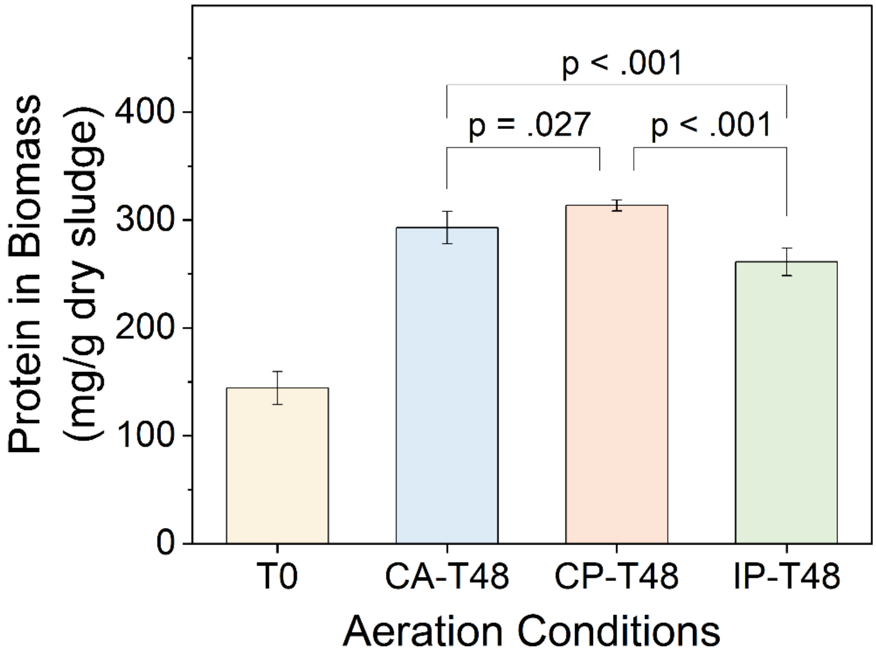

**Figure S10.** The protein content in biomass of the activated sludge system exposed to different aeration conditions: constant aeration (CA), continuous perturbation (CP), and intermittent perturbation (IP) from a starting point to 48 hours. No p-values were calculated for comparison with T0.

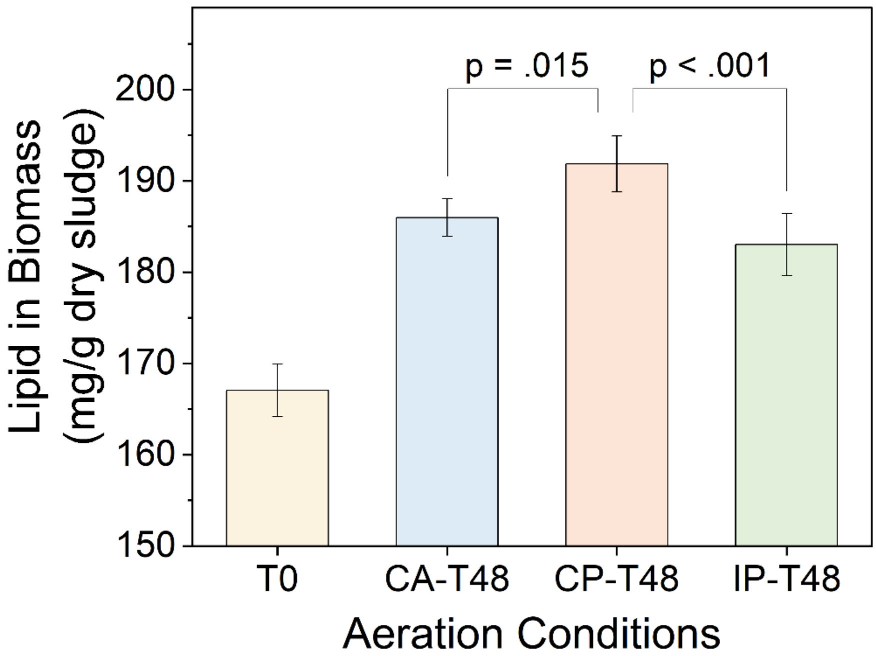

**Figure S11.** The lipid content in biomass of the activated sludge system exposed to different aeration conditions: constant aeration (CA), continuous perturbation (CP), and intermittent perturbation (IP) from a starting point to 48 hours. No p-values were calculated for comparison with T0.

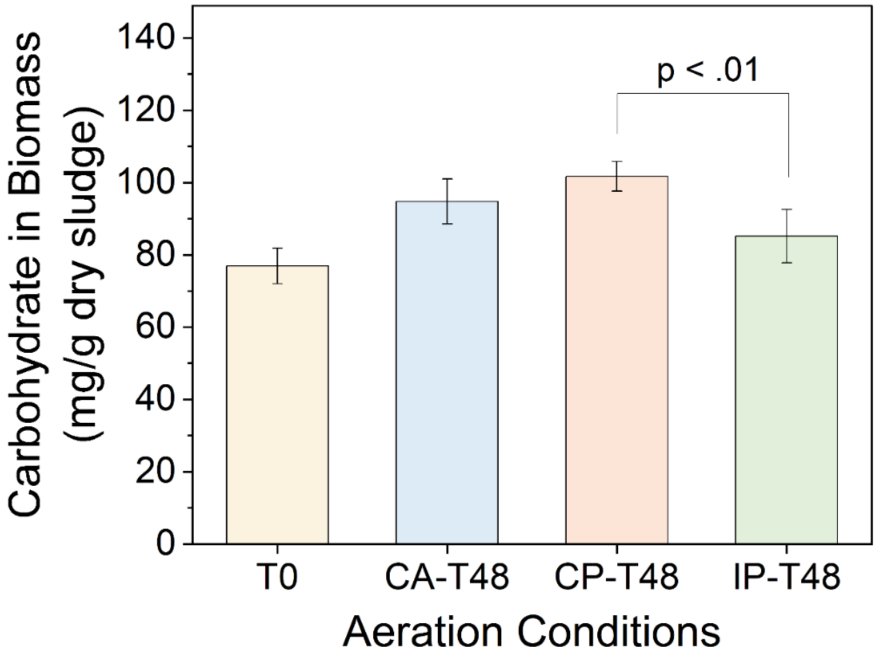

**Figure S12.** The carbohydrate content in biomass of the activated sludge system exposed to different aeration conditions: constant aeration (CA), continuous perturbation (CP), and intermittent perturbation (IP) from a starting point to 48 hours. No p-values were calculated for comparison with T0.

**S2.3. Enzymatic profiling analysis of microbial system as a whole**

The impact of oxygen perturbation strategies on the microbial communities within activated sludge systems was analyzed on enzyme expression levels, revealing potential shifts in systemic metabolic characteristics. Our comprehensive enzymatic spectrum analysis aimed to interpret these metabolic trait shifts under oxygen perturbations (Figure S13, S14), thereby offering insights into the microbial strategic response to varied oxygen environments.

Under CP (Table S5), there was a marked upregulation of enzymes such as DnaK suppressor protein and modulator of FtsH protease, highlighting an intensified microbial stress response induced by oxygen fluctuations. The differential expression of acetyl-CoA carboxylase and fructose-1,6-bisphosphatase II underscores a redirection of carbon utilization pathways, likely optimized for dynamic oxygen levels presented by the CP strategy. Furthermore, an increase in cytochrome c oxidase cbb3-type abundance under CP conditions points to an adaptively enhanced efficiency in oxygen utilization. Concurrently, the varying abundance of aminoacyl-tRNA synthetases indicates that protein synthesis efficiency is affected.

A different trend in enzyme levels was observed under IP (Table S6). Enzymes such as the two-component system NtrC family response regulator and phosphoglycerol transferase exhibited reduced abundance, implying a downregulation of pathways implicated in stress response and membrane biosynthesis. Conversely, enzymes such as ubiquinol-cytochrome c reductase cytochrome c1 subunit and microcin C transport system substrate-binding protein displayed increased abundance, indicative of an upregulated capacity for respiratory energy production and nutrient uptake mechanisms. This trend was supported by the notable upsurge in modulator of FtsH protease and gluconate:H+ symporter, emphasizing a bolstered effort to maintain protein homeostasis and energy balance. The pronounced upregulation of enzymes like fumarate hydratase and sulfate transport system substrate-binding protein also illustrates a tendency toward augmented pathways for energy conservation and sulfur metabolism under IP conditions.

Distinct from the significant influence of aeration patterns on assimilatory pathways (Figure S15, described in results 3.1), enzymes active in dissimilatory pathways exhibited less variability across different aeration conditions. The expression levels of enzymes such as 5,6,7,8-tetrahydromethanopterin hydro-lyase (fae), methylene-tetrahydromethanopterin dehydrogenase (mtdB), methenyltetrahydromethanopterin cyclohydrolase (mch), formylmethanofuran-tetrahydromethanopterin N-formyltransferase (ftr), and formate dehydrogenase (fdoG) are shown in Figure S16. This suggests a stability in the dissimilatory processes that could be essential for maintaining basic cellular functions under variable oxygen conditions.

These findings cumulatively portray a metabolic adaptation landscape where microbial consortia demonstrate a wide-ranging response, optimizing energy generation via carbon metabolism and respiratory chain activities, and modulating anabolic strategies to equilibrate the effects of oxygen perturbations.

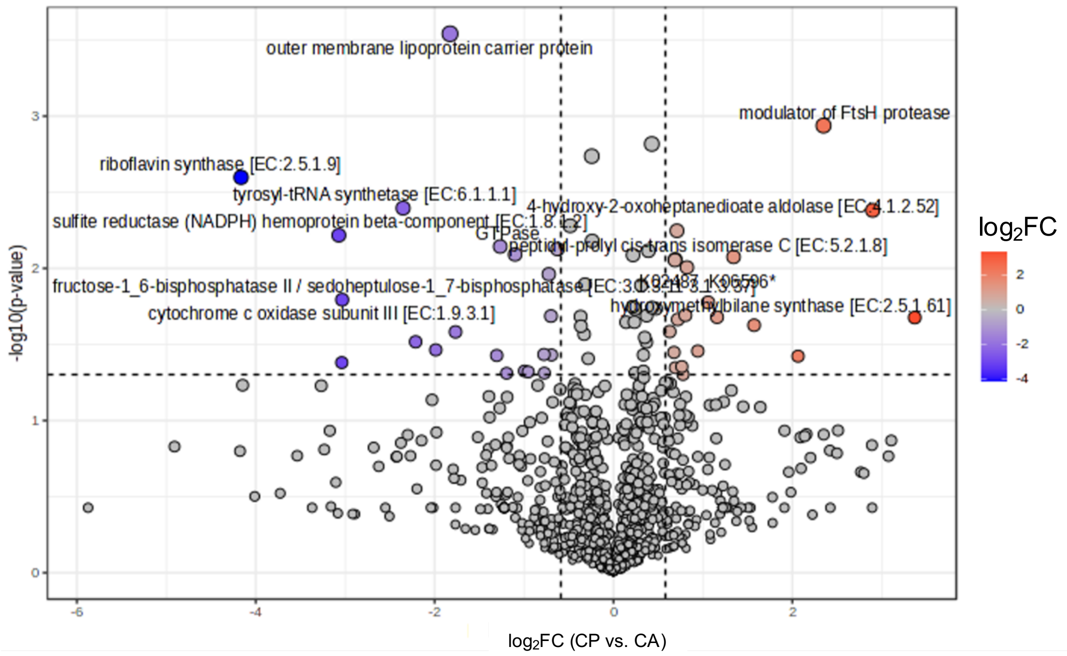

**Figure S13.** Volcano plot (Continuous Perturbation vs. Constant Aeration) representing the differential abundance of proteins in the activated sludge system after CP and CA treatments for 48 hours (T test, P<0.05, $|Log_2FC|>0.58$).

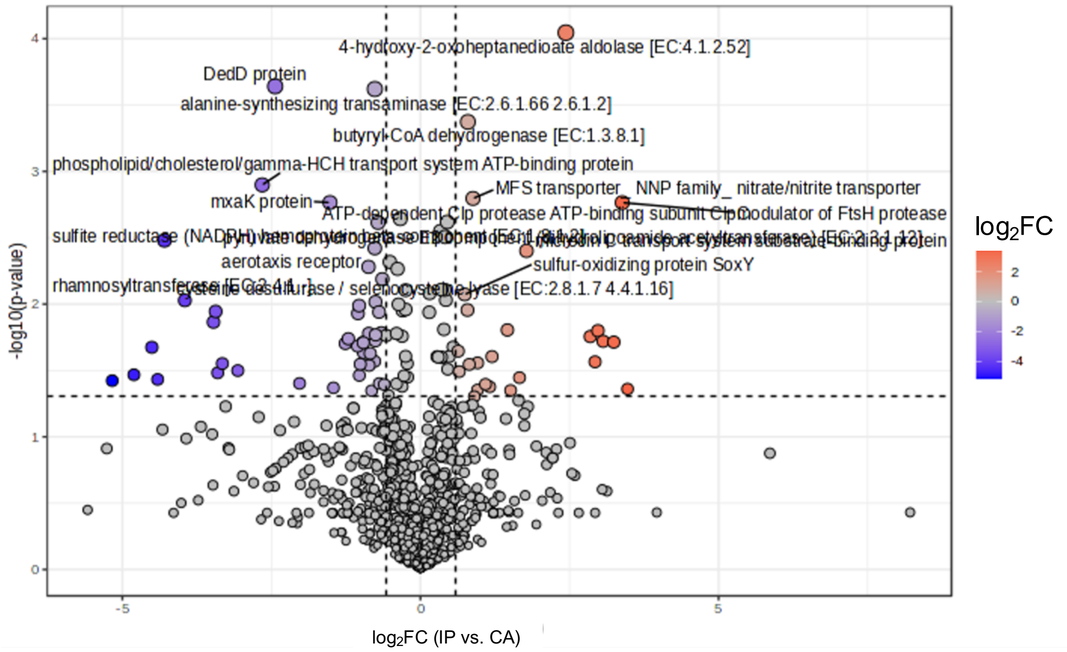

**Figure S14.** Volcano plot (Intermittent Perturbation vs. Constant Aeration) representing the differential abundance of proteins in the activated sludge system after treatment with IP and CA conditions for 48 hours (T test, P<0.05, $|Log_2FC|>0.58$).

**Table S5.** Proteins in the activated sludge microbial system under Continuous Perturbation with significant difference in abundance compared to Constant Aeration (T test, P<0.05, $|Log_2FC|>0.58$).

| Differentially abundant proteins | $Log_2FC$ | p-value |
|---|---|---|
| hydroxymethylbilane synthase [EC:2.5.1.61] | 3.369 | 0.021 |
| 4-hydroxy-2-oxoheptanedioate aldolase [EC:4.1.2.52] | 2.898 | 0.004 |
| modulator of FtsH protease | 2.349 | 0.001 |
| DnaK suppressor protein | 2.064 | 0.038 |
| Xaa-Pro aminopeptidase [EC:3.4.11.9] | 1.572 | 0.024 |
| energy-dependent translational throttle protein EttA | 1.343 | 0.008 |
| competence protein ComEA | 1.158 | 0.021 |
| K02487_K06596* | 1.058 | 0.017 |
| enolase [EC:4.2.1.11] | 0.943 | 0.035 |
| methanol dehydrogenase (cytochrome c) subunit 2 [EC:1.1.2.7] | 0.821 | 0.010 |
| high-affinity iron transporter | 0.804 | 0.020 |
| D-alanyl-D-alanine carboxypeptidase [EC:3.4.16.4] | 0.778 | 0.050 |
| lysyl-tRNA synthetase_ class II [EC:6.1.1.6] | 0.764 | 0.044 |
| cytochrome c oxidase cbb3-type subunit II | 0.719 | 0.022 |
| peptidyl-prolyl cis-trans isomerase C [EC:5.2.1.8] | 0.709 | 0.006 |
| heterogeneous nuclear ribonucleoprotein G | 0.692 | 0.045 |
| butyryl-CoA dehydrogenase [EC:1.3.8.1] | 0.690 | 0.009 |
| outer membrane protein_ adhesin transport system | 0.687 | 0.009 |
| acetyl-CoA carboxylase carboxyl transferase subunit alpha [EC:6.4.1.2 2.1.3.15] | 0.676 | 0.036 |
| cytochrome c oxidase cbb3-type subunit III | 0.628 | 0.026 |
| CBS domain-containing protein | -0.627 | 0.007 |
| phosphate transport system substrate-binding protein | -0.693 | 0.037 |
| HPr kinase/phosphorylase [EC:2.7.11.- 2.7.4.-] | -0.699 | 0.021 |
| fructose-1_6-bisphosphatase II / sedoheptulose-1_7-bisphosphatase [EC:3.1.3.11 3.1.3.37] | -0.723 | 0.011 |
| phosphoglucosamine mutase [EC:5.4.2.10] | -0.773 | 0.049 |
| ribose-phosphate pyrophosphokinase [EC:2.7.6.1] | -0.778 | 0.037 |
| two-component system_ OmpR family_ osmolarity sensor histidine kinase EnvZ [EC:2.7.13.3] | -0.953 | 0.048 |
| 3-phenylpropionate/trans-cinnamate dioxygenase ferredoxin reductase component [EC:1.18.1.3] | -0.995 | 0.047 |
| putative (di)nucleoside polyphosphate hydrolase [EC:3.6.1.-] | -1.101 | 0.008 |
| molybdopterin molybdotransferase [EC:2.10.1.1] | -1.195 | 0.049 |
| GTPase | -1.267 | 0.007 |
| putative iron-regulated protein | -1.306 | 0.037 |
| iron-sulfur cluster insertion protein | -1.766 | 0.026 |

| | | |
|---|---|---|
| outer membrane lipoprotein carrier protein | -1.828 | 0.000 |
| methionyl-tRNA synthetase [EC:6.1.1.10] | -1.987 | 0.034 |
| two-component system_ NtrC family_ sensor histidine kinase PilS [EC:2.7.13.3] | -2.213 | 0.030 |
| tyrosyl-tRNA synthetase [EC:6.1.1.1] | -2.353 | 0.004 |
| cytochrome c oxidase subunit III [EC:1.9.3.1] | -3.034 | 0.016 |
| K00329_K00356* | -3.038 | 0.042 |
| sulfite reductase (NADPH) hemoprotein beta-component [EC:1.8.1.2] | -3.071 | 0.006 |
| riboflavin synthase [EC:2.5.1.9] | -4.165 | 0.003 |

**Table S6.** Proteins in the activated sludge microbial system under Intermittent Perturbation with significant difference in abundance compared to Constant Aeration (T test, $P<0.05$, $|Log_2FC|>0.58$).

| **Differentially abundant proteins** | **$Log_2FC$** | **p-value** |
|---|---|---|
| gluconate:H+ symporter_ GntP family | 3.474 | 0.044 |
| modulator of FtsH protease | 3.385 | 0.002 |
| xanthine dehydrogenase molybdenum-binding subunit [EC:1.17.1.4] | 3.243 | 0.019 |
| threonine synthase [EC:4.2.3.1] | 3.064 | 0.019 |
| sulfate transport system substrate-binding protein | 2.978 | 0.016 |
| L-lactate dehydrogenase complex protein LldG | 2.928 | 0.027 |
| pyridoxal 5_-phosphate synthase pdxS subunit [EC:4.3.3.6] | 2.849 | 0.018 |
| 4-hydroxy-2-oxoheptanedioate aldolase [EC:4.1.2.52] | 2.439 | 0.000 |
| microcin C transport system substrate-binding protein | 1.779 | 0.004 |
| ubiquinol-cytochrome c reductase cytochrome c1 subunit | 1.661 | 0.036 |
| pyrroloquinoline-quinone synthase [EC:1.3.3.11] | 1.513 | 0.045 |
| acyl phosphate:glycerol-3-phosphate acyltransferase [EC:2.3.1.275] | 1.458 | 0.016 |
| glutamate/aspartate transport system ATP-binding protein [EC:7.4.2.1] | 1.200 | 0.025 |
| energy-dependent translational throttle protein EttA | 1.162 | 0.042 |
| fumarate hydratase_ class I [EC:4.2.1.2] | 1.091 | 0.040 |
| acyl-CoA dehydrogenase [EC:1.3.99.-] | 0.958 | 0.028 |
| 3-hydroxybutyryl-CoA dehydrogenase [EC:1.1.1.157] | 0.957 | 0.044 |
| CRISPR-associated protein Csd2 | 0.903 | 0.050 |
| MFS transporter_ NNP family_ nitrate/nitrite transporter | 0.881 | 0.002 |
| TRAP-type transport system periplasmic protein | 0.819 | 0.028 |
| butyryl-CoA dehydrogenase [EC:1.3.8.1] | 0.795 | 0.000 |
| putrescine transport system ATP-binding protein | 0.789 | 0.011 |
| sulfur-oxidizing protein SoxY | 0.742 | 0.008 |

| | | |
|---|---|---|
| phosphomethylpyrimidine synthase [EC:4.1.99.17] | 0.650 | 0.032 |
| universal stress protein A | 0.633 | 0.023 |
| type VI secretion system secreted protein Hcp | -0.606 | 0.040 |
| cysteine desulfurase / selenocysteine lyase [EC:2.8.1.7 4.4.1.16] | -0.639 | 0.006 |
| HPr kinase/phosphorylase [EC:2.7.11.- 2.7.4.-] | -0.656 | 0.016 |
| selenium-binding protein 1 | -0.690 | 0.040 |
| glutamate N-acetyltransferase / amino-acid N-acetyltransferase [EC:2.3.1.35 2.3.1.1] | -0.695 | 0.011 |
| ATP-dependent Clp protease ATP-binding subunit ClpC | -0.712 | 0.002 |
| preprotein translocase subunit SecF | -0.752 | 0.017 |
| acyl-CoA dehydrogenase [EC:1.3.8.7] | -0.755 | 0.010 |
| GTP cyclohydrolase IB [EC:3.5.4.16] | -0.756 | 0.019 |
| alanine-synthesizing transaminase [EC:2.6.1.66 2.6.1.2] | -0.765 | 0.000 |
| pyruvate dehydrogenase E2 component (dihydrolipoamide acetyltransferase) [EC:2.3.1.12] | -0.766 | 0.004 |
| 1_4-alpha-glucan branching enzyme [EC:2.4.1.18] | -0.771 | 0.027 |
| glutamate synthase (NADPH) large chain [EC:1.4.1.13] | -0.819 | 0.045 |
| immune inhibitor A [EC:3.4.24.-] | -0.843 | 0.024 |
| cell division protein FtsA | -0.862 | 0.029 |
| LPS-assembly protein | -0.867 | 0.017 |
| aerotaxis receptor | -0.871 | 0.005 |
| 6_7-dimethyl-8-ribityllumazine synthase [EC:2.5.1.78] | -0.955 | 0.023 |
| putative redox protein | -0.957 | 0.020 |
| glutamate dehydrogenase (NADP+) [EC:1.4.1.4] | -0.965 | 0.019 |
| phosphoribosylaminoimidazolecarboxamide formyltransferase / IMP cyclohydrolase [EC:2.1.2.3 3.5.4.10] | -1.010 | 0.028 |
| 5-methyltetrahydrofolate--homocysteine methyltransferase [EC:2.1.1.13] | -1.025 | 0.034 |
| GTPase | -1.029 | 0.010 |
| adenylylsulfate reductase_ subunit A [EC:1.8.99.2] | -1.048 | 0.012 |
| putative colanic acid biosynthesis UDP-glucose lipid carrier transferase | -1.054 | 0.021 |
| uridylate kinase [EC:2.7.4.22] | -1.210 | 0.018 |
| putative ABC transport system ATP-binding protein | -1.259 | 0.020 |
| molecular chaperone Hsp33 | -1.460 | 0.043 |
| mxaK protein | -1.518 | 0.002 |
| guanylate kinase [EC:2.7.4.8] | -2.028 | 0.040 |
| DedD protein | -2.438 | 0.000 |
| phospholipid/cholesterol/gamma-HCH transport system ATP-binding protein | -2.654 | 0.001 |
| sulfite reductase (NADPH) flavoprotein alpha-component [EC:1.8.1.2] | -3.063 | 0.032 |
| urocanate hydratase [EC:4.2.1.49] | -3.323 | 0.028 |
| nitrite reductase (NADH) large subunit [EC:1.7.1.15] | -3.401 | 0.033 |

| | | |
|---|---|---|
| two-component system_ NtrC family_ sensor histidine kinase PilS [EC:2.7.13.3] | -3.435 | 0.011 |
| methionyl-tRNA synthetase [EC:6.1.1.10] | -3.472 | 0.014 |
| rhamnosyltransferase [EC:2.4.1.-] | -3.952 | 0.009 |
| sulfite reductase (NADPH) hemoprotein beta-component [EC:1.8.1.2] | -4.294 | 0.003 |
| beta-ureidopropionase [EC:3.5.1.6] | -4.409 | 0.037 |
| peptidyl-prolyl cis-trans isomerase A (cyclophilin A) [EC:5.2.1.8] | -4.505 | 0.021 |
| phosphoglycerol transferase [EC:2.7.8.20] | -4.807 | 0.034 |
| two-component system_ NtrC family_ response regulator | -5.170 | 0.038 |

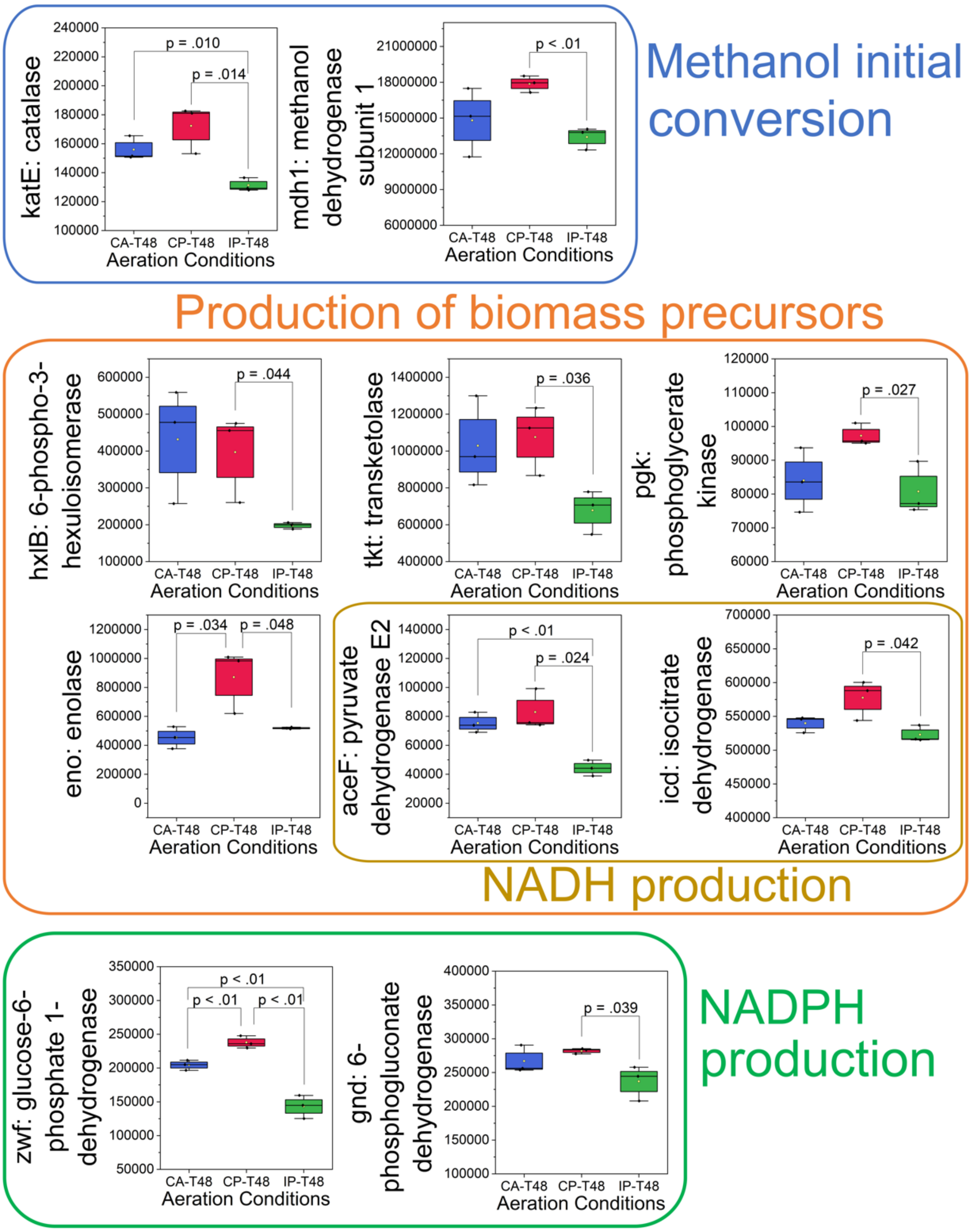

**Figure S15.** The enzymes with significantly different abundances in the assimilation pathways after 48 hours of treatment between two perturbations: constant aeration (CA-T48), continuous perturbation (CP-T48), and intermittent perturbation (IP-T48). P values were obtained from T-tests. The error bars represent standard deviations (biological triplicates; n=3).

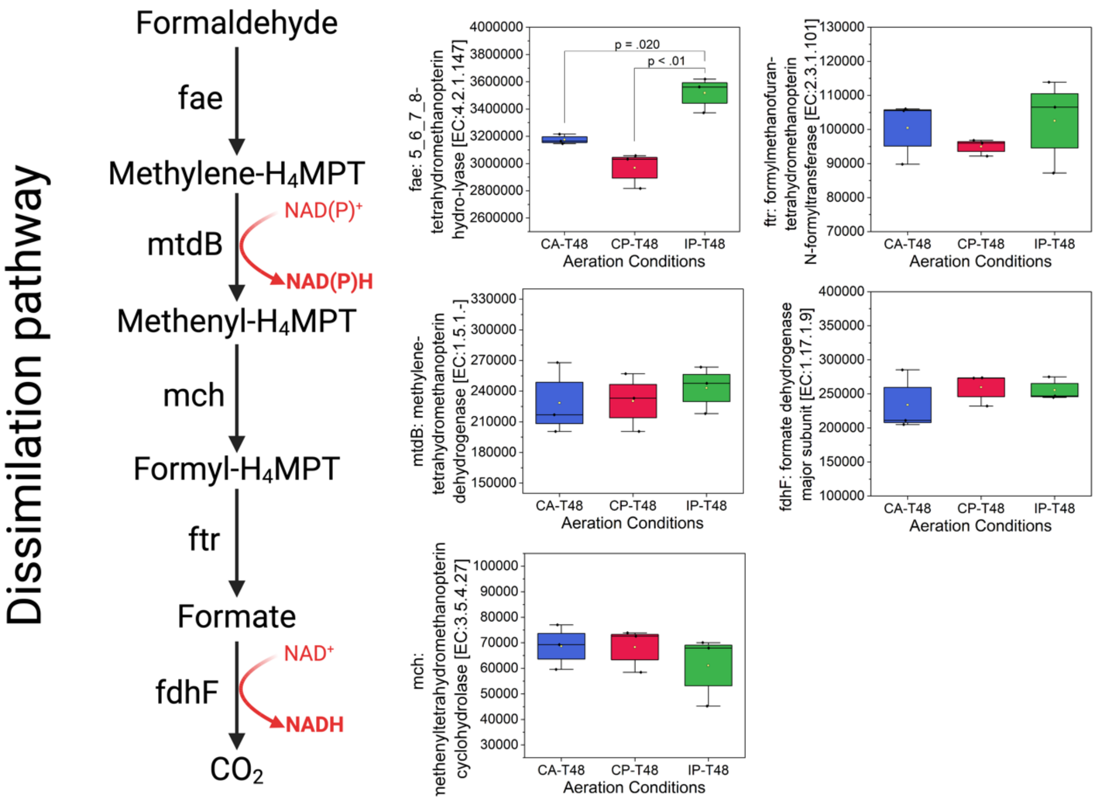

**Figure S16.** The abundance of main enzymes on the dissimilation pathway after 48 hours of treatment under different aeration conditions: constant aeration (CA-T48), continuous perturbation (CP-T48), and intermittent perturbation (IP-T48). P values were obtained from T-tests. The error bars represent standard deviations (biological triplicates; n=3).

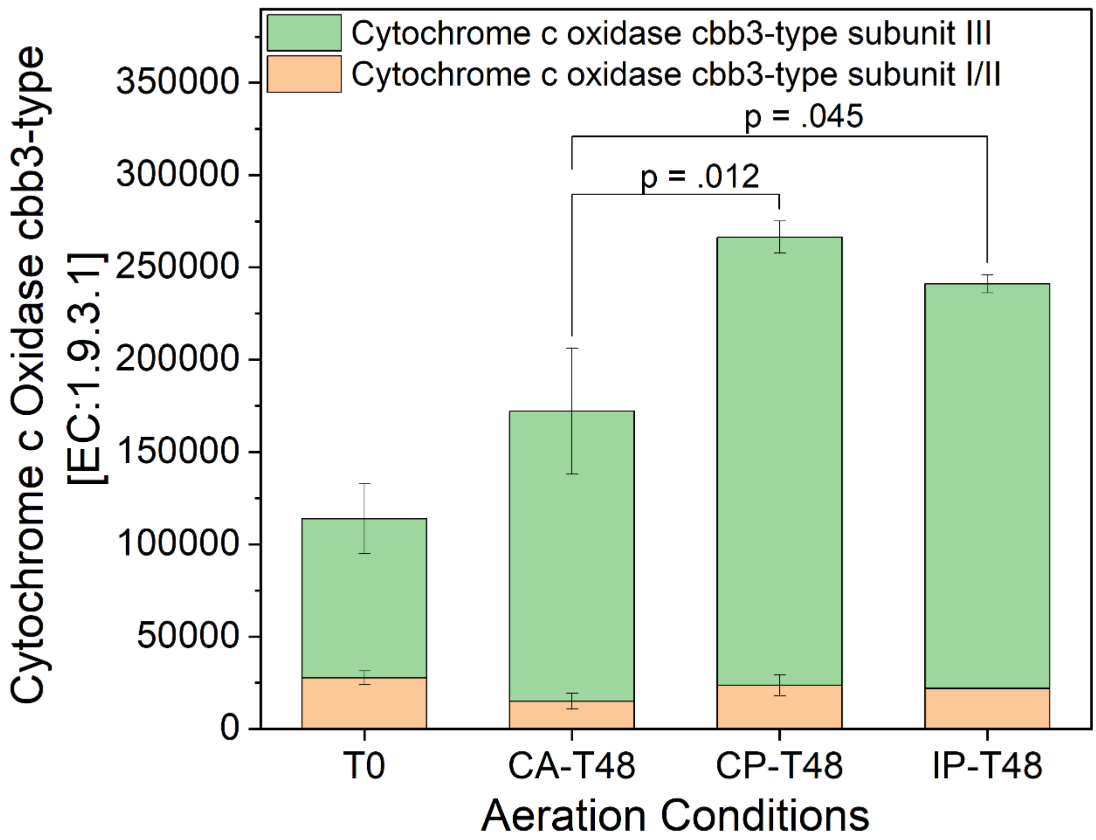

**Figure S17.** The relative concentration of cytochrome cbb3-type oxidase in oxidative phosphorylation under different aeration conditions: constant aeration (CA-T48), continuous perturbation (CP-T48), and intermittent perturbation (IP-T48). P values were obtained from ANOVA Tukey tests. The error bars represent standard deviations (biological triplicates; n=3). No p-values were calculated for comparison with T0.

### S2.4. Metabolite profile analysis

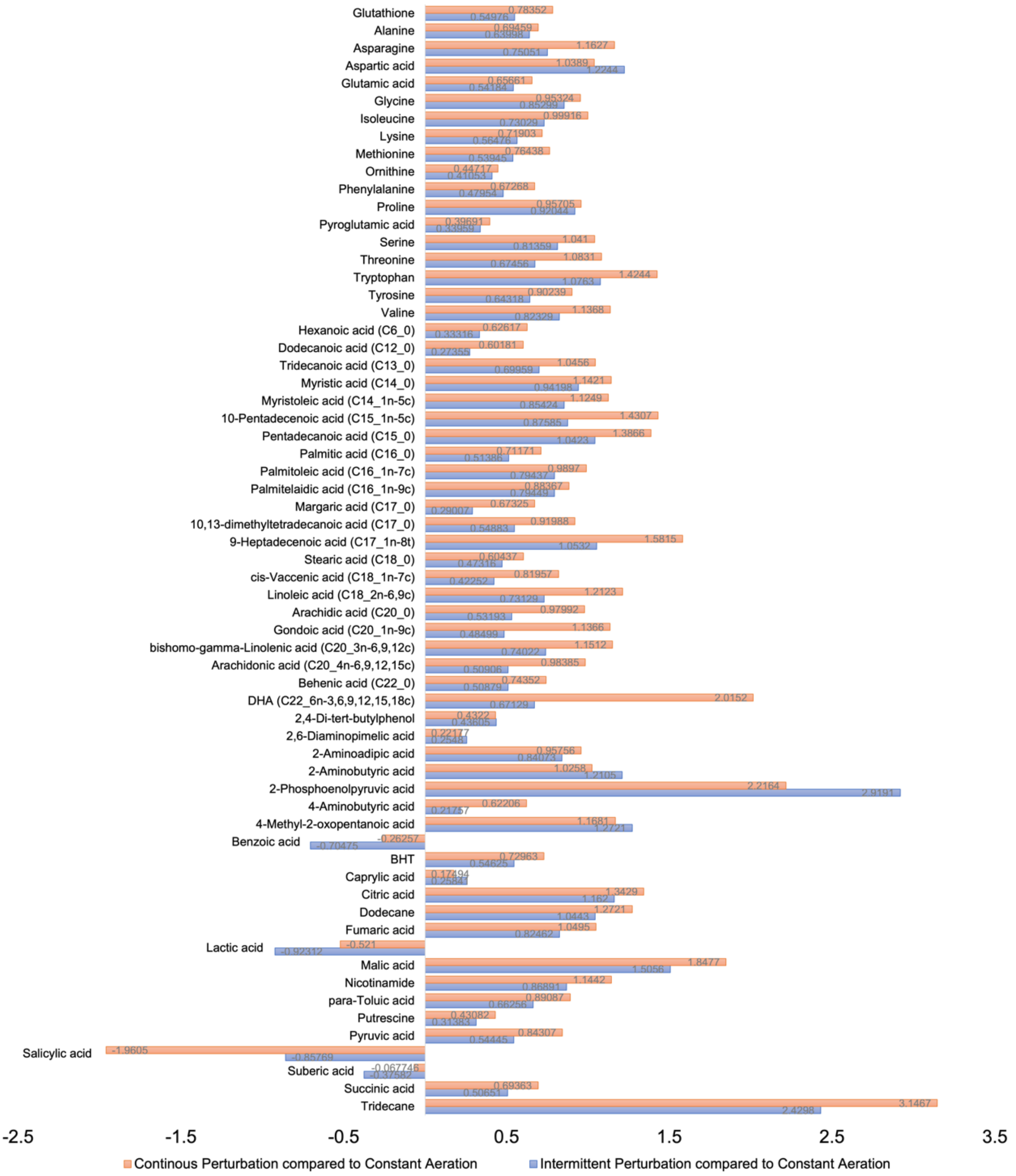

**Figure S18.** Log2 fold change in relative abundance of metabolites in activated sludge exposed to oxygen perturbations compared with constant aeration (technical duplication results for biological triplicates; n=6).

**S2.5. Reactive oxygen and nitrogen species and antioxidant system analysis**

During the 48-hour bioreactor operation, we observed the levels of $O_2^{\bullet -}$ (Figures S19), $H_2O_2$ (Figures S20), OH• (Figures S21), and NO (Figures S22) in the activated sludge systems under different aeration conditions. Among them, $H_2O_2$ and NO concentrations showed distinct differences under different oxygen perturbation treatments. Specifically, under CP conditions, $H_2O_2$ concentrations ranged between 32-56 μM, while under CA conditions, they stabilized between 21-45 μM. IP conditions resulted in $H_2O_2$ concentrations remaining below 20 μM (Figure 20). For NO, IP conditions resulted in the highest concentrations, ranging between 16 and 24 μM, followed by CP conditions (11 to 18 μM) and CA conditions (9 to 13 μM) (Figure 22).

In response to oxidative stress, microbes convert ROS through enzymes in the antioxidant system (Figure S23). Before and after the bioreactor was run for 48 hours, we measured the enzymatic activity of SOD (Figure S24), CAT (Figure S25), and GSH-POD (Figure S26) in activated sludge systems under different aeration conditions. After 48 hours of treatment, the difference in SOD activity in the antioxidant system was not significant. However, the activities of CAT and GSH-POD, which can convert $H_2O_2$, showed that they were specifically affected by aeration patterns. CAT activity appeared to correlate with $H_2O_2$ concentration, with a clear increase under CA and CP, but remained unchanged for IP. Microorganisms also use the GSH system to convert $H_2O_2$ into water. This process requires the reducing power of NADPH for the regeneration of glutathione that gets consumed during the process of $H_2O_2$ decrease. We found that CA and CP conditions led to significantly increased GSH-POD activity relative to IP treatment, which might be due the higher endogenous $H_2O_2$ concentrations under these conditions.

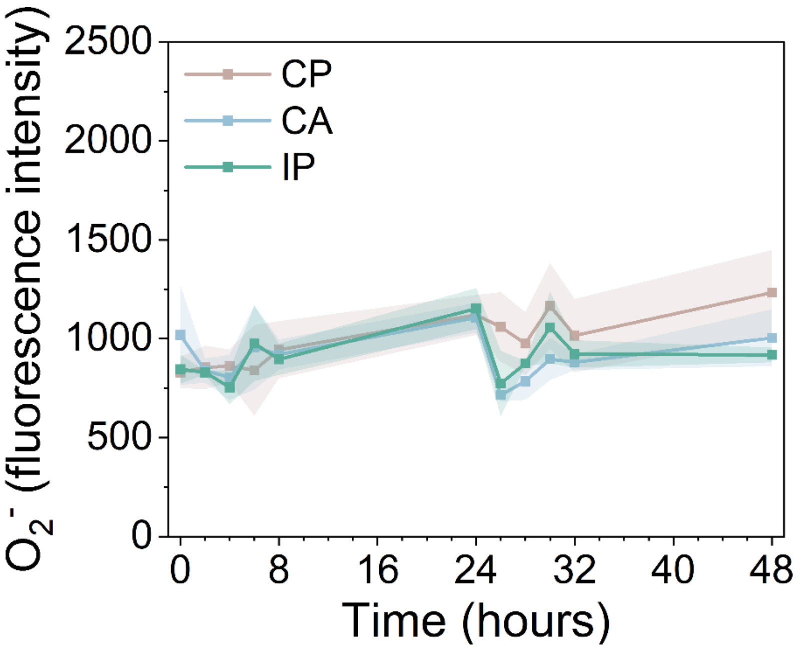

**Figure S19.** The fluorescence intensities of $O_2^{\bullet-}$ under different aeration conditions: constant aeration (CA), continuous perturbation (CP), and intermittent perturbation (IP). The error envelope represents standard deviations (technical duplication results for biological triplicates; n=6).

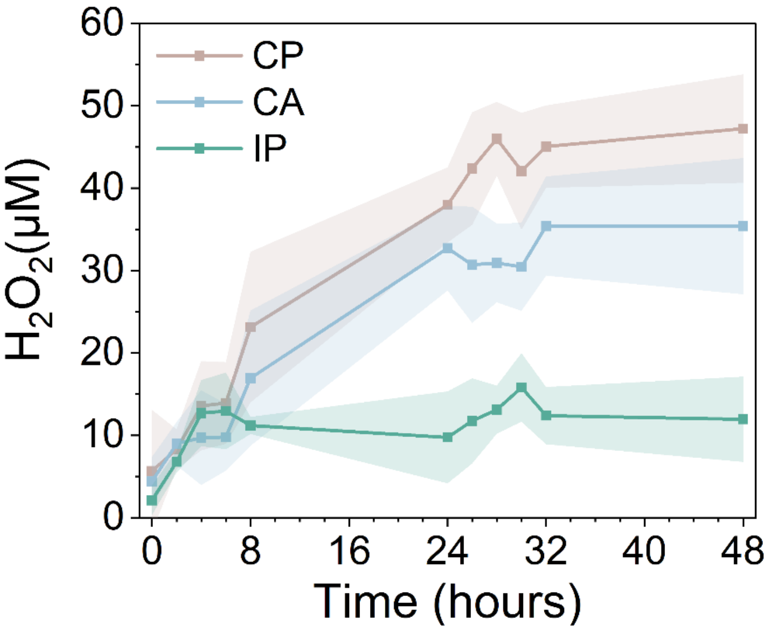

**Figure S20.** The concentration of $H_2O_2$ under different aeration conditions: constant aeration (CA), continuous perturbation (CP), and intermittent perturbation (IP). The error envelope represents standard deviations (technical duplication results for biological triplicates; n=6).

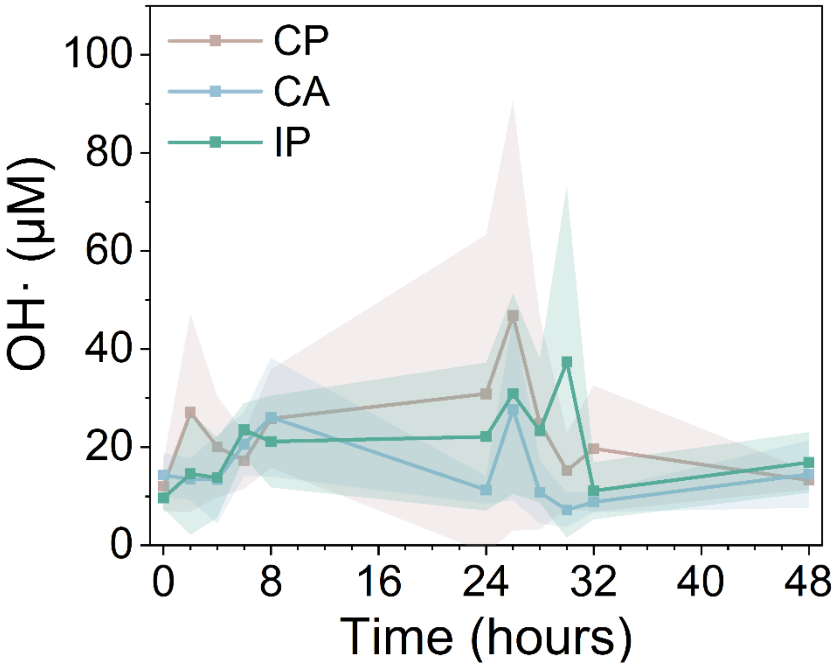

**Figure S21.** The concentration of OH· under different aeration conditions: constant aeration (CA), continuous perturbation (CP), and intermittent perturbation (IP). The error envelope represents standard deviations (technical duplication results for biological triplicates; n=6).

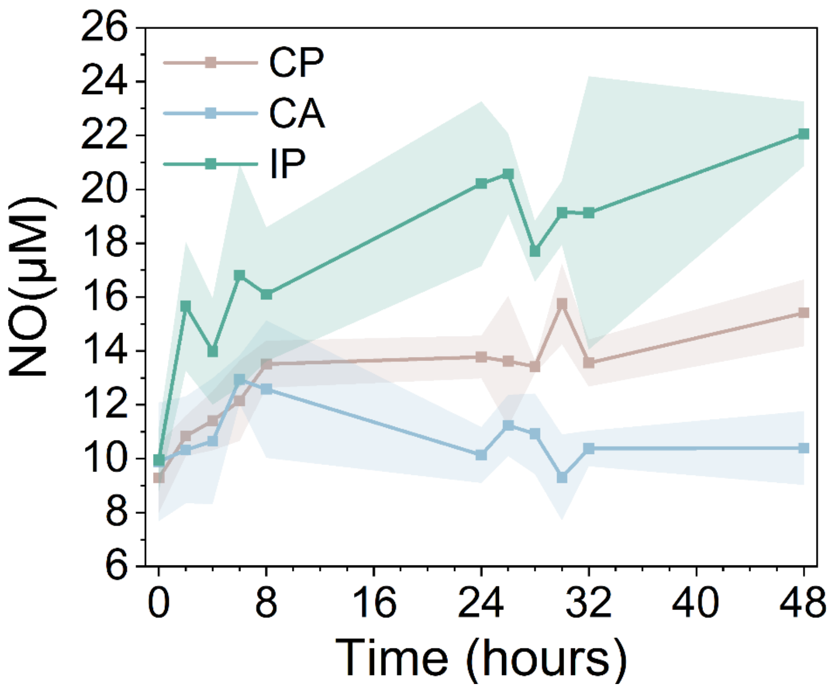

**Figure S22.** The concentration of NO under different aeration conditions: constant aeration (CA), continuous perturbation (CP), and intermittent perturbation (IP). The error envelope represents standard deviations (technical duplication results for biological triplicates; n=6).

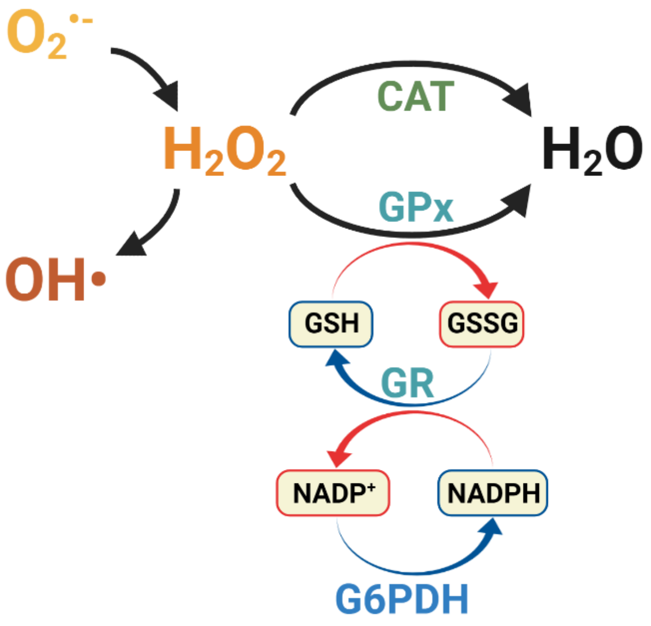

**Figure S23.** Catalase (CAT) and glutathione peroxidase (GPx) system, including the key activities related with $H_2O_2$ scavenging and recycling of the reduced (GSH) and oxidized (GSSG) forms of glutathione.

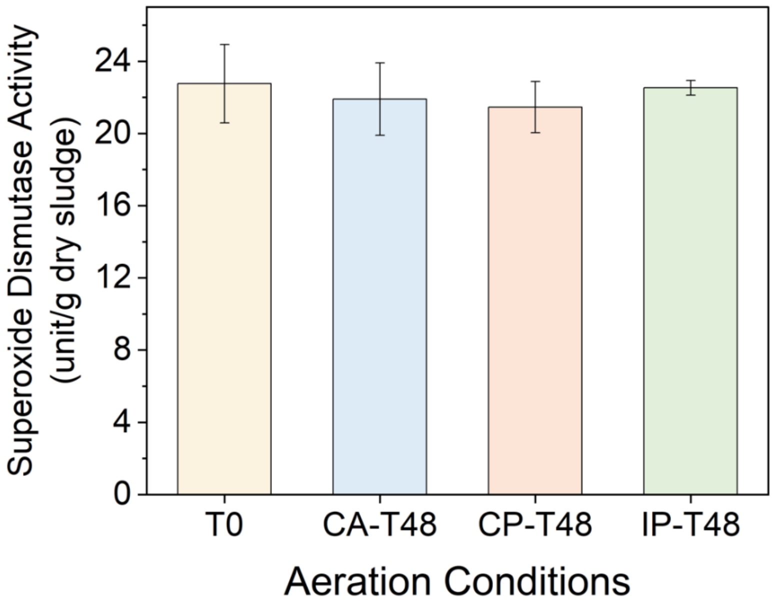

**Figure S24.** The activity of superoxide dismutase following incubation for 48 hours under different aeration conditions: constant aeration (CA), continuous perturbation (CP), and intermittent perturbation (IP). The error bars represent standard deviations (technical duplication results for biological triplicates; n=6).

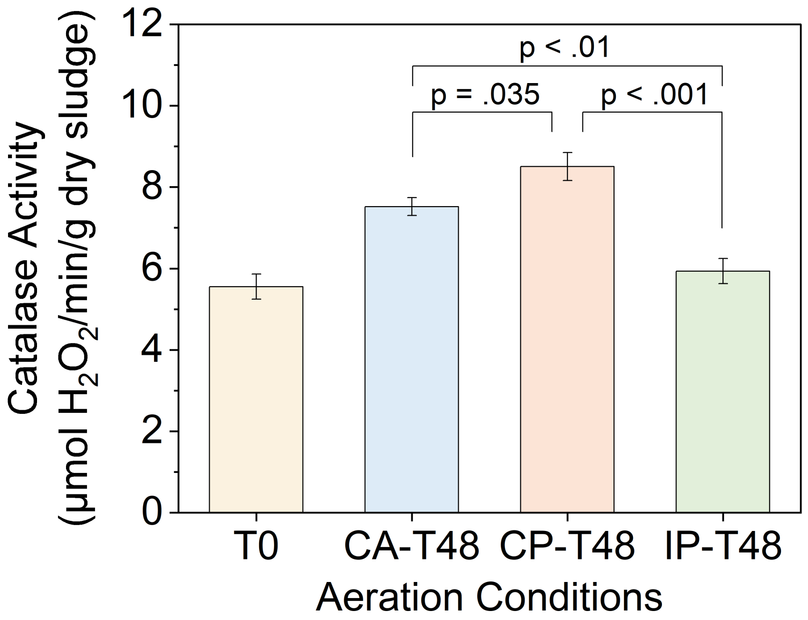

**Figure S25.** The activity of catalase under different aeration conditions: constant aeration (CA), continuous perturbation (CP), and intermittent perturbation (IP). The error bars represent standard deviations (technical duplication results for biological triplicates; n=6).

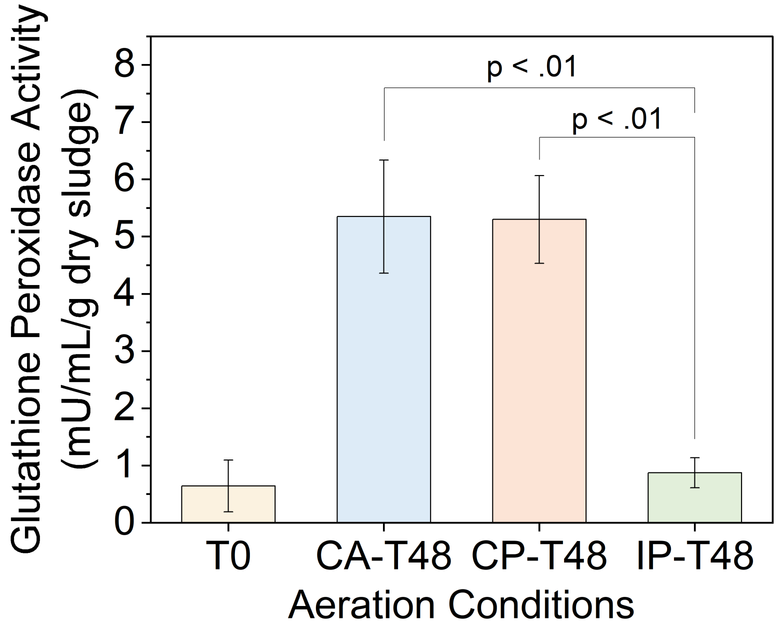

**Figure S26.** The activity of glutathione peroxidase under different aeration conditions: constant aeration (CA), continuous perturbation (CP), and intermittent perturbation (IP). The error bars represent standard deviations (technical duplication results for biological triplicates; n=6).

#### S2.6. Activation of regulons by oxygen variation and reactive oxygen and nitrogen species under oxygen perturbations

**Table S7.** Details of enzymes with significant changes in abundance after 48 hours of treatment under constant aeration (CA). T0 condition represents samples taken before aeration treatment. The numbers following the aeration conditions in the table represent biological replicates.

| Regulon group | Enzyme | T0_1 | T0_2 | T0_3 | CA_1 | CA_2 | CA_3 | P value raw | t ratio | P value adjusted | log2(FC) | Gene name |
|---|---|---|---|---|---|---|---|---|---|---|---|---|
| FNR | glutamate synthase (NADPH) large chain [EC:1.4.1.13] | 0 | 0 | 0 | 13343.6 | 11738.9 | 12230.9 | 0.000013 | 26.2 | 0.019437 | 10.249 | gltB |
| FNR | glucose-6-phosphate 1-dehydrogenase [EC:1.1.1.49 1.1.1.363] | 51293.8 | 46928.1 | 54770.1 | 108453 | 110479 | 113371 | 0.000024 | 22.3 | 0.020135 | 1.119 | G6PD, zwf |
| FNR | nitrate reductase / nitrite oxidoreductase, beta subunit [EC:1.7.5.1 1.7.99.-] | 63565 | 64206.9 | 58338.5 | 34531.3 | 35121.3 | 36032.8 | 0.000149 | 14.04 | 0.040185 | -0.816 | narH, narY, nxrB |
| FNR | large subunit ribosomal protein L23 | 11602.9 | 13112.9 | 12481.7 | 19679.5 | 21064.3 | 20886 | 0.000191 | 13.2 | 0.040185 | 0.728 | RP-L23, MRPL23, rplW |
| FNR | 6-phosphogluconate dehydrogenase [EC:1.1.1.44 1.1.1.343] | 36458.2 | 41905 | 38005.8 | 74904.7 | 84401.2 | 78790.9 | 0.000222 | 12.69 | 0.042723 | 1.033 | PGD, gnd, gntZ |
| FNR | large subunit ribosomal protein L19 | 41388 | 39987.3 | 40078 | 22751.5 | 22513.9 | 27137 | 0.00048 | 10.42 | 0.045874 | -0.746 | RP-L19, MRPL19, rplS |
| FNR | succinate dehydrogenase / fumarate reductase, flavoprotein subunit [EC:1.3.5.1 1.3.5.4] | 32505.6 | 33568.3 | 32822.4 | 18846.7 | 13600.1 | 14757.7 | 0.000445 | 10.62 | 0.045874 | -1.067 | sdhA, frdA |
| FNR | succinate dehydrogenase / fumarate reductase, flavoprotein subunit [EC:1.3.5.1 1.3.5.4] | 66909.2 | 74270.8 | 63588.3 | 37495.1 | 35932.6 | 35068.9 | 0.00058 | 9.919 | 0.04773 | -0.916 | sdhA, frdA |
| FNR | pyruvate dehydrogenase E2 component (dihydrolipoamide acetyltransferase) [EC:2.3.1.12] | 30883.3 | 28182.4 | 25825.2 | 73875.8 | 62028.7 | 74602.1 | 0.00064 | 9.671 | 0.04773 | 1.310 | DLAT, aceF, pdhC |
| ArcA | single-strand DNA-binding protein | 3854.67 | 2985.98 | 2942.09 | 17833.2 | 17392.7 | 16579.2 | 0.000008 | 29.65 | 0.017828 | 2.405 | ssb |

| | | | | | | | | | | | | |
|---|---|---|---|---|---|---|---|---|---|---|---|---|
| ArcA | large subunit ribosomal protein L23 | 11602.9 | 13112.9 | 12481.7 | 19679.5 | 21064.3 | 20886 | 0.000191 | 13.2 | 0.040185 | 0.728 | RP-L23, MRPL23, rplW |
| ArcA | isocitrate lyase [EC:4.1.3.1] | 36238.2 | 35494.3 | 37243.8 | 12996 | 15852.3 | 10178.8 | 0.000169 | 13.6 | 0.040185 | -1.481 | aceA |
| ArcA | oligopeptide transport system substrate-binding protein | 0 | 0 | 0 | 879.704 | 865.326 | 1146.78 | 0.00046 | 10.53 | 0.045874 | 6.559 | oppA, mppA |
| ArcA | isocitrate dehydrogenase [EC:1.1.1.42] | 233333 | 253577 | 229432 | 153664 | 138960 | 149865 | 0.000463 | 10.51 | 0.045874 | -0.695 | IDH1, IDH2, icd |
| ArcA | succinate dehydrogenase / fumarate reductase, flavoprotein subunit [EC:1.3.5.1 1.3.5.4] | 32505.6 | 33568.3 | 32822.4 | 18846.7 | 13600.1 | 14757.7 | 0.000445 | 10.62 | 0.045874 | -1.067 | sdhA, frdA |
| ArcA | succinate dehydrogenase / fumarate reductase, flavoprotein subunit [EC:1.3.5.1 1.3.5.4] | 66909.2 | 74270.8 | 63588.3 | 37495.1 | 35932.6 | 35068.9 | 0.00058 | 9.919 | 0.04773 | -0.916 | sdhA, frdA |
| ArcA | pyruvate dehydrogenase E2 component (dihydrolipoamide acetyltransferase) [EC:2.3.1.12] | 30883.3 | 28182.4 | 25825.2 | 73875.8 | 62028.7 | 74602.1 | 0.00064 | 9.671 | 0.04773 | 1.310 | DLAT, aceF, pdhC |
| ArcA | malate dehydrogenase [EC:1.1.1.37] | 54107 | 61230.5 | 58587.2 | 37943.8 | 36981.2 | 36199.4 | 0.000611 | 9.785 | 0.04773 | -0.646 | mdh |
| SoxRS | glucose-6-phosphate 1-dehydrogenase [EC:1.1.1.49 1.1.1.363] | 51293.8 | 46928.1 | 54770.1 | 108453 | 110479 | 113371 | 0.000024 | 22.3 | 0.020135 | 1.119 | G6PD, zwf |
| SoxRS | Fur family transcriptional regulator, ferric uptake regulator | 6946.27 | 6752.01 | 2797.04 | 21433.1 | 21286.4 | 20790.2 | 0.000329 | 11.47 | 0.044546 | 1.945 | fur, zur, furB |
| OxyR | Fur family transcriptional regulator, ferric uptake regulator | 6946.27 | 6752.01 | 2797.04 | 21433.1 | 21286.4 | 20790.2 | 0.000329 | 11.47 | 0.044546 | 1.945 | fur, zur, furB |
| OxyR | Fe-S cluster assembly protein SufD | 0 | 0 | 0 | 17955.3 | 16294.1 | 13169.6 | 0.000354 | 11.27 | 0.044546 | 10.595 | sufD |
| NsrR | Fe-S cluster assembly protein SufD | 0 | 0 | 0 | 17955.3 | 16294.1 | 13169.6 | 0.000354 | 11.27 | 0.044546 | 10.595 | sufD |
| NsrR | pyruvate dehydrogenase E2 component (dihydrolipoamide acetyltransferase) [EC:2.3.1.12] | 30883.3 | 28182.4 | 25825.2 | 73875.8 | 62028.7 | 74602.1 | 0.00064 | 9.671 | 0.04773 | 1.310 | DLAT, aceF, pdhC |

**Table S8.** Details of enzymes with significant changes in abundance after 48 hours of treatment under continuous perturbation (CP). T0 condition represents samples taken before aeration treatment. The numbers following the aeration conditions in the table represent biological replicates.

| Regulon _group | Enzyme | T0_1 | T0_2 | T0_3 | CP_1 | CP_2 | CP_3 | P value raw | t ratio | P value adjusted | log2(FC) | Gene name |
|---|---|---|---|---|---|---|---|---|---|---|---|---|
| FNR | nitrate reductase / nitrite oxidoreductase, alpha subunit [EC:1.7.5.1 1.7.99.-] | 128817 | 131541 | 129622 | 74390.5 | 73952.5 | 78813.8 | 0.000006 | 31.01 | 0.0048 | -0.780 | narG, narZ, nxrA |
| FNR | glucose-6-phosphate 1-dehydrogenase [EC:1.1.1.49 1.1.1.363] | 51293.8 | 46928.1 | 53941.6 | 139356 | 131864 | 136396 | 0.000009 | 28.5 | 0.004819 | 1.422 | G6PD, zwf |
| FNR | phosphate transport system substrate-binding protein | 1503.36 | 1280.27 | 1303.24 | 0 | 0 | 0 | 0.000043 | 19.23 | 0.008242 | -7.058 | pstS |
| FNR | pyruvate dehydrogenase E2 component (dihydrolipoamide acetyltransferase) [EC:2.3.1.12] | 0 | 0 | 0 | 10091 | 8968.9 | 8671.73 | 0.000028 | 21.39 | 0.008242 | 9.821 | DLAT, aceF, pdhC |
| FNR | large subunit ribosomal protein L19 | 27571.6 | 29083 | 31739.7 | 50633.2 | 49225.6 | 50106 | 0.00009 | 15.96 | 0.011457 | 0.763 | RP-L19, MRPL19, rplS |
| FNR | glutamate dehydrogenase (NADP+) [EC:1.4.1.4] | 45069.3 | 47105.7 | 53463.3 | 136978 | 150359 | 157758 | 0.00011 | 15.16 | 0.011787 | 1.612 | gdhA |
| FNR | nitric oxide dioxygenase [EC:1.14.12.17] | 3917.69 | 18050.2 | 8353.47 | 77825.1 | 70714.9 | 74696.9 | 0.000159 | 13.82 | 0.014215 | 2.880 | hmp, YHB1 |
| FNR | succinyl-CoA synthetase beta subunit [EC:6.2.1.5] | 44089.3 | 49204.2 | 43963.1 | 18868.4 | 18884.2 | 21857.8 | 0.000202 | 12.99 | 0.01614 | -1.203 | sucC |
| FNR | large subunit ribosomal protein L2 | 189864 | 192238 | 197219 | 135393 | 147097 | 141945 | 0.000212 | 12.84 | 0.01614 | -0.449 | RP-L2, MRPL2, rplB |
| FNR | glutamate synthase (NADPH) large chain [EC:1.4.1.13] | 0 | 0 | 0 | 10228.8 | 12804 | 13689.1 | 0.000296 | 11.79 | 0.017929 | 10.226 | gltB |
| FNR | pyruvate dehydrogenase E1 component [EC:1.2.4.1] | 0 | 0 | 0 | 6175.5 | 6544.59 | 8334.28 | 0.00046 | 10.53 | 0.022242 | 9.423 | aceE |
| FNR | formate dehydrogenase major subunit [EC:1.17.1.9] | 61894.9 | 54864.4 | 54193.8 | 89163.7 | 99578.8 | 98065.3 | 0.000693 | 9.472 | 0.02695 | 0.746 | fdoG, fdhF, fdwA |

| | | | | | | | | | | | | |
|---|---|---|---|---|---|---|---|---|---|---|---|---|
| FNR | small subunit ribosomal protein S16 | 25051.6 | 24477.3 | 24013.6 | 49227.8 | 54534.8 | 45123.6 | 0.000787 | 9.164 | 0.028097 | 1.018 | RP-S16, MRPS16, rpsP |
| FNR | nitric oxide dioxygenase [EC:1.14.12.17] | 10041.9 | 7165.11 | 3296.56 | 79067.4 | 57595.1 | 65261 | 0.000778 | 9.191 | 0.028097 | 3.300 | hmp, YHB1 |
| FNR | pyruvate dehydrogenase E1 component [EC:1.2.4.1] | 28571.7 | 39091.4 | 34422.1 | 80135.7 | 88689.2 | 97830.6 | 0.000767 | 9.225 | 0.028097 | 1.385 | aceE |
| FNR | 6-phosphogluconate dehydrogenase [EC:1.1.1.44 1.1.1.343] | 36458.2 | 41905 | 35672 | 78353 | 87525.1 | 97172.9 | 0.001005 | 8.6 | 0.030536 | 1.206 | PGD, gnd, gntZ |
| FNR | dihydrolipoamide dehydrogenase [EC:1.8.1.4] | 14343 | 16994.2 | 16887.4 | 2721.05 | 6230.1 | 2529.17 | 0.001171 | 8.262 | 0.033043 | -2.071 | DLD, lpd, pdhD |
| FNR | large subunit ribosomal protein L3 | 16729.2 | 20642 | 20879.9 | 33149.9 | 36510.4 | 32835.8 | 0.001176 | 8.253 | 0.033043 | 0.815 | RP-L3, MRPL3, rplC |
| FNR | succinate dehydrogenase / fumarate reductase, flavoprotein subunit [EC:1.3.5.1 1.3.5.4] | 66909.2 | 74270.8 | 60954.3 | 36721.8 | 36399.8 | 35819.6 | 0.001295 | 8.047 | 0.034948 | -0.892 | sdhA, frdA |
| FNR | large subunit ribosomal protein L20 | 24603 | 27098.6 | 25740.1 | 39099.7 | 39321.5 | 44492.1 | 0.001348 | 7.962 | 0.035755 | 0.666 | RP-L20, MRPL20, rplT |
| FNR | small subunit ribosomal protein S17 | 99334.3 | 92763.6 | 93220.4 | 140910 | 140616 | 159965 | 0.001515 | 7.721 | 0.037091 | 0.630 | RP-S17, MRPS17, rpsQ |
| FNR | nitric oxide dioxygenase [EC:1.14.12.17] | 16368.4 | 11965.7 | 9646.97 | 78631.8 | 76605.2 | 105977 | 0.001537 | 7.692 | 0.037091 | 2.782 | hmp, YHB1 |
| FNR | small subunit ribosomal protein S3 | 91516.3 | 124789 | 105631 | 186595 | 179738 | 191663 | 0.001544 | 7.683 | 0.037091 | 0.793 | RP-S3, rpsC |
| FNR | NADH-quinone oxidoreductase subunit G [EC:7.1.1.2] | 14122.3 | 16517 | 16044.1 | 8916.84 | 6941.83 | 8806.04 | 0.001654 | 7.544 | 0.038536 | -0.920 | nuoG |
| FNR | type VI secretion system secreted protein Hcp | 9944.86 | 12624.4 | 10023.5 | 3457.18 | 4544.77 | 3427.6 | 0.001784 | 7.394 | 0.039355 | -1.512 | hcp |
| FNR | nitrate reductase / nitrite oxidoreductase, alpha subunit [EC:1.7.5.1 1.7.99.-] | 90178.3 | 101139 | 80103.4 | 40979.5 | 39579.1 | 47873.3 | 0.001942 | 7.229 | 0.040665 | -1.080 | narG, narZ, nxrA |
| FNR | electron transfer flavoprotein beta subunit | 0 | 0 | 0 | 1771.83 | 2707.44 | 2842.61 | 0.001922 | 7.249 | 0.040665 | 7.899 | fixA, etfB |

| | | | | | | | | | | | |
|---|---|---|---|---|---|---|---|---|---|---|---|
| FNR | succinate dehydrogenase / fumarate reductase, flavoprotein subunit [EC:1.3.5.1 1.3.5.4] | 32505.6 | 33568.3 | 36515.2 | 18669.5 | 23060 | 18057.4 | 0.001966 | 7.206 | 0.040702 | -0.779 | sdhA, frdA |
| FNR | phosphate transport system protein | 4789.28 | 4751.33 | 8029.23 | 18566.1 | 15066.2 | 16012.1 | 0.002088 | 7.092 | 0.041483 | 1.499 | phoU |
| FNR | nitrate reductase / nitrite oxidoreductase, beta subunit [EC:1.7.5.1 1.7.99.-] | 70958.4 | 62046.7 | 59955.5 | 39697.4 | 40792.1 | 35574.4 | 0.002346 | 6.874 | 0.044108 | -0.733 | narH, narY, nxrB |
| FNR | nitric oxide dioxygenase [EC:1.14.12.17] | 23041.4 | 17806.5 | 14701.3 | 70317.7 | 88966.8 | 104250 | 0.002367 | 6.858 | 0.044108 | 2.246 | hmp, YHB1 |
| FNR | aerotaxis receptor | 6984.59 | 13861.3 | 17682.4 | 37176.6 | 33922.6 | 34443.5 | 0.002453 | 6.793 | 0.04467 | 1.454 | aer |
| FNR | 2-oxoglutarate dehydrogenase E2 component (dihydrolipoamide succinyltransferase) [EC:2.3.1.61] | 5727.35 | 5971.1 | 4770.45 | 1697.66 | 2820.61 | 1691.7 | 0.002858 | 6.52 | 0.048167 | -1.407 | DLST, sucB |
| FNR | 6-phosphogluconate dehydrogenase [EC:1.1.1.44 1.1.1.343] | 49328.2 | 61628.1 | 67565.8 | 121712 | 102450 | 124953 | 0.003006 | 6.432 | 0.049061 | 0.968 | PGD, gnd, gntZ |
| FNR | nitrogen regulatory protein P-II 2 | 434.683 | 480.853 | 275.186 | 0 | 0 | 0 | 0.003114 | 6.371 | 0.049269 | -5.279 | glnK |
| FNR | cytochrome c biogenesis protein CcmG, thiol:disulfide interchange protein DsbE | 1343.28 | 1078.35 | 1844.81 | 0 | 0 | 0 | 0.003192 | 6.328 | 0.04951 | -7.120 | ccmG, dsbE |
| ArcA | pyruvate dehydrogenase E2 component (dihydrolipoamide acetyltransferase) [EC:2.3.1.12] | 0 | 0 | 0 | 10091 | 8968.9 | 8671.73 | 0.000028 | 21.39 | 0.008242 | 9.821 | DLAT, aceF, pdhC |
| ArcA | succinyl-CoA synthetase beta subunit [EC:6.2.1.5] | 44089.3 | 49204.2 | 43963.1 | 18868.4 | 18884.2 | 21857.8 | 0.000202 | 12.99 | 0.01614 | -1.203 | sucC |
| ArcA | large subunit ribosomal protein L2 | 189864 | 192238 | 197219 | 135393 | 147097 | 141945 | 0.000212 | 12.84 | 0.01614 | -0.449 | RP-L2, MRPL2, rplB |
| ArcA | HTH-type transcriptional regulator, repressor for puuD | 0 | 0 | 0 | 3499.84 | 3960.8 | 2972.92 | 0.00026 | 12.19 | 0.016809 | 8.410 | puuR |
| ArcA | pyruvate dehydrogenase E1 component [EC:1.2.4.1] | 0 | 0 | 0 | 6175.5 | 6544.59 | 8334.28 | 0.00046 | 10.53 | 0.022242 | 9.423 | aceE |
| ArcA | pyruvate dehydrogenase E1 component [EC:1.2.4.1] | 28571.7 | 39091.4 | 34422.1 | 80135.7 | 88689.2 | 97830.6 | 0.000767 | 9.225 | 0.028097 | 1.385 | aceE |
| ArcA | dihydrolipoamide dehydrogenase [EC:1.8.1.4] | 14343 | 16994.2 | 16887.4 | 2721.05 | 6230.1 | 2529.17 | 0.001171 | 8.262 | 0.033043 | -2.071 | DLD, lpd, pdhD |

| | | | | | | | | | | | | |
|---|---|---|---|---|---|---|---|---|---|---|---|---|
| ArcA | large subunit ribosomal protein L3 | 16729.2 | 20642 | 20879.9 | 33149.9 | 36510.4 | 32835.8 | 0.001176 | 8.253 | 0.033043 | 0.815 | RP-L3, MRPL3, rplC |
| ArcA | succinate dehydrogenase / fumarate reductase, flavoprotein subunit [EC:1.3.5.1 1.3.5.4] | 66909.2 | 74270.8 | 60954.3 | 36721.8 | 36399.8 | 35819.6 | 0.001295 | 8.047 | 0.034948 | -0.892 | sdhA, frdA |
| ArcA | small subunit ribosomal protein S17 | 99334.3 | 92763.6 | 93220.4 | 140910 | 140616 | 159965 | 0.001515 | 7.721 | 0.037091 | 0.630 | RP-S17, MRPS17 , rpsQ |
| ArcA | small subunit ribosomal protein S3 | 91516.3 | 124789 | 105631 | 186595 | 179738 | 191663 | 0.001544 | 7.683 | 0.037091 | 0.793 | RP-S3, rpsC |
| ArcA | NADH-quinone oxidoreductase subunit G [EC:7.1.1.2] | 14122.3 | 16517 | 16044.1 | 8916.84 | 6941.83 | 8806.04 | 0.001654 | 7.544 | 0.038536 | -0.920 | nuoG |
| ArcA | succinate dehydrogenase / fumarate reductase, flavoprotein subunit [EC:1.3.5.1 1.3.5.4] | 32505.6 | 33568.3 | 36515.2 | 18669.5 | 23060 | 18057.4 | 0.001966 | 7.206 | 0.040702 | -0.779 | sdhA, frdA |
| ArcA | long-chain fatty acid transport protein | 15791.9 | 17034.8 | 18369.6 | 6018.41 | 0 | 0 | 0.002148 | 7.038 | 0.041483 | -3.089 | fadL |
| ArcA | 2-oxoglutarate dehydrogenase E2 component (dihydrolipoamide succinyltransferase) [EC:2.3.1.61] | 5727.35 | 5971.1 | 4770.45 | 1697.66 | 2820.61 | 1691.7 | 0.002858 | 6.52 | 0.048167 | -1.407 | DLST, sucB |
| ArcA | aconitate hydratase 2 / 2-methylisocitrate dehydratase [EC:4.2.1.3 4.2.1.99] | 39372.6 | 47809.9 | 41372.4 | 24136.3 | 27041.2 | 22902.9 | 0.003015 | 6.426 | 0.049061 | -0.795 | acnB |
| ArcA | citrate synthase [EC:2.3.3.1] | 30776.5 | 30300.9 | 28436 | 16184 | 21491.1 | 19092.4 | 0.002975 | 6.449 | 0.049061 | -0.657 | CS, gltA |
| ArcA | malate synthase [EC:2.3.3.9] | 4689.24 | 8134.97 | 7493.19 | 0 | 0 | 0 | 0.003058 | 6.401 | 0.049183 | -9.372 | aceB, glcB |
| SoxRS | glucose-6-phosphate 1-dehydrogenase [EC:1.1.1.49 1.1.1.363] | 51293.8 | 46928.1 | 53941.6 | 139356 | 131864 | 136396 | 0.000009 | 28.5 | 0.004819 | 1.422 | G6PD, zwf |
| SoxRS | superoxide dismutase, Fe-Mn family [EC:1.15.1.1] | 0 | 0 | 0 | 12008.1 | 10964.6 | 12161.4 | 0.000006 | 31.15 | 0.0048 | 10.162 | SOD2 |
| SoxRS | superoxide dismutase, Fe-Mn family [EC:1.15.1.1] | 4038.84 | 4909.95 | 5352.44 | 0 | 0 | 0 | 0.000247 | 12.35 | 0.016649 | -8.865 | SOD2 |
| SoxRS | superoxide dismutase, Fe-Mn family [EC:1.15.1.1] | 15653.4 | 19203.1 | 18300.7 | 62974.8 | 64592.4 | 75652.5 | 0.000265 | 12.13 | 0.016809 | 1.935 | SOD2 |
| SoxRS | superoxide dismutase, Fe-Mn family [EC:1.15.1.1] | 9493.36 | 6448.72 | 8601.4 | 0 | 0 | 0 | 0.000825 | 9.053 | 0.028097 | -8.865 | SOD2 |

| | | | | | | | | | | | | |
|---|---|---|---|---|---|---|---|---|---|---|---|---|
| OxyR | starvation-inducible DNA-binding protein | 0 | 0 | 0 | 3997.29 | 3338.11 | 3162.57 | 0.000161 | 13.77 | 0.014215 | 8.419 | dps |
| OxyR | peroxiredoxin (alkyl hydroperoxide reductase subunit C) [EC:1.11.1.15] | 13807.1 | 17070.5 | 17383.3 | 53242.7 | 61643.3 | 66440.2 | 0.000385 | 11.03 | 0.019612 | 1.910 | PRDX2_4, ahpC |
| OxyR | type VI secretion system secreted protein Hcp | 9944.86 | 12624.4 | 10023.5 | 3457.18 | 4544.77 | 3427.6 | 0.001784 | 7.394 | 0.039355 | -1.512 | hcp |
| OxyR | peroxiredoxin (alkyl hydroperoxide reductase subunit C) [EC:1.11.1.15] | 93723.5 | 112062 | 102932 | 67876.4 | 68866.3 | 69810.5 | 0.003066 | 6.397 | 0.049183 | -0.580 | PRDX2_4, ahpC |
| NorR | nitric oxide dioxygenase [EC:1.14.12.17] | 3917.69 | 18050.2 | 8353.47 | 77825.1 | 70714.9 | 74696.9 | 0.000159 | 13.82 | 0.014215 | 2.880 | hmp, YHB1 |
| NorR | nitric oxide reductase subunit B [EC:1.7.2.5] | 7643.01 | 5734.95 | 8830.95 | 36022 | 44572.7 | 40398.9 | 0.000233 | 12.53 | 0.016478 | 2.446 | norB |
| NorR | nitric oxide dioxygenase [EC:1.14.12.17] | 10041.9 | 7165.11 | 3296.56 | 79067.4 | 57595.1 | 65261 | 0.000778 | 9.191 | 0.028097 | 3.300 | hmp, YHB1 |
| NorR | nitric oxide dioxygenase [EC:1.14.12.17] | 16368.4 | 11965.7 | 9646.97 | 78631.8 | 76605.2 | 105977 | 0.001537 | 7.692 | 0.037091 | 2.782 | hmp, YHB1 |
| NorR | nitric oxide dioxygenase [EC:1.14.12.17] | 23041.4 | 17806.5 | 14701.3 | 70317.7 | 88966.8 | 104250 | 0.002367 | 6.858 | 0.044108 | 2.246 | hmp, YHB1 |
| NsrR | pyruvate dehydrogenase E2 component (dihydrolipoamide acetyltransferase) [EC:2.3.1.12] | 0 | 0 | 0 | 10091 | 8968.9 | 8671.73 | 0.000028 | 21.39 | 0.008242 | 9.821 | DLAT, aceF, pdhC |
| NsrR | nitric oxide dioxygenase [EC:1.14.12.17] | 3917.69 | 18050.2 | 8353.47 | 77825.1 | 70714.9 | 74696.9 | 0.000159 | 13.82 | 0.014215 | 2.880 | hmp, YHB1 |
| NsrR | pyruvate dehydrogenase E1 component [EC:1.2.4.1] | 0 | 0 | 0 | 6175.5 | 6544.59 | 8334.28 | 0.00046 | 10.53 | 0.022242 | 9.423 | aceE |
| NsrR | nitric oxide dioxygenase [EC:1.14.12.17] | 10041.9 | 7165.11 | 3296.56 | 79067.4 | 57595.1 | 65261 | 0.000778 | 9.191 | 0.028097 | 3.300 | hmp, YHB1 |
| NsrR | pyruvate dehydrogenase E1 component [EC:1.2.4.1] | 28571.7 | 39091.4 | 34422.1 | 80135.7 | 88689.2 | 97830.6 | 0.000767 | 9.225 | 0.028097 | 1.385 | aceE |
| NsrR | large subunit ribosomal protein L20 | 24603 | 27098.6 | 25740.1 | 39099.7 | 39321.5 | 44492.1 | 0.001348 | 7.962 | 0.035755 | 0.666 | RP-L20, MRPL20, rplT |
| NsrR | nitric oxide dioxygenase [EC:1.14.12.17] | 16368.4 | 11965.7 | 9646.97 | 78631.8 | 76605.2 | 105977 | 0.001537 | 7.692 | 0.037091 | 2.782 | hmp, YHB1 |
| NsrR | type VI secretion system secreted protein Hcp | 9944.86 | 12624.4 | 10023.5 | 3457.18 | 4544.77 | 3427.6 | 0.001784 | 7.394 | 0.039355 | -1.512 | hcp |
| NsrR | nitric oxide dioxygenase [EC:1.14.12.17] | 23041.4 | 17806.5 | 14701.3 | 70317.7 | 88966.8 | 104250 | 0.002367 | 6.858 | 0.044108 | 2.246 | hmp, YHB1 |

**Table S9.** Details of enzymes with significant changes in abundance after 48 hours of treatment under intermittent perturbation (IP). T0 condition represents samples taken before aeration treatment. The numbers following the aeration conditions in the table represent biological replicates.

| Regulon _group | Enzyme | T0_1 | T0_2 | T0_3 | IP_1 | IP_2 | IP_3 | P value raw | t ratio | P value adjusted | log2(FC) | Gene name |
|---|---|---|---|---|---|---|---|---|---|---|---|---|
| FNR | MFS transporter, NNP family, nitrate/nitrite transporter | 0 | 0 | 0 | 1611.06 | 1612.92 | 1414.33 | 0.00002 | 23.47 | 0.004804 | 7.241 | NRT, narK, nrtP, nasA |
| FNR | dihydrolipoamide dehydrogenase [EC:1.8.1.4] | 14343 | 16994.2 | 17976.4 | 0 | 0 | 0 | 0.000111 | 15.15 | 0.012389 | -10.651 | DLD, lpd, pdhD |
| FNR | pyruvate dehydrogenase E1 component [EC:1.2.4.1] | 0 | 0 | 0 | 13523.1 | 11544.3 | 14622.2 | 0.000125 | 14.69 | 0.013646 | 10.338 | aceE |
| FNR | succinate dehydrogenase / fumarate reductase, flavoprotein subunit [EC:1.3.5.1 1.3.5.4] | 66909.2 | 74270.8 | 75185.9 | 36015.6 | 34855.8 | 36099.7 | 0.000162 | 13.76 | 0.015671 | -1.016 | sdhA, frdA |
| FNR | cytochrome c biogenesis protein CcmG, thiol:disulfide interchange protein DsbE | 35928.6 | 37188.6 | 28821.5 | 0 | 0 | 0 | 0.000199 | 13.05 | 0.017328 | -11.699 | ccmG, dsbE |
| FNR | large subunit ribosomal protein L4 | 6650.65 | 4212.22 | 7579.41 | 25768.7 | 22398.4 | 24509.6 | 0.000211 | 12.86 | 0.017328 | 1.978 | RP-L4, MRPL4, rplD |
| FNR | nitric oxide dioxygenase [EC:1.14.12.17] | 10041.9 | 7165.11 | 8532.84 | 46180.6 | 36744.4 | 42583.7 | 0.000318 | 11.58 | 0.019519 | 2.286 | hmp, YHB1 |
| FNR | nitric oxide dioxygenase [EC:1.14.12.17] | 23041.4 | 17806.5 | 5607.52 | 123041 | 106145 | 98425.9 | 0.000463 | 10.51 | 0.022614 | 2.818 | hmp, YHB1 |
| FNR | nitric oxide dioxygenase [EC:1.14.12.17] | 16368.4 | 11965.7 | 13527.5 | 103693 | 106942 | 81653.4 | 0.000488 | 10.37 | 0.022638 | 2.804 | hmp, YHB1 |
| FNR | dihydrolipoamide dehydrogenase [EC:1.8.1.4] | 0 | 0 | 0 | 9352.34 | 8991 | 12133.2 | 0.000514 | 10.23 | 0.022655 | 9.957 | DLD, lpd, pdhD |
| FNR | large subunit ribosomal protein L3 | 3354.29 | 7951.83 | 5785.83 | 19601.4 | 18533 | 18037.7 | 0.000754 | 9.266 | 0.027228 | 1.717 | RP-L3, MRPL3, rplC |
| FNR | nitrate reductase gamma subunit [EC:1.7.5.1 1.7.99.-] | 1636.22 | 500.417 | 623.513 | 7783.74 | 7119.62 | 5927.31 | 0.000761 | 9.246 | 0.027228 | 2.916 | narI, narV |

| | | | | | | | | | | | | |
|---|---|---|---|---|---|---|---|---|---|---|---|---|
| FNR | large subunit ribosomal protein L13 | 12905.1 | 10446.9 | 14088.5 | 22676.8 | 24681.3 | 23822 | 0.000768 | 9.222 | 0.027228 | 0.927 | RP-L13, MRPL13, rplM |
| FNR | nitrate reductase / nitrite oxidoreductase, alpha subunit [EC:1.7.5.1 1.7.99.-] | 6949.85 | 10530.8 | 5312.05 | 22066.1 | 24808.3 | 23707.3 | 0.000781 | 9.183 | 0.027353 | 1.631 | narG, narZ, nxrA |
| FNR | glucose-6-phosphate 1-dehydrogenase [EC:1.1.1.49 1.1.1.363] | 51293.8 | 46928.1 | 52416.4 | 78230.3 | 79974.1 | 72145.8 | 0.000792 | 9.15 | 0.027436 | 0.613 | G6PD, zwf |
| FNR | glutamate synthase (NADPH) small chain [EC:1.4.1.13] | 117102 | 115531 | 106139 | 71232.9 | 59144.3 | 70268.6 | 0.000881 | 8.9 | 0.028255 | -0.756 | gltD |
| FNR | aminomethyltransferase [EC:2.1.2.10] | 7206.69 | 6742.15 | 6435.45 | 1049 | 933.808 | 2661.24 | 0.000948 | 8.73 | 0.028439 | -2.134 | gcvT, AMT |
| FNR | outer membrane protein | 19083.4 | 14650.7 | 13670.4 | 36925.3 | 41676.9 | 35414.5 | 0.000911 | 8.823 | 0.028439 | 1.266 | ompW |
| FNR | NADH-quinone oxidoreductase subunit H [EC:7.1.1.2] | 0 | 0 | 0 | 1997.16 | 1346.6 | 1516.3 | 0.001143 | 8.315 | 0.031393 | 7.308 | nuoH |
| FNR | nitric oxide dioxygenase [EC:1.14.12.17] | 3917.69 | 18050.2 | 9216.51 | 111616 | 101066 | 76993.2 | 0.001456 | 7.802 | 0.035162 | 3.216 | hmp, YHB1 |
| FNR | pyruvate dehydrogenase E1 component [EC:1.2.4.1] | 66745.4 | 80522 | 68765.8 | 33193 | 39480.4 | 35733.5 | 0.001541 | 7.687 | 0.036253 | -0.995 | aceE |
| FNR | thiol peroxidase, atypical 2-Cys peroxiredoxin [EC:1.11.1.15] | 0 | 3380 | 1043.91 | 10256.2 | 9522.9 | 9042.83 | 0.00155 | 7.675 | 0.036277 | 2.704 | tpx |
| FNR | glutamate synthase (NADPH) small chain [EC:1.4.1.13] | 0 | 0 | 0 | 3804.87 | 3030.79 | 4785.55 | 0.001585 | 7.63 | 0.036538 | 8.566 | gltD |
| FNR | cytochrome c biogenesis protein CcmG, thiol:disulfide interchange protein DsbE | 0 | 1388.88 | 0 | 4778.5 | 5989.53 | 6629.17 | 0.001707 | 7.481 | 0.037582 | 3.647 | ccmG, dsbE |
| FNR | pyruvate dehydrogenase E1 component [EC:1.2.4.1] | 28571.7 | 39091.4 | 29625.5 | 59263.4 | 64831.8 | 59170.3 | 0.001711 | 7.477 | 0.037582 | 0.914 | aceE |
| FNR | nitrate reductase / nitrite oxidoreductase, alpha subunit [EC:1.7.5.1 1.7.99.-] | 17417.3 | 19535.3 | 17620.3 | 30193.2 | 37416.8 | 35978.4 | 0.002103 | 7.078 | 0.04083 | 0.925 | narG, narZ, nxrA |
| FNR | nitrogen regulatory protein P-II 2 | 434.683 | 480.853 | 290.807 | 0 | 0 | 0 | 0.002161 | 7.027 | 0.041143 | -5.298 | glnK |
| FNR | nitrite reductase (NADH) large subunit [EC:1.7.1.15] | 2410.15 | 3074.87 | 1861.81 | 0 | 0 | 0 | 0.002213 | 6.983 | 0.041831 | -7.904 | nirB |
| FNR | 2-oxoglutarate dehydrogenase E1 component [EC:1.2.4.2] | 8884.89 | 11703.4 | 11125.8 | 4542.22 | 3636.23 | 2267.75 | 0.002829 | 6.538 | 0.047366 | -1.602 | OGDH, sucA |

| | | | | | | | | | | | | |
|---|---|---|---|---|---|---|---|---|---|---|---|---|
| FNR | formate dehydrogenase major subunit [EC:1.17.1.9] | 8232.41 | 0 | 5409.66 | 21490.3 | 22623.2 | 27036.4 | 0.002889 | 6.501 | 0.047605 | 2.383 | fdoG, fdhF, fdwA |
| FNR | glutamate dehydrogenase (NAD(P)+) [EC:1.4.1.3] | 50636.8 | 56534.6 | 63059.5 | 15299.7 | 28127 | 24687.1 | 0.002918 | 6.483 | 0.047605 | -1.321 | GLUD1_2, gdhA |
| FNR | thiol peroxidase, atypical 2-Cys peroxiredoxin [EC:1.11.1.15] | 0 | 4154.32 | 4929.34 | 13820.2 | 18287.3 | 15356.9 | 0.003151 | 6.35 | 0.048771 | 2.386 | tpx |
| FNR | thiol peroxidase, atypical 2-Cys peroxiredoxin [EC:1.11.1.15] | 5992.6 | 10846.7 | 4390.66 | 24673.8 | 32292.8 | 24849.7 | 0.003126 | 6.364 | 0.048771 | 1.946 | tpx |
| FNR | type VI secretion system secreted protein Hcp | 59093.2 | 61728.9 | 52071.8 | 32891.5 | 37559.6 | 28356.1 | 0.003244 | 6.3 | 0.049468 | -0.807 | hcp |
| ArcA | dihydrolipoamide dehydrogenase [EC:1.8.1.4] | 14343 | 16994.2 | 17976.4 | 0 | 0 | 0 | 0.000111 | 15.15 | 0.012389 | -10.651 | DLD, lpd, pdhD |
| ArcA | pyruvate dehydrogenase E1 component [EC:1.2.4.1] | 0 | 0 | 0 | 13523.1 | 11544.3 | 14622.2 | 0.000125 | 14.69 | 0.013646 | 10.338 | aceE |
| ArcA | succinate dehydrogenase / fumarate reductase, flavoprotein subunit [EC:1.3.5.1 1.3.5.4] | 66909.2 | 74270.8 | 75185.9 | 36015.6 | 34855.8 | 36099.7 | 0.000162 | 13.76 | 0.015671 | -1.016 | sdhA, frdA |
| ArcA | large subunit ribosomal protein L4 | 6650.65 | 4212.22 | 7579.41 | 25768.7 | 22398.4 | 24509.6 | 0.000211 | 12.86 | 0.017328 | 1.978 | RP-L4, MRPL4, rplD |
| ArcA | single-strand DNA-binding protein | 3854.67 | 2985.98 | 6618.72 | 24653.3 | 21035.7 | 24865.1 | 0.000328 | 11.49 | 0.019519 | 2.390 | ssb |
| ArcA | dihydrolipoamide dehydrogenase [EC:1.8.1.4] | 0 | 0 | 0 | 9352.34 | 8991 | 12133.2 | 0.000514 | 10.23 | 0.022655 | 9.957 | DLD, lpd, pdhD |
| ArcA | H+-translocating NAD(P) transhydrogenase subunit alpha [EC:1.6.1.2 7.1.1.1] | 12534 | 5369.34 | 23142 | 72331.7 | 64639.9 | 65958.9 | 0.000686 | 9.496 | 0.027146 | 2.306 | pntA |
| ArcA | large subunit ribosomal protein L3 | 3354.29 | 7951.83 | 5785.83 | 19601.4 | 18533 | 18037.7 | 0.000754 | 9.266 | 0.027228 | 1.717 | RP-L3, MRPL3, rplC |
| ArcA | HTH-type transcriptional regulator, repressor for puuD | 0 | 0 | 0 | 7062.66 | 8304.42 | 10174.7 | 0.00071 | 9.413 | 0.027228 | 9.702 | puuR |
| ArcA | long-chain acyl-CoA synthetase [EC:6.2.1.3] | 19342.3 | 21247.3 | 21940 | 9494.42 | 9884.29 | 6414.72 | 0.000806 | 9.109 | 0.027588 | -1.278 | ACSL, fadD |
| ArcA | acetyl-CoA acyltransferase [EC:2.3.1.16] | 0 | 0 | 0 | 7382.85 | 5606.82 | 5200.9 | 0.000826 | 9.05 | 0.027865 | 9.212 | fadA, fadI |

| | | | | | | | | | | | | |
|---|---|---|---|---|---|---|---|---|---|---|---|---|
| ArcA | outer membrane protein | 19083.4 | 14650.7 | 13670.4 | 36925.3 | 41676.9 | 35414.5 | 0.000911 | 8.823 | 0.028439 | 1.266 | ompW |
| ArcA | NADH-quinone oxidoreductase subunit H [EC:7.1.1.2] | 0 | 0 | 0 | 1997.16 | 1346.6 | 1516.3 | 0.001143 | 8.315 | 0.031393 | 7.308 | nuoH |
| ArcA | isocitrate dehydrogenase [EC:1.1.1.42] | 62106.5 | 49226.1 | 58961.8 | 25033.8 | 26399.4 | 21365.1 | 0.001447 | 7.815 | 0.035119 | -1.226 | IDH1, IDH2, icd |
| ArcA | pyruvate dehydrogenase E1 component [EC:1.2.4.1] | 66745.4 | 80522 | 68765.8 | 33193 | 39480.4 | 35733.5 | 0.001541 | 7.687 | 0.036253 | -0.995 | aceE |
| ArcA | thiol peroxidase, atypical 2-Cys peroxiredoxin [EC:1.11.1.15] | 0 | 3380 | 1043.91 | 10256.2 | 9522.9 | 9042.83 | 0.00155 | 7.675 | 0.036277 | 2.704 | tpx |
| ArcA | pyruvate dehydrogenase E1 component [EC:1.2.4.1] | 28571.7 | 39091.4 | 29625.5 | 59263.4 | 64831.8 | 59170.3 | 0.001711 | 7.477 | 0.037582 | 0.914 | aceE |
| ArcA | putrescine transport system substrate-binding protein | 24391.7 | 19413.9 | 32589.2 | 54813.8 | 56691.2 | 52868 | 0.001836 | 7.338 | 0.039182 | 1.105 | potF |
| ArcA | long-chain fatty acid transport protein | 15791.9 | 17034.8 | 16034.1 | 0 | 5031.57 | 5479.27 | 0.002065 | 7.113 | 0.040827 | -2.217 | fadL |
| ArcA | 2-oxoglutarate dehydrogenase E1 component [EC:1.2.4.2] | 8884.89 | 11703.4 | 11125.8 | 4542.22 | 3636.23 | 2267.75 | 0.002829 | 6.538 | 0.047366 | -1.602 | OGDH, sucA |
| ArcA | aconitate hydratase 2 / 2-methylisocitrate dehydratase [EC:4.2.1.3 4.2.1.99] | 12904.2 | 12393.5 | 14466.6 | 8037.99 | 6745.8 | 5288.27 | 0.002889 | 6.5 | 0.047605 | -0.986 | acnB |
| ArcA | malate dehydrogenase [EC:1.1.1.37] | 33942.1 | 37358.1 | 41838.1 | 15988.8 | 22419 | 17357.6 | 0.003139 | 6.357 | 0.048771 | -1.021 | mdh |
| ArcA | thiol peroxidase, atypical 2-Cys peroxiredoxin [EC:1.11.1.15] | 0 | 4154.32 | 4929.34 | 13820.2 | 18287.3 | 15356.9 | 0.003151 | 6.35 | 0.048771 | 2.386 | tpx |
| ArcA | thiol peroxidase, atypical 2-Cys peroxiredoxin [EC:1.11.1.15] | 5992.6 | 10846.7 | 4390.66 | 24673.8 | 32292.8 | 24849.7 | 0.003126 | 6.364 | 0.048771 | 1.946 | tpx |
| SoxRS | 3-deoxy-7-phosphoheptulonate synthase [EC:2.5.1.54] | 0 | 0 | 0 | 14403.6 | 13238 | 14903.9 | 0.000009 | 28.74 | 0.00445 | 10.438 | aroF, aroG, aroH |
| SoxRS | 3-deoxy-7-phosphoheptulonate synthase [EC:2.5.1.54] | 0 | 0 | 0 | 17582.7 | 14710.8 | 19046.6 | 0.000177 | 13.44 | 0.015956 | 10.709 | aroF, aroG, aroH |
| SoxRS | glucose-6-phosphate 1-dehydrogenase [EC:1.1.1.49 1.1.1.363] | 51293.8 | 46928.1 | 52416.4 | 78230.3 | 79974.1 | 72145.8 | 0.000792 | 9.15 | 0.027436 | 0.613 | G6PD, zwf |
| SoxRS | outer membrane protein | 19083.4 | 14650.7 | 13670.4 | 36925.3 | 41676.9 | 35414.5 | 0.000911 | 8.823 | 0.028439 | 1.266 | ompW |
| SoxRS | superoxide dismutase, Fe-Mn family [EC:1.15.1.1] | 15653.4 | 19203.1 | 21666.9 | 52822.2 | 49184 | 40963.1 | 0.00182 | 7.355 | 0.039025 | 1.339 | SOD2 |

| | | | | | | | | | | | | |
|---|---|---|---|---|---|---|---|---|---|---|---|---|
| OxyR | peroxiredoxin (alkyl hydroperoxide reductase subunit C) [EC:1.11.1.15] | 26235.7 | 31658.5 | 28124.1 | 5345.89 | 8827.95 | 6261.78 | 0.000326 | 11.5 | 0.019519 | -2.074 | PRDX2_4, ahpC |
| OxyR | peroxiredoxin (alkyl hydroperoxide reductase subunit C) [EC:1.11.1.15] | 13807.1 | 17070.5 | 14865.3 | 42280.6 | 44892.6 | 37869.3 | 0.000308 | 11.68 | 0.019519 | 1.451 | PRDX2_4, ahpC |
| OxyR | peroxiredoxin (alkyl hydroperoxide reductase subunit C) [EC:1.11.1.15] | 14682.3 | 7780.54 | 17077.3 | 34160.1 | 33913.2 | 36366.2 | 0.001712 | 7.475 | 0.037582 | 1.401 | PRDX2_4, ahpC |
| OxyR | Fe-S cluster assembly protein SufB | 6137.41 | 11510.1 | 2895.44 | 25735 | 25277 | 28395.7 | 0.001888 | 7.284 | 0.039486 | 1.951 | sufB |
| OxyR | Fe-S cluster assembly protein SufD | 0 | 0 | 0 | 11613.3 | 19456.9 | 19704.3 | 0.003114 | 6.371 | 0.048771 | 10.693 | sufD |
| OxyR | type VI secretion system secreted protein Hcp | 59093.2 | 61728.9 | 52071.8 | 32891.5 | 37559.6 | 28356.1 | 0.003244 | 6.3 | 0.049468 | -0.807 | hcp |
| NorR | nitric oxide dioxygenase [EC:1.14.12.17] | 10041.9 | 7165.11 | 8532.84 | 46180.6 | 36744.4 | 42583.7 | 0.000318 | 11.58 | 0.019519 | 2.286 | hmp, YHB1 |
| NorR | nitric oxide dioxygenase [EC:1.14.12.17] | 23041.4 | 17806.5 | 5607.52 | 123041 | 106145 | 98425.9 | 0.000463 | 10.51 | 0.022614 | 2.818 | hmp, YHB1 |
| NorR | nitric oxide dioxygenase [EC:1.14.12.17] | 16368.4 | 11965.7 | 13527.5 | 103693 | 106942 | 81653.4 | 0.000488 | 10.37 | 0.022638 | 2.804 | hmp, YHB1 |
| NorR | nitric oxide reductase subunit B [EC:1.7.2.5] | 7643.01 | 5734.95 | 3530.61 | 23989.7 | 22870.4 | 19592 | 0.000744 | 9.301 | 0.027228 | 1.975 | norB |
| NorR | nitric oxide dioxygenase [EC:1.14.12.17] | 3917.69 | 18050.2 | 9216.51 | 111616 | 101066 | 76993.2 | 0.001456 | 7.802 | 0.035162 | 3.216 | hmp, YHB1 |
| NsrR | large subunit ribosomal protein L35 | 4680.96 | 3864.32 | 2372.82 | 21942.4 | 22555.9 | 21600.4 | 0.000015 | 25.15 | 0.004555 | 2.598 | RP-L35, MRPL35, rpmI |
| NsrR | pyruvate dehydrogenase E1 component [EC:1.2.4.1] | 0 | 0 | 0 | 13523.1 | 11544.3 | 14622.2 | 0.000125 | 14.69 | 0.013646 | 10.338 | aceE |
| NsrR | nitric oxide dioxygenase [EC:1.14.12.17] | 10041.9 | 7165.11 | 8532.84 | 46180.6 | 36744.4 | 42583.7 | 0.000318 | 11.58 | 0.019519 | 2.286 | hmp, YHB1 |
| NsrR | nitric oxide dioxygenase [EC:1.14.12.17] | 23041.4 | 17806.5 | 5607.52 | 123041 | 106145 | 98425.9 | 0.000463 | 10.51 | 0.022614 | 2.818 | hmp, YHB1 |
| NsrR | nitric oxide dioxygenase [EC:1.14.12.17] | 16368.4 | 11965.7 | 13527.5 | 103693 | 106942 | 81653.4 | 0.000488 | 10.37 | 0.022638 | 2.804 | hmp, YHB1 |
| NsrR | nitric oxide dioxygenase [EC:1.14.12.17] | 3917.69 | 18050.2 | 9216.51 | 111616 | 101066 | 76993.2 | 0.001456 | 7.802 | 0.035162 | 3.216 | hmp, YHB1 |
| NsrR | pyruvate dehydrogenase E1 component [EC:1.2.4.1] | 66745.4 | 80522 | 68765.8 | 33193 | 39480.4 | 35733.5 | 0.001541 | 7.687 | 0.036253 | -0.995 | aceE |
| NsrR | pyruvate dehydrogenase E1 component [EC:1.2.4.1] | 28571.7 | 39091.4 | 29625.5 | 59263.4 | 64831.8 | 59170.3 | 0.001711 | 7.477 | 0.037582 | 0.914 | aceE |
| NsrR | Fe-S cluster assembly protein SufB | 6137.41 | 11510.1 | 2895.44 | 25735 | 25277 | 28395.7 | 0.001888 | 7.284 | 0.039486 | 1.951 | sufB |

| | | | | | | | | | | | | |
|---|---|---|---|---|---|---|---|---|---|---|---|---|
| NsrR | monothiol glutaredoxin | 2639.77 | 4299.22 | 6517.72 | 20871 | 15259.2 | 17734.3 | 0.002414 | 6.822 | 0.043934 | 2.001 | grxD, GLRX5 |
| NsrR | Fe-S cluster assembly protein SufD | 0 | 0 | 0 | 11613.3 | 19456.9 | 19704.3 | 0.003114 | 6.371 | 0.048771 | 10.693 | sufD |
| NsrR | type VI secretion system secreted protein Hcp | 59093.2 | 61728.9 | 52071.8 | 32891.5 | 37559.6 | 28356.1 | 0.003244 | 6.3 | 0.049468 | -0.807 | hcp |

#### S2.7. Inhibition of TCA cycle and promotion of amino acid and fatty acid synthesis by oxygen perturbations compared to constant aeration

The microbial system, as a consortium of different species, has inherent complexity that leads to differences in how various types of microbes regulate the orthologous protein. Considering the entire microbial system alone may obscure the subtle response mechanisms of certain types of microbes to different ventilation conditions, especially when microbes with low enzyme expression abundance exhibit different regulatory mechanisms from those with high enzyme expression abundance. Therefore, 4850 proteins distinguished by microbial origin were used for analyzing the response mechanisms of microbes to changes in oxygen environments. Table S10 shows information on proteins with significantly different abundances under continuous and intermittent oxygen perturbations compared to constant aeration, respectively. The more detailed specific enzyme information can be found in Table S11 and S12.

**Table S10.** Enzymes with significant abundance differences in individual activated sludge microorganisms after 48 hours of oxygen perturbation treatment compared to constant aeration treatment (*P*<0.05).

| **TCA cycle** | | | | |
|---|---|---|---|---|
| Function | Enzyme | Gene | $Log_2FC$ (CP vs. CA) | $Log_2FC$ (IP vs. CA) |
| provides metabolic intermediates and NADH[17] | malate dehydrogenase | *mdh* | -2.17 | -2.18, -0.91, -0.54, and 0.27 |
| | isocitrate dehydrogenase | *icd* | -0.94, and 0.23 | -3.16, and -0.45 |
| | 2-oxoglutarate dehydrogenase | *sucB* | -0.18 | |
| | | *sucA* | | -1.53 |
| **Nitrogen fixation metabolism** | | | | |
| Function | Enzyme | Gene | $Log_2FC$ (CP vs. CA) | $Log_2FC$ (IP vs. CA) |
| regulates intracellular glutamate and glutamine levels[18] | glutamate dehydrogenase | *gdhA* | | -1.34, -1.09, -0.48, and 2.88 |
| | glutamine synthetase | *glnA* | -0.40 | -1.05, and -0.14 |
| | glutamate synthase | *gltB* | | -5.97, -3.46, and -2.60 |
| **Amino acid metabolism** | | | | |
| Function | Enzyme | Gene | $Log_2FC$ (CP vs. CA) | $Log_2FC$ (IP vs. CA) |
| branched-chain amino acid | acetolactate synthase | *ilvB* | 0.71 | 2.20, 1.38, and -1.82 |
| | ketol-acid reductoisomerase | *ilvC* | 0.35 | 0.46, and -0.25 |

| | | | | |
|---|---|---|---|---|
| biosynthesis[19] | branched-chain amino acid aminotransferase | *ilvE* | | 1.07 |
| glycine biosynthesis[20] | glycine hydroxymethyltransferase | *glyA* | 0.47 | |
| phenylalanine and tyrosine biosynthesis[21] | chorismate mutase / prephenate dehydratase | *pheA* | | 3.30 |
| arginine biosynthesis[22] | argininosuccinate synthase | *argG* | | 1.79 |
| tryptophan biosynthesis[23] | 3-deoxy-7-phosphoheptulonate synthase | *aroF* | | 1.70, 0.81, and 0.46 |
| threonine biosynthesis[24] | threonine synthase | *thrC* | | 3.11 |
| alanine biosynthesis[25] | alanine-synthesizing transaminase | *alaA* | | 1.46, and -1.21 |
| alanine and serine degradation[26] | alanine-glyoxylate transaminase/serine-glyoxylate transaminase/serine-pyruvate transaminase | *AGXT* | -0.34 | -0.79 |
| **Fatty acid metabolism** | | | | |
| Function | Enzyme | Gene | $Log_2FC$ (CP vs. CA) | $Log_2FC$ (IP vs. CA) |
| fatty acid biosynthesis[27] | acetyl-CoA carboxylase | *accA* | 0.69 | 2.26 and -0.72 |
| | enoyl-[acyl-carrier protein] reductase | *fabI* | 1.11 | |
| | | *accC* | | 1.43 |
| fatty acid chain elongation cycle[28] | 3-oxoacyl-[acyl-carrier protein] reductase | *fabG* | 0.96 | |
| | 3-oxoacyl-[acyl-carrier-protein] synthase | *fabF* | | 0.83 |
| unsaturated fatty acid synthesis[29] | The 3-oxoacyl-[acyl-carrier-protein]_synthase | *fabB* | -2.86 | |
| degradation of long-chain fatty acids[30] | the long-chain acyl-CoA synthetase | *fadD* | | -3.37 |
| degradation of short-chain fatty acids[31] | The 3-hydroxybutyryl-CoA dehydrogenase | *fadB* | | 0.96 |

**Table S11.** Details of enzymes with significant differential abundance related to amino acid and fatty acid metabolism in activated sludge microbial systems after 48 hours of continuous perturbation (CP) compared to constant aeration (CA) treatment. The numbers following the aeration conditions in the table represent biological replicates.

| Enzyme | CA_1 | CA_2 | CA_3 | CP_1 | CP_2 | CP_3 | $Log_2FC$ [CPvs.CA] | -log10(p) | Gene name |
|---|---|---|---|---|---|---|---|---|---|
| 4-hydroxy-2-oxoheptanedioate aldolase [EC:4.1.2.52] | 0 | 0 | 0 | 12913.6 | 7626.76 | 10850.3 | 3.6249 | 2.4762 | hpaI, hpcH |
| adenosylhomocysteinase [EC:3.3.1.1] | 848.791 | 1556.41 | 597.886 | 2051.5 | 3520.25 | 2288.75 | 1.3882 | 1.4023 | ahcY |
| butyryl-CoA dehydrogenase [EC:1.3.8.1] | 5533.96 | 7362.92 | 5976.39 | 12069.7 | 13048.4 | 17300.1 | 1.1683 | 2.0065 | ACADS, bcd |
| enoyl-[acyl-carrier protein] reductase I [EC:1.3.1.9 1.3.1.10] | 7231.31 | 5550.41 | 1841.48 | 9678.36 | 10927.4 | 10983.9 | 1.1112 | 1.5767 | fabI |
| glucose-6-phosphate 1-dehydrogenase [EC:1.1.1.49 1.1.1.363] | 10031.5 | 7902.43 | 3737.2 | 14668.5 | 15793.3 | 12072.7 | 0.97286 | 1.496 | G6PD, zwf |
| 3-oxoacyl-[acyl-carrier protein] reductase [EC:1.1.1.100] | 2379.88 | 1647.79 | 2468.02 | 4937.34 | 4040.53 | 3674.42 | 0.96184 | 1.9644 | fabG |
| acetyl-CoA C-acetyltransferase [EC:2.3.1.9] | 12398.5 | 12094.7 | 10703.3 | 25173 | 16946.6 | 25276.7 | 0.93724 | 1.727 | atoB |
| leucyl aminopeptidase [EC:3.4.11.1] | 11352.4 | 6293.3 | 8855.26 | 16105.7 | 14344.7 | 14928.8 | 0.77599 | 1.8143 | CARP, pepA |
| lysyl-tRNA synthetase, class II [EC:6.1.1.6] | 11563.8 | 10698.8 | 5288.07 | 17300 | 13766.3 | 15729.8 | 0.7643 | 1.3544 | KARS, lysS |
| acetolactate synthase I/II/III large subunit [EC:2.2.1.6] | 6988.05 | 7472.31 | 3458.66 | 9942.17 | 9408.8 | 9889.81 | 0.70649 | 1.3801 | L, ilvB, ilvG, ilvI |
| acetyl-CoA carboxylase carboxyl transferase subunit alpha [EC:6.4.1.2 2.1.3.15] | 19201 | 35261.9 | 32221.7 | 50389.3 | 42679.3 | 47033.3 | 0.69263 | 1.5208 | accA |
| butyryl-CoA dehydrogenase [EC:1.3.8.1] | 15826.1 | 14049.7 | 16513.6 | 20696 | 26529.8 | 24712.3 | 0.63296 | 1.981 | ACADS, bcd |
| adenosylhomocysteinase [EC:3.3.1.1] | 20997.3 | 20055.5 | 17587.5 | 31776 | 29071.3 | 28155.6 | 0.60196 | 2.6122 | ahcY |
| glycine hydroxymethyltransferase [EC:2.1.2.1] | 5492.11 | 5340.39 | 5933.17 | 6861.59 | 8226.34 | 8150.49 | 0.471 | 1.9728 | glyA, SHMT |
| ketol-acid reductoisomerase [EC:1.1.1.86] | 23380.6 | 20583.3 | 26641 | 30995.2 | 31717.5 | 27201.4 | 0.34878 | 1.3428 | ilvC |

| | | | | | | | | | |
|---|---|---|---|---|---|---|---|---|---|
| imidazoleglycerol-phosphate dehydratase [EC:4.2.1.19] | 32684.8 | 35447.5 | 33970.7 | 37114.2 | 44587.5 | 47492.8 | 0.33952 | 1.3241 | hisB |
| glucose-6-phosphate 1-dehydrogenase [EC:1.1.1.49 1.1.1.363] | 108453 | 110479 | 113371 | 139356 | 131864 | 136396 | 0.29471 | 3.1886 | G6PD, zwf |
| isocitrate dehydrogenase [EC:1.1.1.42] | 22022.9 | 20817.4 | 24710.1 | 27070.8 | 25427.1 | 26844.3 | 0.23212 | 1.449 | IDH1, IDH2, icd |
| 2-oxoglutarate dehydrogenase E2 component (dihydrolipoamide succinyltransferase) [EC:2.3.1.61] | 57481.4 | 56151.8 | 57912.9 | 47966.1 | 49659.2 | 54195.9 | -0.17622 | 1.5642 | DLST, sucB |
| alanine-glyoxylate transaminase / serine-glyoxylate transaminase / serine-pyruvate transaminase [EC:2.6.1.44 2.6.1.45 2.6.1.51] | 199027 | 197284 | 172456 | 158705 | 154640 | 136512 | -0.33837 | 1.6489 | AGXT |
| glutamine synthetase [EC:6.3.1.2] | 159374 | 192975 | 188230 | 121012 | 157089 | 131856 | -0.39903 | 1.3581 | glnA, GLUL |
| alanine-glyoxylate transaminase / serine-glyoxylate transaminase / serine-pyruvate transaminase [EC:2.6.1.44 2.6.1.45 2.6.1.51] | 248134 | 232110 | 269752 | 205408 | 171489 | 143751 | -0.52657 | 1.6651 | AGXT |
| S-adenosylmethionine synthetase [EC:2.5.1.6] | 13000.1 | 16403.6 | 11724.4 | 5406.26 | 8029.64 | 8856.82 | -0.88355 | 1.645 | metK |
| 3-phenylpropionate/trans-cinnamate dioxygenase ferredoxin reductase component [EC:1.18.1.3] | 9576.3 | 10180 | 7187.33 | 6137.83 | 1960.11 | 5420.26 | -0.99504 | 1.3253 | hcaD |
| phenylacetaldehyde dehydrogenase [EC:1.2.1.39] | 2983.91 | 2496.63 | 3373.87 | 743.15 | 0 | 2000.45 | -1.6443 | 1.5078 | feaB |
| isocitrate dehydrogenase [EC:1.1.1.42] | 2255.78 | 2716.02 | 1535.13 | 1213.74 | 0 | 0 | -1.9371 | 1.5621 | IDH1, IDH2, icd |
| methionyl-tRNA synthetase [EC:6.1.1.10] | 3223.59 | 3148.84 | 1575.55 | 1288.54 | 190.482 | 525.82 | -1.9871 | 1.4641 | MARS, metG |
| malate dehydrogenase [EC:1.1.1.37] | 1387.56 | 1701.76 | 1437.88 | 0 | 524.162 | 421.213 | -2.1684 | 2.6295 | mdh |
| tyrosyl-tRNA synthetase [EC:6.1.1.1] | 4426.19 | 3468.37 | 3654.92 | 0 | 0 | 1615.08 | -2.5241 | 2.3341 | YARS, tyrS |
| thioredoxin reductase (NADPH) [EC:1.8.1.9] | 7892.73 | 5028.8 | 5530.52 | 0 | 0 | 0 | -2.6313 | 2.3684 | trxB, TRR |
| 3-oxoacyl-[acyl-carrier-protein] synthase I [EC:2.3.1.41] | 1883.49 | 3655.58 | 4635.74 | 0 | 999.26 | 0 | -2.8626 | 1.5834 | fabB |
| dihydroxy-acid dehydratase [EC:4.2.1.9] | 4688.84 | 6265.19 | 2734.47 | 0 | 0 | 0 | -3.6352 | 1.8312 | ilvD |

**Table S12.** Details of enzymes with significant differential abundance related to amino acid and fatty acid metabolism in activated sludge microbial systems after 48 hours of intermittent perturbation (IP) compared to constant aeration (CA) treatment. The numbers following the aeration conditions in the table represent biological replicates.

| Enzyme | CA_1 | CA_2 | CA_3 | IP_1 | IP_2 | IP_3 | $Log_2FC$ [IPvs.CA] | -log10(p) | Gene name |
|---|---|---|---|---|---|---|---|---|---|
| 3-hydroxyacyl-CoA dehydrogenase / 3-hydroxy-2-methylbutyryl-CoA dehydrogenase [EC:1.1.1.35 1.1.1.178] | 0 | 0 | 0 | 8306.69 | 13747.2 | 14783.2 | 3.733 | 2.3167 | HSD17B10 |
| chorismate mutase / prephenate dehydratase [EC:5.4.99.5 4.2.1.51] | 0 | 0 | 348.696 | 1331.22 | 1762.44 | 1711.2 | 3.299 | 3.0236 | pheA |
| 4-hydroxy-2-oxoheptanedioate aldolase [EC:4.1.2.52] | 0 | 0 | 0 | 7017.29 | 7476.08 | 8344.06 | 3.1659 | 4.1923 | hpaI, hpcH |
| threonine synthase [EC:4.2.3.1] | 2252.61 | 0 | 0 | 8315.84 | 5689.42 | 12378.1 | 3.1079 | 1.7224 | thrC |
| glutamate dehydrogenase (NAD(P)+) [EC:1.4.1.3] | 0 | 0 | 0 | 6126.26 | 10979.4 | 10039.4 | 2.8846 | 2.2051 | GLUD1_2, gdhA |
| 3-oxoacyl-[acyl-carrier protein] reductase [EC:1.1.1.100] | 0 | 0 | 0 | 4152.69 | 3912.16 | 8556.58 | 2.824 | 1.4627 | fabG |
| 3-hydroxyacyl-CoA dehydrogenase [EC:1.1.1.35] | 0 | 5232.01 | 0 | 13420.1 | 10292.9 | 16693.4 | 2.7451 | 2.0227 | fadN |
| acetyl-CoA carboxylase, biotin carboxylase subunit [EC:6.4.1.2 6.3.4.14] | 3292.86 | 3880.35 | 0 | 15073.5 | 12416.3 | 7699.38 | 2.2622 | 1.7175 | accC |
| acetolactate synthase I/II/III large subunit [EC:2.2.1.6] | 481.255 | 0 | 0 | 1003.07 | 1029.87 | 1058.09 | 2.1978 | 2.4712 | L, ilvB, ilvG, ilvI |
| imidazoleglycerol-phosphate dehydratase [EC:4.2.1.19] | 0 | 2366.47 | 0 | 4259.37 | 3775.33 | 5131.08 | 1.9906 | 1.9342 | hisB |
| argininosuccinate synthase [EC:6.3.4.5] | 0 | 2174.52 | 1475.09 | 5456.64 | 3633.07 | 4531.73 | 1.7879 | 1.8784 | argG, ASS1 |
| alcohol dehydrogenase [EC:1.1.1.1] | 4330.74 | 9424.09 | 11476.9 | 32008.7 | 25258.7 | 29408.6 | 1.7804 | 2.6769 | adh |
| 3-deoxy-7-phosphoheptulonate synthase [EC:2.5.1.54] | 1681.3 | 2882.63 | 0 | 6950.04 | 3847.31 | 4661.71 | 1.703 | 1.374 | aroF, aroG, aroH |
| glycerate dehydrogenase [EC:1.1.1.29] | 11613 | 7881.74 | 5939.01 | 26555.8 | 28996.7 | 22526 | 1.6182 | 2.6531 | hprA |

| | | | | | | | | | |
|---|---|---|---|---|---|---|---|---|---|
| alanine-synthesizing transaminase [EC:2.6.1.66 2.6.1.2] | 2656.91 | 3424.3 | 1155.28 | 4642 | 8158.3 | 7149.68 | 1.463 | 1.5716 | alaA |
| butyryl-CoA dehydrogenase [EC:1.3.8.1] | 5533.96 | 7362.92 | 5976.39 | 15852.1 | 17810.7 | 18006.6 | 1.453 | 3.6148 | ACADS, bcd |
| 6-phosphogluconate dehydrogenase [EC:1.1.1.44 1.1.1.343] | 4440.37 | 6412.9 | 3759.69 | 16209.7 | 13378.6 | 9813.99 | 1.431 | 1.8291 | PGD, gnd, gntZ |
| enoyl-[acyl-carrier protein] reductase I [EC:1.3.1.9 1.3.1.10] | 385.124 | 802.051 | 419.343 | 1497.05 | 1295.58 | 1523.1 | 1.4257 | 2.3979 | fabI |
| methylmalonyl-CoA mutase [EC:5.4.99.2] | 3361.18 | 2949.56 | 0 | 7273.43 | 6045.21 | 4897.96 | 1.4004 | 1.5725 | MUT |
| acetolactate synthase I/II/III large subunit [EC:2.2.1.6] | 6988.05 | 7472.31 | 3458.66 | 18046.9 | 16826.9 | 11610.4 | 1.3752 | 1.8145 | L, ilvB, ilvG, ilvI |
| enoyl-[acyl-carrier protein] reductase I [EC:1.3.1.9 1.3.1.10] | 7231.31 | 5550.41 | 1841.48 | 15298.9 | 9195.1 | 11550.3 | 1.3015 | 1.3953 | fabI |
| phenylalanyl-tRNA synthetase alpha chain [EC:6.1.1.20] | 2573.11 | 333.864 | 3227.48 | 4708.45 | 4993.99 | 5043.59 | 1.2653 | 1.5048 | FARSA, pheS |
| 2-isopropylmalate synthase [EC:2.3.3.13] | 26486.3 | 22192.8 | 13453.3 | 42563.2 | 54474.4 | 40099.5 | 1.1422 | 1.8851 | leuA, IMS |
| branched-chain amino acid aminotransferase [EC:2.6.1.42] | 13188.7 | 10230.5 | 9286.29 | 23283.6 | 26227.7 | 19317.2 | 1.0735 | 2.1819 | ilvE |
| glucose-6-phosphate 1-dehydrogenase [EC:1.1.1.49 1.1.1.363] | 10031.5 | 7902.43 | 3737.2 | 14862.8 | 17778.4 | 12899.2 | 1.0714 | 1.5705 | G6PD, zwf |
| 3-hydroxybutyryl-CoA dehydrogenase [EC:1.1.1.157] | 11293 | 4997.09 | 14354.3 | 23138.4 | 16709 | 19660.7 | 0.95746 | 1.3535 | paaH, hbd, fadB, mmgB |
| 3-oxoacyl-[acyl-carrier-protein] synthase II [EC:2.3.1.179] | 5496.28 | 3129.55 | 8203.64 | 10748.9 | 9649.82 | 9598.71 | 0.83385 | 1.3558 | fabF |
| 3-deoxy-7-phosphoheptulonate synthase [EC:2.5.1.54] | 9812.91 | 7571.19 | 11852.5 | 17582.7 | 14710.8 | 19046.6 | 0.81231 | 1.8463 | aroF, aroG, aroH |
| acetyl-CoA C-acetyltransferase [EC:2.3.1.9] | 8688.33 | 6432.96 | 10837.7 | 14649.8 | 13418.2 | 15457.2 | 0.74562 | 1.8541 | atoB |
| adenosylhomocysteinase [EC:3.3.1.1] | 20997.3 | 20055.5 | 17587.5 | 23852.8 | 36598.4 | 30022.1 | 0.6256 | 1.3023 | ahcY |
| butyryl-CoA dehydrogenase [EC:1.3.8.1] | 15826.1 | 14049.7 | 16513.6 | 26036.6 | 22574.2 | 22838.2 | 0.62312 | 2.4802 | ACADS, bcd |
| acetyl-CoA C-acetyltransferase [EC:2.3.1.9] | 18677.5 | 11109.9 | 14679.4 | 21922.8 | 20740.6 | 22719 | 0.55617 | 1.4346 | atoB |
| aspartyl-tRNA synthetase [EC:6.1.1.12] | 60793.9 | 55097 | 49922.2 | 74117.4 | 79500 | 89094.8 | 0.54969 | 2.0492 | aspS |
| 3-deoxy-7-phosphoheptulonate synthase [EC:2.5.1.54] | 8295.19 | 12709.4 | 9867.46 | 14403.6 | 13238 | 14903.9 | 0.46271 | 1.3177 | aroF, aroG, aroH |

| | | | | | | | | | |
|---|---|---|---|---|---|---|---|---|---|
| ketol-acid reductoisomerase [EC:1.1.1.86] | 23380.6 | 20583.3 | 26641 | 29712.1 | 33969.3 | 33225.6 | 0.45683 | 1.7949 | ilvC |
| glycine hydroxymethyltransferase [EC:2.1.2.1] | 5492.11 | 5340.39 | 5933.17 | 7183.81 | 7347.88 | 8095.02 | 0.43252 | 2.3799 | glyA, SHMT |
| malate dehydrogenase [EC:1.1.1.37] | 37943.8 | 36981.2 | 36199.4 | 46367.1 | 47719.2 | 40179.5 | 0.27292 | 1.5025 | mdh |
| glutamine synthetase [EC:6.3.1.2] | 185741 | 180374 | 191343 | 161003 | 177790 | 167716 | -0.13828 | 1.3641 | glnA, GLUL |
| ketol-acid reductoisomerase [EC:1.1.1.86] | 125124 | 142801 | 129391 | 110963 | 117783 | 105174 | -0.25078 | 1.5136 | ilvC |
| isocitrate dehydrogenase [EC:1.1.1.42] | 37012.8 | 33880 | 28760.2 | 25033.8 | 26399.4 | 21365.1 | -0.45301 | 1.4645 | IDH1, IDH2, icd |
| glutamate dehydrogenase (NADP+) [EC:1.4.1.4] | 58198.7 | 75486.6 | 63043.3 | 40762.4 | 54516.5 | 45662.8 | -0.48111 | 1.3315 | gdhA |
| glucose-6-phosphate 1-dehydrogenase [EC:1.1.1.49 1.1.1.363] | 108453 | 110479 | 113371 | 78230.3 | 79974.1 | 72145.8 | -0.52867 | 3.5967 | G6PD, zwf |
| malate dehydrogenase [EC:1.1.1.37] | 18062.7 | 15074.4 | 15311.1 | 9922.56 | 11288.5 | 12072 | -0.54165 | 1.9353 | mdh |
| cysteine desulfurase / selenocysteine lyase [EC:2.8.1.7 4.4.1.16] | 8528.62 | 7283.51 | 7572.54 | 4305.38 | 5095.72 | 5616.77 | -0.63888 | 2.1881 | sufS |
| 3-hydroxyacyl-[acyl-carrier-protein] dehydratase [EC:4.2.1.59] | 64477.3 | 68427.2 | 54310.7 | 45907.3 | 32354.8 | 40099.1 | -0.6615 | 1.7887 | fabZ |
| ATP phosphoribosyltransferase regulatory subunit | 9096.36 | 8275.54 | 9926.73 | 4778.04 | 5349.03 | 7085.67 | -0.66535 | 1.79 | hisZ |
| glutamate N-acetyltransferase / amino-acid N-acetyltransferase [EC:2.3.1.35 2.3.1.1] | 114304 | 110221 | 96326.6 | 62214.8 | 80500.9 | 55512.1 | -0.69475 | 1.9402 | argJ |
| acetolactate synthase I/III small subunit [EC:2.2.1.6] | 32515.9 | 31757.4 | 24561.6 | 18743.8 | 18643.8 | 16863.7 | -0.71147 | 1.9395 | S, ilvH, ilvN |
| acetyl-CoA carboxylase, biotin carboxylase subunit [EC:6.4.1.2 6.3.4.14] | 26317.7 | 33943.6 | 28609.1 | 17020.1 | 18090.6 | 18730 | -0.72301 | 2.1399 | accC |
| glucose-6-phosphate 1-dehydrogenase [EC:1.1.1.49 1.1.1.363] | 50566.3 | 58194.6 | 48367.3 | 31676.6 | 32302.5 | 30339.6 | -0.73633 | 2.6362 | G6PD, zwf |
| alanine-glyoxylate transaminase / serine-glyoxylate transaminase / serine-pyruvate transaminase [EC:2.6.1.44 2.6.1.45 2.6.1.51] | 199027 | 197284 | 172456 | 82683.9 | 124272 | 122304 | -0.78861 | 2.1189 | AGXT |
| glutamate synthase (NADPH) small chain [EC:1.4.1.13] | 5906.87 | 6776.69 | 8167.04 | 3804.87 | 3030.79 | 4785.55 | -0.84333 | 1.6815 | gltD |

| | | | | | | | | | |
|---|---|---|---|---|---|---|---|---|---|
| malate dehydrogenase [EC:1.1.1.37] | 37732.6 | 34989.2 | 32023.8 | 15988.8 | 22419 | 17357.6 | -0.90945 | 2.51 | mdh |
| glutamine synthetase [EC:6.3.1.2] | 159374 | 192975 | 188230 | 74472.4 | 107306 | 78995.9 | -1.0517 | 2.5013 | glnA, GLUL |
| glutamate dehydrogenase (NADP+) [EC:1.4.1.4] | 52309.9 | 82554.8 | 82786.8 | 26817.4 | 48163.6 | 27360.6 | -1.0886 | 1.4493 | gdhA |
| 2-isopropylmalate synthase [EC:2.3.3.13] | 20724.8 | 20483.6 | 22216.7 | 11203.6 | 11774.8 | 5968.8 | -1.1316 | 2.4027 | leuA, IMS |
| alanine-synthesizing transaminase [EC:2.6.1.66 2.6.1.2] | 30980.1 | 34054.3 | 35154.6 | 15951.3 | 13548.2 | 13762.6 | -1.2115 | 3.6869 | alaA |
| 5-methyltetrahydrofolate--homocysteine methyltransferase [EC:2.1.1.13] | 12520.8 | 13431.1 | 10582.9 | 0 | 6469.03 | 7582.12 | -1.2515 | 1.5421 | metH, MTR |
| glutamate dehydrogenase (NADP+) [EC:1.4.1.4] | 124066 | 238103 | 168630 | 63156.8 | 77082.3 | 70006.5 | -1.3361 | 1.4815 | gdhA |
| putrescine---pyruvate transaminase [EC:2.6.1.113] | 27277.6 | 24159.2 | 13584.1 | 6488.93 | 9705.79 | 9532.91 | -1.3376 | 1.4262 | spuC |
| 4-hydroxy-tetrahydrodipicolinate reductase [EC:1.17.1.8] | 1366.07 | 2308.04 | 1518.61 | 789.335 | 514.252 | 585.756 | -1.4586 | 1.6545 | dapB |
| 2-oxoglutarate dehydrogenase E1 component [EC:1.2.4.2] | 10129.6 | 10446.3 | 9530.12 | 4542.22 | 3636.23 | 2267.75 | -1.5271 | 3.1075 | OGDH, sucA |
| glucose-6-phosphate 1-dehydrogenase [EC:1.1.1.49 1.1.1.363] | 8268.22 | 13690.2 | 15087.4 | 3289.55 | 6591.63 | 0 | -1.8136 | 1.5143 | G6PD, zwf |
| acetolactate synthase I/II/III large subunit [EC:2.2.1.6] | 2284.3 | 1550.67 | 1352.38 | 0 | 1046.68 | 0 | -1.8238 | 1.449 | L, ilvB, ilvG, ilvI |
| malate dehydrogenase [EC:1.1.1.37] | 1387.56 | 1701.76 | 1437.88 | 308.722 | 0 | 630.168 | -2.1777 | 2.4494 | mdh |
| glutamate synthase (NADPH) large chain [EC:1.4.1.13] | 13343.6 | 11738.9 | 12230.9 | 0 | 0 | 0 | -2.604 | 4.5892 | gltB |
| glycerate 2-kinase [EC:2.7.1.165] | 2667.32 | 3415.55 | 4765.6 | 0 | 0 | 0 | -2.761 | 2.1319 | gck, gckA, GLYCTK |
| leucyl aminopeptidase [EC:3.4.11.1] | 2915.88 | 1389.1 | 1218.11 | 0 | 525.34 | 0 | -2.9087 | 1.3389 | CARP, pepA |
| homoserine dehydrogenase [EC:1.1.1.3] | 6439.98 | 9962.55 | 4387.65 | 0 | 0 | 0 | -3.0173 | 1.6931 | hom |
| isocitrate dehydrogenase [EC:1.1.1.42] | 2255.78 | 2716.02 | 1535.13 | 0 | 0 | 0 | -3.1595 | 2.3032 | IDH1, IDH2, icd |
| urocanate hydratase [EC:4.2.1.49] | 16313.6 | 6147.39 | 13389.1 | 0 | 0 | 0 | -3.2809 | 1.6226 | hutU, UROC1 |
| long-chain acyl-CoA synthetase [EC:6.2.1.3] | 683.895 | 1392.65 | 1778.56 | 0 | 267.18 | 0 | -3.3655 | 1.6171 | ACSL, fadD |

| | | | | | | | | | |
|---|---|---|---|---|---|---|---|---|---|
| (3R)-3-hydroxyacyl-CoA dehydrogenase / 3a,7a,12a-trihydroxy-5b-cholest-24-enoyl-CoA hydratase / enoyl-CoA hydratase 2 [EC:1.1.1.- 4.2.1.107 4.2.1.119] | 1825.28 | 2360.37 | 4134.73 | 0 | 0 | 0 | -3.4051 | 1.6423 | HSD17B4 |
| glutamate synthase (NADPH) large chain [EC:1.4.1.13] | 4922.56 | 9393.24 | 5542.29 | 0 | 0 | 0 | -3.4574 | 1.8989 | gltB |
| methionyl-tRNA synthetase [EC:6.1.1.10] | 3223.59 | 3148.84 | 1575.55 | 0 | 640.137 | 0 | -3.5203 | 1.8656 | MARS, metG |
| beta-ureidopropionase [EC:3.5.1.6] | 503.237 | 590.054 | 1275.77 | 0 | 0 | 0 | -4.4086 | 1.4325 | UPB1, pydC |
| glutamate synthase (NADPH) large chain [EC:1.4.1.13] | 3998.79 | 5593.22 | 3176.86 | 0 | 0 | 0 | -5.9708 | 2.3855 | gltB |

(14) Gao, Z.; Guéguen, C. Size distribution of absorbing and fluorescing DOM in Beaufort sea,

Canada Basin. *Deep Sea Res. Part Oceanogr. Res. Pap.* **2017**, *121*, 30–37. https://doi.org/10.1016/j.dsr.2016.12.014.
(15) Chen, M.; Park, M.; Kim, J.-H.; Shinn, Y. J.; Lee, Y. K.; Hur, J. Exploring pore water biogeochemical characteristics as environmental monitoring proxies for a $CO_2$ storage project in Pohang Basin, South Korea. *Mar. Pollut. Bull.* **2018**, *137*, 331–338. https://doi.org/10.1016/j.marpolbul.2018.10.036.
(16) Vuoristo, K. S.; Mars, A. E.; Sanders, J. P. M.; Eggink, G.; Weusthuis, R. A. Metabolic engineering of TCA cycle for production of chemicals. *Trends Biotechnol.* **2016**, *34* (3), 191–197. https://doi.org/10.1016/j.tibtech.2015.11.002.
(17) Schreier, H. J. Biosynthesis of glutamine and glutamate and the assimilation of ammonia. in *Bacillus subtilis* and other gram-positive bacteria; John Wiley & Sons, Ltd, 1993; pp 281–298. https://doi.org/10.1128/9781555818388.ch20.
(18) Tojo, S.; Satomura, T.; Kumamoto, K.; Hirooka, K.; Fujita, Y. Molecular mechanisms underlying the positive stringent response of the *Bacillus subtilis* ilv-leu operon, involved in the biosynthesis of branched-chain amino acids. *J. Bacteriol.* **2008**, *190* (18), 6134–6147. https://doi.org/10.1128/jb.00606-08.
(19) Shen, P.; Chao, H.; Jiang, C.; Long, Z.; Wang, C.; Wu, B. Enhancing production of L-Serine by increasing the *glyA* gene expression in *methylobacterium* Sp. MB200. *Appl. Biochem. Biotechnol.* **2010**, *160* (3), 740–750. https://doi.org/10.1007/s12010-009-8551-x.
(20) Ikeda, M.; Ozaki, A.; Katsumata, R. Phenylalanine production by metabolically engineered corynebacterium glutamicum with the *pheA* gene of *Escherichia coli*. *Appl. Microbiol. Biotechnol.* **1993**, *39* (3), 318–323. https://doi.org/10.1007/BF00192085.
(21) Schramm, T.; Lempp, M.; Beuter, D.; Sierra, S. G.; Glatter, T.; Link, H. High-throughput enrichment of temperature-sensitive argininosuccinate synthetase for two-stage citrulline production in *E. coli*. *Metab. Eng.* **2020**, *60*, 14–24. https://doi.org/10.1016/j.ymben.2020.03.004.
(22) Schoppel, K.; Trachtmann, N.; Mittermeier, F.; Sprenger, G. A.; Weuster-Botz, D. Metabolic control analysis of L-tryptophan producing *Escherichia coli* applying targeted perturbation with shikimate. *Bioprocess Biosyst. Eng.* **2021**, *44* (12), 2591–2613. https://doi.org/10.1007/s00449-021-02630-7.
(23) Zhang, Y.; Meng, Q.; Ma, H.; Liu, Y.; Cao, G.; Zhang, X.; Zheng, P.; Sun, J.; Zhang, D.; Jiang, W.; Ma, Y. Determination of key enzymes for threonine synthesis through in vitro metabolic pathway analysis. *Microb. Cell Factories* **2015**, *14* (1), 86. https://doi.org/10.1186/s12934-015-0275-8.
(24) Kim, S. H.; Schneider, B. L.; Reitzer, L. Genetics and regulation of the major enzymes of alanine synthesis in *Escherichia coli*. *J. Bacteriol.* **2010**, *192* (20), 5304–5311. https://doi.org/10.1128/jb.00738-10.
(25) Oikawa, T. Alanine, Aspartate, and Asparagine Metabolism in Microorganisms. In *amino acid biosynthesis ~ pathways, regulation and metabolic engineering*; Wendisch, V. F., Ed.; Springer: Berlin, Heidelberg, 2007; pp 273–288. https://doi.org/10.1007/7171_2006_062.
(26) Beld, J.; John Lee, D.; D. Burkart, M. Fatty acid biosynthesis revisited: structure elucidation and metabolic engineering. *Mol. Biosyst.* **2015**, *11* (1), 38–59. https://doi.org/10.1039/C4MB00443D.
(27) Lee, S.; Lee, S.; Yoon, Y. J.; Lee, J. Enhancement of long-chain fatty acid production in *Escherichia coli* by coexpressing genes, including *fabf*, involved in the elongation cycle of fatty acid biosynthesis. *Appl. Biochem. Biotechnol.* **2013**, *169* (2), 462–476. https://doi.org/10.1007/s12010-012-9987-y.
(28) Yuan, Y.; Sachdeva, M.; Leeds, J. A.; Meredith, T. C. Fatty acid biosynthesis in *Pseudomonas Aeruginosa* is initiated by the FabY class of β-ketoacyl acyl carrier protein synthases. *J. Bacteriol.* **2012**, *194* (19), 5171–5184. https://doi.org/10.1128/jb.00792-12.
(29) Mashek, D. G.; Li, L. O.; Coleman, R. A. Long-chain acyl-Coa synthetases and fatty acid channeling. *Future Lipidol.* **2007**, *2* (4), 465–476. https://doi.org/10.2217/17460875.2.4.465.
(30) DiRusso, C. C. Primary sequence of the *Escherichia coli fadBA* operon, encoding the fatty acid-oxidizing multienzyme complex, Indicates a High Degree of Homology to Eucaryotic Enzymes. *J. Bacteriol.* **1990**, *172* (11), 6459–6468. https://doi.org/10.1128/jb.172.11.6459-

6468.1990.
